# An open-sourced bioinformatic pipeline for the processing of Next-Generation Sequencing derived nucleotide reads: Identification and authentication of ancient metagenomic DNA

**DOI:** 10.1101/2020.04.20.050369

**Authors:** Thomas C. Collin, Konstantina Drosou, Jeremiah Daniel O’Riordan, Tengiz Meshveliani, Ron Pinhasi, Robin N. M. Feeney

## Abstract

Bioinformatic pipelines optimised for the processing and assessment of metagenomic ancient DNA (aDNA) are needed for studies that do not make use of high yielding DNA capture techniques. These bioinformatic pipelines are traditionally optimised for broad aDNA purposes, are contingent on selection biases and are associated with high costs. Here we present a bioinformatic pipeline optimised for the identification and assessment of ancient metagenomic DNA without the use of expensive DNA capture techniques. Our pipeline actively conserves aDNA reads, allowing the application of a bioinformatic approach by identifying the shortest reads possible for analysis (22-28bp). The time required for processing is drastically reduced through the use of a 10% segmented non-redundant sequence file (229 hours to 53). Processing speed is improved through the optimisation of BLAST parameters (53 hours to 48). Additionally, the use of multi-alignment authentication in the identification of taxa increases overall confidence of metagenomic results. DNA yields are further increased through the use of an optimal MAPQ setting (MAPQ 25) and the optimisation of the duplicate removal process using multiple sequence identifiers (a 4.35-6.88% better retention). Moreover, characteristic aDNA damage patterns are used to bioinformatically assess ancient vs. modern DNA origin throughout pipeline development. Of additional value, this pipeline uses open-source technologies, which increases its accessibility to the scientific community.

## Introduction

Optimised bioinformatic pipelines are of particular importance in the broad study of metagenomics, which consist of large datasets of multi-origins, and the associated complexities of large-scale computational processes such as comparative sequence alignment and multiple taxa identifications. The emerging field of ancient metagenomics adds to these processing complexities with the need for additional steps in the separation and authentication of ancient sequences from modern sequences. Currently, there are few pipelines available for the analysis of ancient metagenomic DNA (aDNA)^1–4^ The limited number of bioinformatic pipelines for aDNA metagenomics can be attributed to a low yield of DNA compared to that achieved in modern metagenomic DNA extraction. This results in the use of high-cost DNA capture techniques to improve aDNA yields to levels suitable for bioinformatic assessment^3–7^. In parallel, the lack of bioinformatic pipelines optimised for the processing of lower aDNA yields (i.e. those studies which do not use high-cost DNA capture techniques) deters researchers from exploring and developing alternative methods of aDNA extraction for metagenomic purposes.

Those metagenomic studies performed using DNA capture techniques, necessary for existing bioinformatic pipelines, allow for the implementation of “quick” bioinformatic comparisons using RefSeq or Blastn megablast options due to higher yields of DNA^5–7^. Due to higher yields, however, these pipelines, by their nature, often compromise between read conservation and processing time, thus losing sequences throughout individual computational steps. In doing so these pipelines do not account for the nature of metagenomic aDNA which, having originated from multiple sources (e.g. bone^8^, soil^3,9^), can vary in aDNA yields and in the degree of damage over time leading to a potential loss and underrepresentation of aDNA sequences.

The damage pattern of aDNA serves as a useful tool in the distinction between ancient and modern DNA sequences^10,11^. Characteristically, ancient sequences should consist of heavily fragmented DNA strands^6,12,13^, depurination, depyrimidination and deamination events^14,15^ and miscoding lesions^16,17^. These characteristics therefore form the central damage pattern for aDNA authentication in the development of this bioinformatic pipeline. In addition to the high cost associated with DNA capture techniques, a further issue arises from the direct selection of target taxa for probe generation prior to sequencing. This action inevitably introduces a selection bias into a study and could prevent the metagenomic analysis of all aDNA present within a sample with targeted DNA yields overshadowing untargeted yields^7,18,19^. The additional cost associated with computation and long processing times for comparative analysis acts as a barrier to more widespread use of aDNA metagenomics in fields such as archaeology, bioanthropology and paleoenvironmental sciences. Bioinformatic costs associated with computation to achieve faster processing speeds or the use of third-party interface platforms are usually unavoidable due to the large demands on processing time and the steep learning curve needed to gain proficiency in the open-source alternatives, which are often less user-friendly and lack the benefit of customer support and easily accessible manuals.

Using a method developed for the extraction of metagenomic aDNA from anthropogenic sediment (sediment that has come into contact with past human activity) without the use of DNA capture techniques^9,20,21^, we present a bioinformatic pipeline optimised for the identification and authentication of metagenomic aDNA that can be applied to studies yielding comparatively lower yields of aDNA. The development of the bioinformatic pipeline involves four fundamental underpinnings, in which it must:

- Cater to low yielding metagenomic aDNA and conserve reads wherever possible
- Be able to process data within a reasonable timeframe and allowing for multiple taxa identifications
- Be developed using open-source technologies and software wherever possible to facilitate universal access and to bypass financial barriers
- Be accompanied by a step-by-step user-friendly manual, to facilitate its use without coding and programming expertise (supplementary; SI)

## Results and Discussion

### Identifying the smallest retrievable aDNA fragment and establishing minimum sequence length threshold

The smallest length of retrievable aDNA fragments were identified through the digital visualisation of aligned DNA sequences using UGENE^22^ and Geneious R10^23^ software. To test for the smallest retrievable aDNA fragment, adapter sequences were removed using Cutadapt^24^, with a minimal sequence length threshold (cut) of 15bp. This threshold was chosen after a cut of 0 was initially used and quality analysis of sequences using FastQC software^25^ showed the absence of DNA sequences below 15bp (Figure 1). The resultant file sequences were then compared to those of the National Center for Biotechnology Information’s (NCBI) genomic nucleotide database using Basic Local Alignment Search Tool (BLAST)^26–28^. BLAST results were imported into MEtaGenome ANalyzer (MEGAN)^1^ for visualisation of genomic assignments. Mammalia and Plantae assignments at the taxonomic genus level were assigned a unique identification number. Using the random number generator specified in the SI (step 9), unique identification numbers were selected and used for further in-depth alignment to its associated reference genome using Burrows-Wheeler Aligner (BWA)^29^. Alignments were converted to SAM format^30^ and imported into DNA visualisation software. This visualisation allowed for the manual identification of (C>T, G>A) damage patterns at the terminal ends of aligned sequences against a reference genome. Fragments without these characteristics were deemed modern in origin.

**Figure 1.**
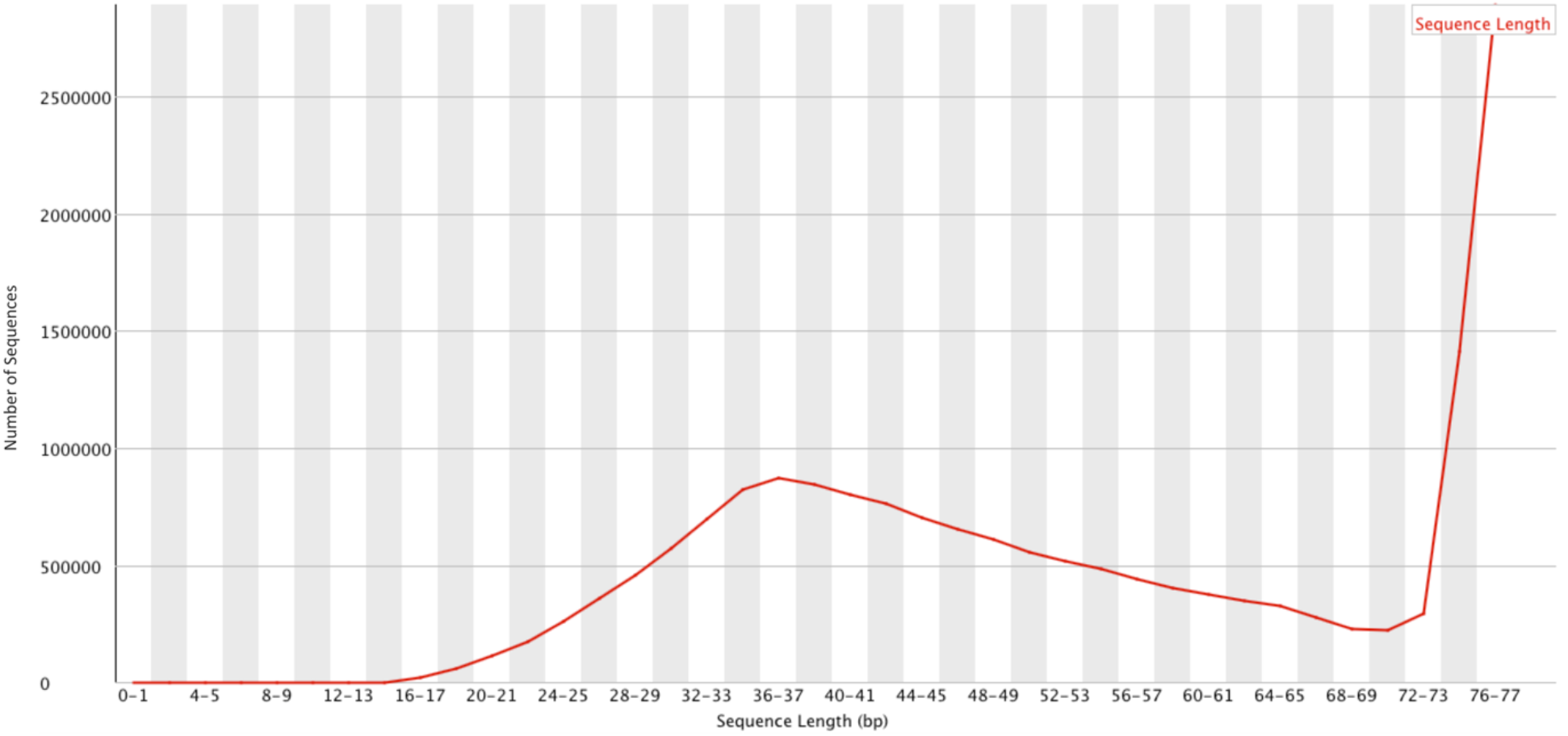
Distribution of Sequence Length After Adapter Removal Using FastQC. The number of sequences are plotted against their respective sequence lengths. The red line represents the peak number of sequences for each length. The plot shows the absence of DNA sequences below 15bp.

This procedure resulted in the identification of ancient sequences ranging from 22 - 28bp in length using the extraction method outlined by Collin^9,20^. For comparison, using Dabney’s method which was developed for aDNA extraction from bone^31^, Slon^3^ found that the lowest extractable reads were 35bp when using a ladder that mimicked aDNA. This suggests that the method used in this study is capable of extracting shorter aDNA fragments and as such may achieve a wider range of representative sequences. This is particularly important when considering the nature of DNA damage over time which not only results in increased fragmentation through oxidative strand breakages^10,11^, but also the potentially disproportionate lesion damage to genomes with high cytosine content^16,17,32^ which could otherwise be overlooked. It should be noted however that the shorter the DNA sequence, the more prone it is to misalignment errors^33^. For this reason, a cut of 28bp was selected as a conservative threshold for the purpose of this study.

### Creation of a non-redundant, 10% representative sample file for comparative analysis

One of the most common issues with the bioinformatic assessment of DNA sequences is the processing power required (corresponding to speed) and time it takes to process data. This is especially true for metagenomic data, where multiple taxa identifications are sought within a single sample. The size of a file is proportional to the time required for a process to take place. Therefore, reducing the size of a file will reduce the time required for processing.

A non-redundant sample file was created using FastX Toolkit^34^. FastX’s collapser program merges all sequence repetitions (duplicates) for a region of coding into a single representative sequence, while maintaining read count data. This is performed for all duplicates until only unique sequences remain (Figure 2A). This reduction in sequences reduces the file size considerably (1.8GB – 956MB) thus reducing associated processing time.

**Figure 2.**
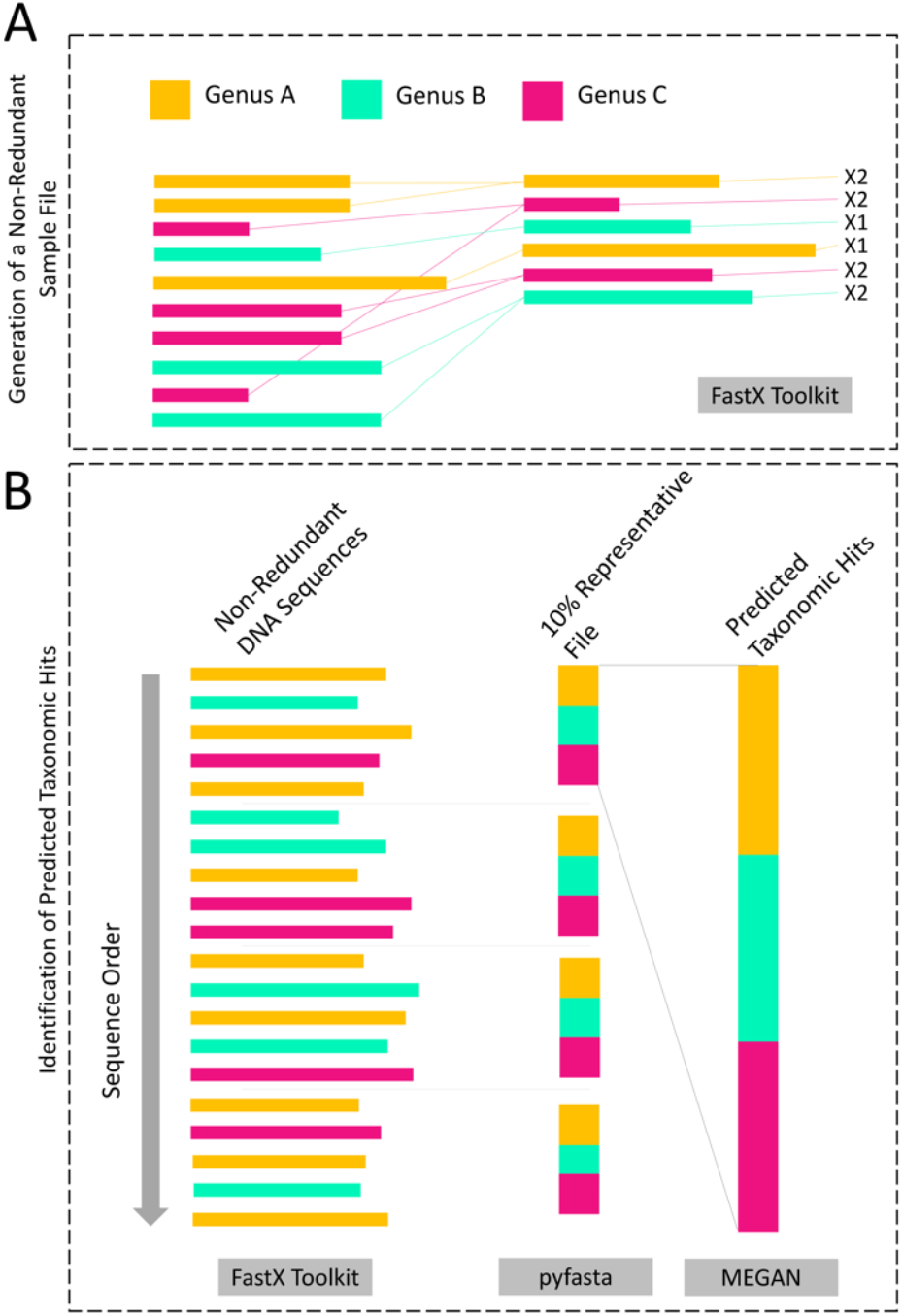
Schematic Overview for the Generation of Non-Redundant Segmented (10%) File for Comparative Analysis and Identification of Predicted Taxonomic Hits. (**A**) Generation of non-redundant file. Identically repeating sequences are merged into a single representative sequence for each occurrence in the generation of a non-redundant sample file. (**B**) Generation of the segmented (10%) file and identification of predicted taxonomic hits. Different taxa are represented by varying colours and the software used for each step is listed in the grey boxes.

This pipeline further improves the processing time of comparative analysis by splitting the non-redundant sample file into 10 representative files of equal size, each file being representative of 10% of the total sequences within an entire sample. This was achieved using pyfasta^35^. This action resulted in a further reduction of file size (956MB – 96MB per 10% file). A representative file is made possible due to two properties of DNA extraction. Firstly, multiple sequences representing the various coding regions for taxa are extracted, increasing the likelihood of representative distribution between the files. Secondly, NGS platforms sequence DNA fragments in a random order as they enter the flowcell. The corresponding sequence data is saved in this same order, meaning DNA sequences should be randomly distributed throughout the representative files (Figure 2B).

To validate that these files are representative of an entire sample’s sequences, the full sample identifications were compared to predictions made using the 10% sample files. Samples were processed using a cut of 28bp before being made non-redundant and split into 10 representative files. To eliminate the potential for bias, a randomised number generator (SI, step 9) was used to select one of the representative files for comparative alignment using BLAST. Comparative alignment was performed and resulting BLAST files imported into MEGAN using a bit-score (‘min-score’) of 40 within the top 10% of best alignments, and the default “naïve” lowest common ancestor (LCA) algorithm as described by Huson^1^. The bit-score measures the similarity of a sample sequence to a comparative sequence through complex computations^36^. Huson^1^ used a bit-score of 30 for MEGAN analysis of an ancient mammoth dataset, here we use 40 as a more conservative threshold. The resulting genomic assignments for the 10% representative file were compared to the total genomic assignments achieved from the unsplit originating non-redundant file. The mean percentage difference and standard error between expected hits based on the 10% files and actual hits achieved with the 100% file were calculated. The expected total genomic assignments predicted by the 10% file was accurate to those achieved within the 100% file within 0.007% (±1.101 SEM). The use of a 10% representative non-redundant file drastically reduced the amount of time required for comparative alignment processing: processing time for the representative file was 76.86% less than the time required for the original unaltered file (229 hours to 53

### BLAST parameters in comparative analysis for time conservation

BLAST is a multi-platform algorithm that allows users to query sample sequences against a specified database^26^. In the case of a metagenomics study, we recommend use of NCBI’s entire nucleotide database^37^ (https://www.ncbi.nlm.nih.gov/). BLAST has a series of options that allow the user to optimise the comparative analysis process for the data being queried^26–28^. The ‘task exec’ option is the most important of these, allowing users to specify the type of search required. The task blastn and task blastn-short options are best suited for inter-taxon comparisons using short sample sequences, the latter of these being optimised for sequences shorter than 50bp^38^. In the context of aDNA most authentic sequences fall within the range of 30-70bp^6,12,13^. The use of blastn-short is therefore unsuitable for aDNA analyses, and thus blastn is utilised herein.

Specification of the task blastn option automatically sets the amount of base pairs required to confirm a match between a query sequence and a reference sequence as 11bp^38^. This is referred to as word_size. The lower the word_size set, the more homologous sequences will be detected regardless of high fragmentation and DNA damage patterns. While this is beneficial for a study assessing aDNA sequences, the lower the word_size set the more processing power and time is required. This introduces a point of compromise between highest homology and shortest processing time. We used a word_size setting of 14bp, half the value of the smallest available query sequence after cut threshold (28bp) was accounted for. Using the 10% representative, non-redundant sample file, this improved processing time by 10.42% when compared to use of default settings (53 hours to 48). The resulting BLAST files were compressed to reduce file size and to allow for faster transfer into MEGAN (16 minutes compared to 27).

### Confirmation of a taxon’s presence using multi-alignment authentication

The use of a multi-alignment authentication approach to metagenomic DNA sequences reduces the likelihood of misalignment errors^3,7,21^. The first authentication is undertaken using BLAST with MEGAN for the comparative analysis and assignment of genomic sequences. This is facilitated through the use of bit-scores, measuring the similarity between a query sequence and a reference^36^. BLAST results were imported into MEGAN and parameters were set using a conservative bit-score of 40 within the top 10% of best alignments, and the default “naïve” LCA algorithm. By using the LCA algorithm, reads are assigned across a taxonomy^1^. Sequences that have a bit-score within the specified percentage of best alignments within a taxonomy are binned into the lowest possible common ancestor position. Those sequences that align to multiple taxa within a grouping are binned into a higher taxonomic level until multiple assignments are no longer occurring. To reduce the chances of false positive identifications of taxa at the family or genus level a minimum of 1% of the total assigned reads was necessary to accept a taxon as present and for use in downstream analyses^3,21^. The resulting taxa identifications are used as the main taxon labels from this point forward, and they inform the user on which reference sequences to download for in-depth sequence alignment using BWA, the second authentication.

BWA is a software package for mapping low-divergent sequences to a large reference genome^29^. Because the software uses low-divergent sequences it is considered more stringent than mass comparative alignment tools such as BLAST. BWA has a variety of modes to select from depending on the data being queried. Typically for ancient DNA either BWA aln or BWA-MEM are utilised^8,39^. While BWA-MEM is recommended for the use of sequences between 70-100bp^40^, we recommend the use of BWA aln for aDNA sequences between 30-70bp for its ability to retain more reads based on the current literature^41^ with a disabled seed length (−l 1000). Furthermore, it has been reported that the use of BWA aln over BWA-MEM conserves more single nucleotide polymorphisms (SNPs)^40,41^ useful for in-depth analysis of a taxon. Seed length refers to the amount of base pairs within a read required to match sequentially between a query and reference sequence for a match to be made^29^. While the use of seed length can considerably improve the processing time required for alignment to complete^41,42^, due to both the damaged nature of aDNA fragments and the multiple-origin sources of DNA, we recommend disabling seed length to allow for the alignment and conservation of damaged reads^43^. Upon completion of alignment a minimum threshold of 250 genomic hits are deemed necessary for a taxon to be processed downstream. This is because alignments with less than 250 reads were often found insufficient for mapDamage to plot damage patterns effectively. In some cases, MGmapper^44^ was also utilised for additional authentication of taxa using the default settings and a minimum score of 20. This can, however, add to the overall processing time of genomic fragments and was not deemed a necessary step for the confirmation of a taxon for further analysis. Of additional importance, the use of mult-alignment authentication for the identification of a taxon reduces the potential for human derived selection bias by removing the ambiguity of reference genome selection. The requirement of multiple genomic hits using multiple authentication methods increases confidence in overall metagenomic results.

### Identification of optimal mapping quality score for ancient metagenomic authentication

Mapping quality (MAPQ) refers to the degree of confidence that a sequence is correctly mapped to reference genome coordinates. In aDNA research a MAPQ of between 25-30 is typically used to extract aligned reads from poorly and non-aligned reads^21,43,45,46^. A MAPQ of 25-30 corresponds to a map accuracy of 99.68 - 99.9% (−10 log10 P), while allowing for representation of damaged sequences which typically score lower than their modern counterparts owing to deamination events at the terminal ends of sequences.

To identify the optimum mapping quality, aligned sequences were imported into DNA visualisation software^22,23^ after in-depth alignment using BWA. Visualisation of DNA sequences allowed for the manual identification of mapping qualities associated with reads possessing misincorporation events as a consequence of deamination and lesion damage^10,11,47^. The majority of sequences possessing this damage fell into the MAPQ range of 25-30. Of note, however, was the aligner’s inability to ascertain a read’s single point of origin when it fell within a repeat element of a genome. This typically resulted in a MAPQ between 20-30. The heavily fragmented nature of aDNA and the occurrence of conserved sequence regions across taxa increases the likelihood of repeating sequences along a genome in conserved regions. As such, while the use of a higher MAPQ increases confidence in results, it can also result in the loss of authentic aDNA.

To test the percentage difference in authentic ancient reads passing into subsequent steps using a MAPQ of 25 and 30, sequences were processed and taxonomic assignments were identified using MEGAN. All taxonomic assignments were given a unique identification number and using the random number generator (SI, step 9) were selected for in-depth alignment to its associated reference genome using BWA. Aligned sequences were extracted from the originating file using SAMtool’s^30^ with a MAPQ specification of either 25 or 30. Sequences were sorted by coordinate and duplicates removed before authentication of DNA damage patterns using mapDamage^48,49^. Both MAPQ scores resulted in the identification of authentic aDNA fragments with a damage pattern >10%. Using a MAPQ of 25 resulted in the greater retention of sequences (23.43%) and an increased damage profile (8.33% C>T, 8.22% G>A) of DNA sequences when compared to a MAPQ of 30. As such, 25 was selected as the optimum MAPQ.

### Conservation of mapped sequences during PCR duplicate removal

Duplicate sequences arise during the sequencing process when two or more copies of the same DNA molecule cross over onto different primer lawns within a flowcell during bridge PCR amplification. For this reason, we define a duplicate as the presence of two or more identical DNA sequences assigned to a single sample. PCR duplicates can be problematic in the assessment of authentic DNA sequences, most commonly arising in the proportional over-representation of specific areas of coding^50^. This is referred to as amplification bias or base-composition bias. To ensure the integrity of authentic DNA data, and to mitigate the potential effects of duplicate sequences, they are bioinformatically removed.

The removal of PCR duplicates from NGS-derived data often involves the use of either SAMtools or Picard Tools^30,51^. Both these methods identify duplicate sequences by the external coordinate location of outer mapped reads. In cases where two or more sequences have the exact same 5’ position coordinates, the sequence with the highest map quality score is retained and all other sequences are removed. SAMtools^30^ differs in that the same function can also be accomplished on the reverse, 3’ end of a sequence if specified. Furthermore, SAMtools is not applicable to unpaired sequences or those mapped to different chromosomes^52^. This means that a sequence with the same 5’ start coordinate as another sequence, but mapped to a different chromosome, will be marked as a duplicate and removed. Picard Tools^51^ avoids this issue by taking into account the intrachromosomal sequences. Additionally, Picard Tools takes into account soft-clipping of bases at the 5’ position of mapped reads, performing calculations to locate where the 5’ position would be if the entire sequence were mapped to the reference genome^52^. However, the use of external coordinate location as a method for duplicate removal in both commonly used methods cannot account for internal sequence variations such as SNPs, resulting in a potential loss of authentic DNA sequences. SNPs represent one of the most common types of genetic variation that can be used for detailed interpretation of a taxon^53^. The conservation of sequences increases the likelihood of retaining these SNPs. Additionally, these conserved reads play a crucial role in the statistical measurement of aDNA damage patterns, with a better assessment of deamination frequency over base position of reads^49^. This is especially true in the context of an exploratory metagenomic study without use of DNA capture techniques, where the DNA representative of a taxon is often small (1Kbp – 100Kbp) in number.

aweSAM^54^ is a SAM assembly collapser, that uses a sequence’s coordinates at the 5’ and 3’ end along with strand information as the unique insert identifiers, while keeping the sequence with the highest MAPQ. The use of multiple unique insert identifiers to locate a duplicate facilitates the conservation of reads that may be lost through other duplicate removal tools, such as SAMtools and Picard Tools. We found that use of aweSAM resulted in the conservation of between 4.35 – 6.88% of total mapped sequences when compared to SAMtools and Picard Tools. However, the increased complexity of additional unique insert identifiers used has an unfavourable effect on the processing time and the memory required, which represents a trade-off between read conservation and duration of a process. Individual users may wish to adjust this compromise based on their requirements.

### Assessment and authentication of aDNA damage patterns

DNA damage patterns were identified using mapDamage. mapDamage is a computation framework that quantifies aDNA damage patters among NGS-derived sequences^48,49^, using a statistical model based on the damage profile of aDNA fragments described by Briggs^10^.

Taxa are deemed ancient by assessing the frequency of C>T base substitutions at the 5’ terminus along with G>A base substitutions at the 3’ terminus of queried sequences. Depending on the source of DNA and the age of a context or specimen, different damage frequencies may be set as a minimum threshold to accept a sample as ancient^47^. Here we use two frequencies: a lower threshold of ≥0.05 at both terminal ends (representing 5% damage), and a higher threshold of ≥0.10 for terminal ends (representing 10% damage). As the extraction method9,20 used in this study is designed for the exploration of ancient metagenomic DNA within anthropogenic sediments without the use of DNA capture techniques, yields of authentic ancient sequences can be lower. As such, ≥0.05 can be used as the lowest threshold for a taxa to be considered potentially ancient for further study using subsequent DNA capture techniques. The reduced potential for selection biases using this exploratory method makes the lower threshold an acceptable compromise. Ultimately, if a taxa reached the higher threshold of ≥0.10 it can be considered definitively ancient in origin.

## Conclusion

The bioinformatic pipeline demonstrated here actively conserves reads by identifying the shortest DNA sequences available. This pipeline displays a substantial decrease in the amount of time required for the processing of metagenomic DNA through the generation of non-redundant 10% representative files and optimisation of BLAST parameters. Multiple-alignment authentication ensures confidence in the authenticity of taxa identifications. Furthermore, the additional conservation of aDNA sequences is achieved through the use of the optimal MAPQ setting, and the use of multiple sequence identifiers within the duplicate removal process.

This bioinformatic pipeline can be used in the exploratory assessment of metagenomic aDNA, when used in conjunction with an extraction method without the use of DNA capture techniques. The use of two damage thresholds allows for the future selection of DNA probes for subsequent in-depth metagenomic studies. The use of open-sourced bioinformatic software throughout the pipeline reduces the cost burden of many bioinformatic software packages, and thus increases the accessibility of metagenomic analyses.

## Methods

### Genomic material tested

The genomic material used for the authentication of this bioinformatic pipeline originate from anthropogenic sediments taken from two archaeological sites: Drumclay Crannóg, Co. Fermanagh, Northern Ireland and Satsurblia Cave, Imereti, Georgia.

Drumclay Crannóg is an Irish Early-Medieval to Pre-Industrial archaeological site located in county Fermanagh, Northern Ireland (54°2133.18N 7°3724.18W). Anthropogenic sediments were secured in bulk from six locations representing a primary occupation layer dating to the CE 10-11^*th*^ C; The garden (SN4818), the garden pathway to the house (SN3746), the wall/ wall packing of the house (SN4526), the northern compartment within the house (SN4537), the hearth (SN4551), and the southern ‘bed’ compartment (SN4592).

Satsurblia Cave is an Upper-Palaeolithic (29,000 BCE - 14,000 BCE) archaeological cave site located in the geographic region of the Southern Caucasus (42°2238.05N 42°363.40E). Anthropogenic sediments were secured in bulk from two areas of anthropogenic activity dating to approximately the same time-frame; one associated with tool processing (Area A; SAT17 LS30-35) and the other associated with hearth use (Area B; SAT17 LS36-40)^55^.

All anthropogenic sediments were excavated, sampled and stored in bulk, following conventional archaeological excavation techniques and standards^56,57^.

### Extraction, preparation and sequencing of genomic material

DNA extraction, library preparation and indexing steps were undertaken in an EU grade B (ISO 5) clean room under EU grade A (ISO 5) unilateral air-flow hoods at a dedicated aDNA laboratory, University College Dublin (UCD), Ireland. The laboratory surfaces were periodically cleaned with 10% sodium hypochlorite solution, DNA-OFF (MPBIO 11QD0500) and 70% ethanol, and all utensils and equipment were treated likewise after use and sterilised by UV irradiation when not in use. Tyvek suits (DuPont TYV217), hair nets (Superior bouffant01), face masks (Superior SKU83524), and nitrile gloves were used to limit contamination. PCR and subsequent steps were undertaken in an EU grade C laboratory (ISO 7) due to increased sample stability upon amplification. Extraction of DNA was performed as outlined by Dabney^31^ with optimisations described by Collin^9,20^ and libraries prepared using Meyer and Kircher^58^. PCR Amplification was performed as outlined by Gamba^13^ at a rate of 15 cycles and cleaned as specified by Collin^9,20^. Analysis of PCR reaction concentrations were performed on an Agilent 2100 Bioanalyser following the instructions of the manufacturer. Based on these concentrations, samples were pooled into a 20ng working solution. Concentration and molarity (nmol/L) of the working solution were ascertained using the Bioanalyser and a Qubit4 for fluorometric quantification following manufacturer guidelines. Sequencing was undertaken at UCD Conway Institute of Biomolecular and Biomedical Research on an Illumina NextSeq 500/550 using the high output v2 (75 cycle) reagent kit (Illumina TG-160-2005).

### Computational hardware specifications

Non-BLAST process analyses were performed in UCD using a Mac mini (late 2014) with 3GHz Intel Core i7 processor, 16GB 1600 MHz DDR3 RAM and a total storage of 1TB. BLAST processing was performed using the University of Manchester CSF3 system: BLAST-process v2.9.0 at “Skylake” facility using parallel job functionality on 16 or 32 core configurations using a Single Note Multi-core (SMP) setup with 6GB per core. Skylake facility is made up of 864 cores: 27 nodes of 2×16-core Intel Xeon Gold 6130 CPU @2.10GHz + 192GB RAM + 100Gb/s (4X EDR) mlx5 Mellanox InfiniBand. The total storage available for files in the lustre file system (known as scratch) is 692TB. CSF Nodes are based on CentOS Linux 7.4.1708.

### Identification of the smallest retrievable aDNA fragment

Adapter sequences were removed from raw sequencing files using cutadapt v1.9.124 with a minimum sequence length or “cut” threshold of 0bp and imported into FastQC^25^ for smallest sequence length available (15bp). Samples were re-cut using the smallest identified sequence length above (15bp), with a minimum overlap of 1 to reduce the over-cutting of bases that closely match an adapter sequence. Resulting sequences were comparatively analysed to a localised NCBI genomic database (2019) using BLAST^26–28^ task_blastn and a word_size of 14. BLAST results were input into MEGAN v6.2.1^31^ using a bit-score of 40 within the top 10% of best alignments, and the default “naïve” LCA algorithm. All Mammalia and Plantae assignments at the genus taxonomic level were assigned a unique identification number. Using the random number generator specified in SI (step 9), unique numbers were selected and aligned to their associated reference genome using BWA v07.5a.r405^29^ aln function with a disabled seed (−l 1000) and converted to SAM format using BWA samse function. Aligned sequences were individually imported into UGENE v1.32^22^ and Geneious R10 software for visualisation and manual identification of (C>T, G>A) misincorporation events at the terminal ends of sequences. The smallest identifiable fragment with misincorporation characteristics indicative of deamination DNA damage was deemed the smallest retrievable aDNA fragment, and thus considered the minimum sequence length threshold for subsequent applications of cutadapt.

### Validation of the non-redundant, 10% representative sample file for comparative analysis

Adapter sequences were removed from raw sequencing files using cutadapt v1.9.1^24^ with a cut of 28bp and a minimum overlap of 1. Non-redundant sample files were created using the fastx_collapser function of the Fastx toolkit v0.0.13^34^ and split into 10 (−n 10) new files of similar size using the split function of pyfasta v0.5.2^35^. Comparative alignment to a localised NCBI genomic database (2019) was undertaken using BLAST^26–28^ task_blastn and a word_size of 14. BLAST results were input into MEGAN v6.2.1^31^ using a bit-score of 40 within the top 10% of best alignments, and the default “naïve” LCA algorithm. Mean percentage difference between the genomic hits achieved from the 10% representative file and the total genomic assignments achieved from the unsplit non-redundant file generated after fastx_collapser was calculated. Standard deviation was further calculated in order to obtain the standard error of the mean.

### Identification of optimal mapping quality score

Adapter sequences were removed using cutadapt v1.9.1^24^, a cut of 28bp and minimum overlap of 1. Non-redundant sample files were created using the fastx_collapser function of Fastx toolkit v0.0.13^34^ and split into 10 (−n 10) files using the split function of pyfasta v0.5.2^35^. Sequences were comparatively analysed to a localised NCBI genomic database (2019) using BLAST^26–28^ task_blastn and a word_size of 14. BLAST results were input into MEGAN v6.2.1^31^ using a bit-score of 40 within the top 10% of best alignments, and the default “naïve” LCA algorithm. To reduce the chances of false positive identifications of taxa at the family or genus level a minimum of 1% of the total assigned reads was necessary to accept a taxon as present and use for downstream analyses^3,21^. Sequences passing this threshold were aligned to their corresponding genome using the original cut fasta file and BWA v07.5a.r405^29^ aln function with a disabled seed (−l 1000). SAI file alignments were converted to SAM format using BWA samse function. Aligned SAM files were imported into UGENE v1.32^22^ and Geneious R10^23^ software for visualisation of damaged sequences and their aligner-assigned MAPQ.

To compare MAPQ of 25 and 30, taxonomic assignments derived from MEGAN analysis were assigned a unique identifier number and randomly selected using the random number generator (SI, step 9), for BWA alignment and conversion to SAM as specified above. Mapped sequences were extracted using SAMtools v1.3.1^30^ view function and a MAPQ of 25 and 30 to create two separate comparative files. Sequences were sorted by coordinate using SAMtools sort function and duplicate sequences removed using aweSAM_collapser.sh shell script^54^. MapDamage2.0^49^ was used to quantify DNA damage through the presence of misincorporation events (C>T, G>A) at the terminal ends of the sequences. Percentage difference of total sequences identified and ancient damage patterns was calculated using the mean variation between MAPQ 25 and 30.

### Comparison of duplicate removal tools

Adapter sequences were removed using cutadapt v1.9.1^24^ with a cut of 28bp and minimum overlap of 1. Creation of mon-redundant sample files was facilitated by the fastx_collapser function of Fastx toolkit v0.0.13^34^ and subsequently split into 10 (−n 10) files using split function of pyfasta v0.5.2^35^. Comparative alignment to a localised NCBI genomic database (2019) was facilitated by BLAST^26–28^ using task_blastn function and a word_size of 14. BLAST results were input into MEGAN v6.2.1^31^ using a bit-score of 40 within the top 10% of best alignments, and the default “naïve” LCA algorithm. A minimum of 1% of the total assigned reads was necessary to accept a taxon as present and use for downstream analyses^3,21^. Passing sequences were aligned to their corresponding genome using the original cut fasta file and BWA v07.5a.r405^29^ aln function with a disabled seed (−l 1000). SAI file alignments were converted to SAM format using BWA samse. Sequences were filtered using a MAPQ of 25 and sorted using SAMtools v1.3.1^30^. A threshold of 250 total aligned reads were required to proceed with downstream analysis.

Duplicate sequences were removed from resulting files using three methods:

- SAMtools’ “rmdup” function with the option for removal of single-end matches at the 5’ location only^30^
- Picard Tools “MarkDuplicates” function with the option for removing duplicates from the output file^51^
- aweSAM_collapser as specified by developers^54^

Percentage difference was calculated from the variation between reads passing each duplicate removal process.

### The fully developed bioinformatic pipeline

Adapter sequences are removed using cutadapt v1.9.1^24^ with a minimum sequence length of 28bp based on smallest ancient fragments retrievable. A minimum overlap of 1 is used to reduce over-cutting of bases that closely match an adapter sequence. Non-redundant samples are created using the fastx_collapser function of the Fastx toolkit v0.0.13^34^ and split into 10 (−n 10) files of similar size using the split function of pyfasta v0.5.2^35^. The resulting files are considered representative of 10% of the initial non-redundant sample sequences.

The most commonly occurring genera are identified by cross-referencing trimmed representative non-redundant sequencing data with a localised genomic database downloaded (2019) from NCBI. Cross-referencing for genera identification is facilitated through BLAST^26–28^ using task_blastn and a word_size of 14.

Resulting output files are imported into MEGAN Community Edition v.6.2.1^31^ for taxonomic assessment. LCA parameters use a bit-score of 40 within the top 10% of best alignments, and the default “naïve” LCA algorithm. To reduce chances of false positive identifications of taxa at the family or genus level a minimum of 1% total assigned reads is necessary to accept a taxon as present and use for downstream analyses^3,21^. MGmapper^44^ can be employed in additional identifications of genera using default settings and a minimum score of 20.

Following acceptance of a taxon, samples are aligned to their corresponding genome using the original trimmed fasta file and BWA v07.5a.r405^29^ aln function with a disabled seed (−l 1000) allowing damaged sequences to align. Sequences are mapped and filtered for a minimum MAPQ of 25, then sorted using samtools v1.3.1^30^. At this point a minimum threshold of 250 total aligned reads are required to proceed with downstream processes. Duplicates are removed using aweSAM_collapser.sh shell script54, allowing users to read from both the 5’ and 3’ coordinates and retain the read with highest MAPQ. MapDamage2.0^49^ is used to quantify DNA damage through the presence of C to T substitutions on the 5’ end and G to A substitutions on the 3’ end of the sequences. A minimum value of 5-10% on both ends is used for a taxon to be identified as ancient. Read lengths are calculated through cumulative observation and quartile calculation. Phred quality scores and %GC are assessed using FastQC^25^ (Figure 3).

**Figure 3.**
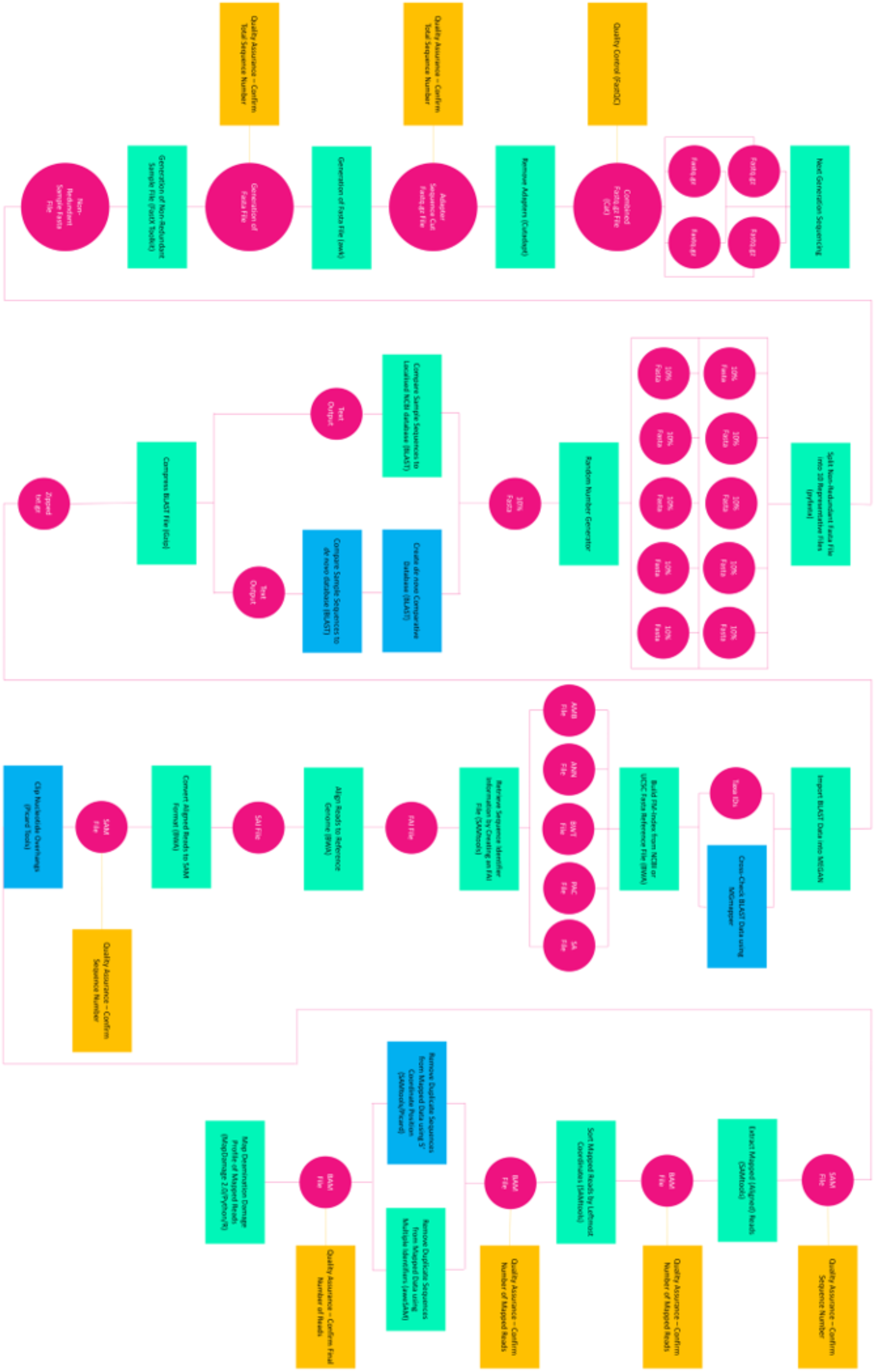
Schematic Overview of the Developed Bioinformatic Pipeline. Processes and their associated software are represented by rectangular boxes. Teal boxes represent processes that form part of the core bioinformatic pipeline. Yellow boxes represent quality control and assurance steps. Blue boxes represent optional processes within the bioinformatic pipeline (see SI). Pink circles represent process output generated (files).

## Supporting information

Supplemental User Manual

## Acknowledgements

We would like to thank Dr Eileen Reilly (Site Entomologist), Dr Nora Bermingham (Site Director) and Jacqueline McDowell (Historic Environment Division, Department for Communities, Northern Ireland) for access to the Drumclay samples used within this study. Likewise, we thank Dr Mareike Stahlschmidt (Site Geoarchaeologist) for providing the Satsurblia samples used within this study. We thank the Computational Shared Facility at the University of Manchester for use of its systems and we would further like to thank George O. T. Merces for his insights into the structure of this paper. This project was funded by the Medical Trainee PhD Scholarship, Anatomy, School of Medicine, UCD awarded to T.C.C.

## Author Contributions

T.C.C. conceived the project. R.N.M.F. and R.P. supervised the project. T.C.C. wrote the paper with contributions from K.D., J.D.O.R., R.N.M.F and R.P. Ancient DNA analysis was performed by T.C.C. Bioinformatic pipeline development performed by T.C.C. Bioinformatic training provided by K.D. and J.D.O.R. Samples bioinformatically processed in University College Dublin by T.C.C. and in University of Manchester CSF3 by K.D. and T.C.C. Bioinformatic assistance provided by J.D.O.R. and K.D. Sample access facilitated by T.M.

## Supplementary Information

### Next Generation Sequencing

Modern day next generation sequencing platforms (Illumina) use a process known as sequencing by synthesis. In this process, a DNA library is prepared into a sequencing sample pool by fragmenting DNA into pieces no larger than 200 base pairs (bp). Custom identifying adapters are added to the ends of a DNA strand and the sample pool is flowed across the solid surface of the flowcell^1^. A flowcell can be described as a glass slide with channels through which polymerases, deoxynucleotide triphosphates (dNTPs) and buffers can be passed. Each nucleotide passing into a flowcell is annealed to a complementary fluorescently tagged dNTP that also serves as a reversible terminator for polymerisation^1,2^. The flowcells allow for clonal clustering of DNA molecules in a PCR process known as bridge amplification^3,4^. The clustering allows for greater fluorescence and improved image analysis. Each flowcell consists of four lanes supplied by a single reservoir, allowing for simultaneous processing of millions of template DNA molecules.

Unlike other sequencing devices, NextSeqs devices employed here use 2-channel sequencing; only two types of fluorescent dyes are used, red and green^5^. “C” nucleotides are tagged with a red fluorophore, “T” with a green fluorophore and “A” nucleotides are tagged using both red and green fluorophores (imaging as yellow). The remaining “G” nucleotides are not tagged with a dye. Towards the end of each sequencing cycle a red and green image are taken and a cleaving enzyme released to remove the fluorophore tagged dNTP for the next annealing cycle to begin and so on. Resulting image data is converted into nucleotide sequences for each of the flowcell lanes^5^. This data is output as four fastq.gz files of similar size. These files form the source material for subsequent bioinformatic assessment.

### The Bioinformatic Pipeline: USER MANUAL

Here, we present a detailed step-by-step instructional manual for the assessment of metagenomic ancient DNA (aDNA) from anthropogenic sediments. Each step will highlight why a process is undertaken, what a process does to the inputting genomic data and how to perform each process using a Linux based operating system. Examples are provided for each step. Note that not all Linux operating systems are the same, however the examples throughout should function on a bash-based Unix shell environment. The shell environment will be referenced as “Terminal” throughout.

#### Getting started: How to set up a PATH

In order to begin calling commands in your Terminal, it is important that they are located in a folder that is part of your PATH on your computer. This is a list of directories your Terminal will search to find the requested executable command. In Linux, common folders that exist on your PATH by default include /usr/bin (common utilities), /usr/local/bin (user-installed executables), /usr/local/sbin (user-installed executables) and /usr/sbin (system daemons and system utilities). By typing the below command into your Terminal, you will see what folders are currently in your PATH:

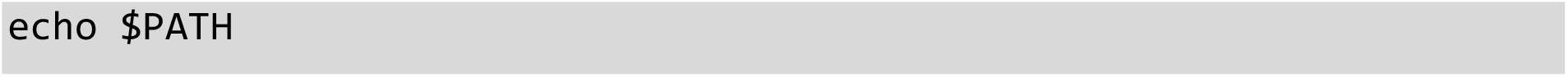

Where $PATH is a global variable in your Terminal to define where executable commands are located.

In instances when you are installing new programs, you may choose not to include them in your computer’s default PATH directories. This allows you to maintain custom programs in a separate directory as well as ensure that default programs are not overwritten by a program of the same name. In order to add a custom directory to your PATH, you can type the below command:

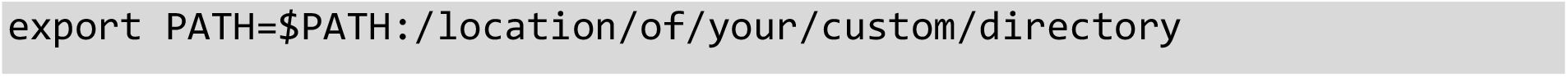

It is important to note however, that the above setting will modify your PATH for that shell session only. Additional shell sessions will not load the custom directory into the PATH. In order to ensure that your custom directory is loaded into the PATH at every shell launch varies depending on the Linux distribution being used. Here, we will use the **.bash_profile** file for the bash shell.

In a new shell window, type the below command to launch **vim**, a command line text editor using the Terminal window.

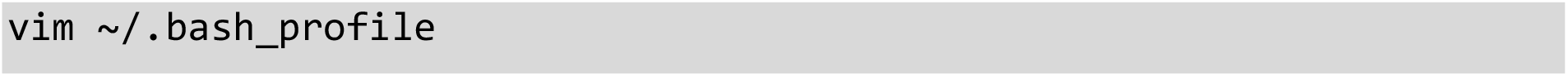

Press the **i** character to enter insert mode. Next, add the export line described above to this file. Once inserted, press the **Esc** key followed by typing **:wq!** and then press **Enter**. **Esc** will exit insert mode, the : will enter a command, **w** will write to the file being edited and **q!** will exit the vim program.

Now, with each launch of your shell, your custom directory will be loaded into the PATH.

#### Step 1. Concatenate Sequencer Output Files

Upon completion of a NextSeq sequencing run, the system will output four zipped files (fastq.gz) for each sample sequenced. In this step the user will combine these four files into one large zipped file representing all extracted sequences for the given sample. The **cat** program, short for “concatenate” (meaning “to link”) allows the Terminal to read multiple files sequentially and output them to the Terminal. To run from the Terminal, navigate to the directory containing the desired NextSeq output files and type:

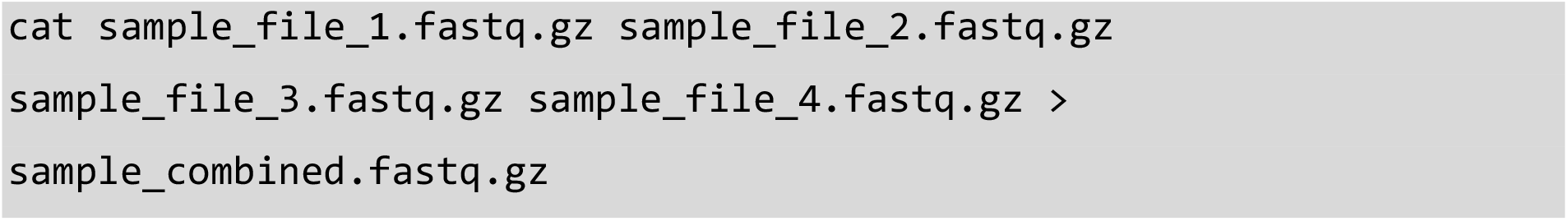

*Example:

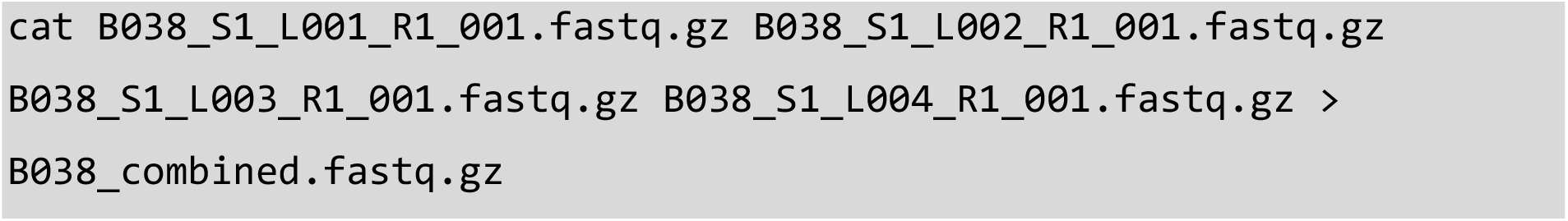

*In the example “B038” represents a unique sample code

The redirection command, executed by using **>**, is used to write the Terminal output to a specified file, instead of to the Terminal output, known as standard out or **stdout**. This file will either be created if it doesn’t exist or overwritten if it does exist.

Note that within the NextSeq output file names the “L001-4” refers to the lane within the sequencing flowcell in which the data originated. Combining the four files into one reduces the need for repeated processing at subsequent steps.

#### Step 2. Quality Control of NGS data (FastQC)

FastQC^6^ is a software package that allows the user to perform quality control checks on raw sequence data coming from NGS platforms. FastQC can be run as a program or directly using the Terminal. To run from the Terminal, navigate to file directory and type the following:

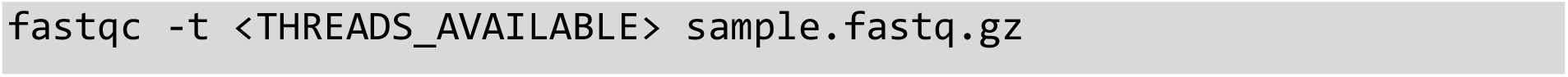

Example:

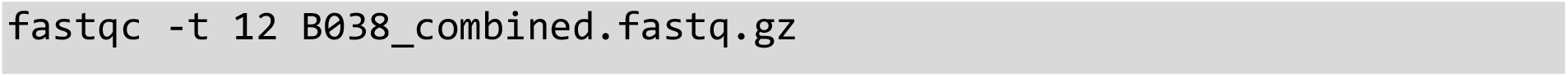

The **-t** option refers to the amount of processing power or “threads” the user wants to dedicate to the program. Increasing the amount of threads available will increase the processing speed. For a large file, such as those resulting from metagenomic studies, we would recommend using a minimum of 12 threads. The number of threads available to your program would depend on the number of processors and cores each processor has in your computer.

This will result in the generation of two files: a **sample.fastqc.html** containing the quality analysis and a zipped folder (containing the same information).

From the “basic statistics” module shows the total number of sequences, sequences tagged as poor quality, largest sequence length, and %GC can be viewed.

The “per base sequence quality” module displays the average phred quality score (Y axis) of all sequences by base position within a read (X axis). For aDNA sequences, the user should still obtain high phred quality scores for the majority of sequences (preferably above 30). Phred scores are important for understanding the quality of obtained sequences^7^. The higher the phred value the better base cell accuracy is achieved (Table SI 1).

**Table SI 1.**
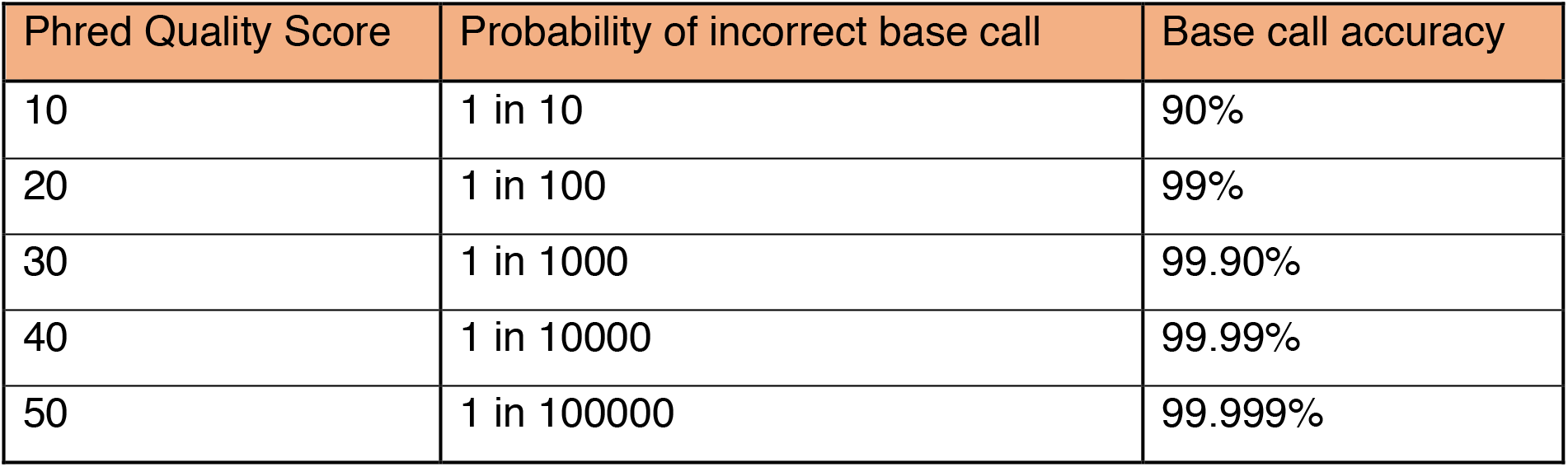
Understanding Phred Quality Scores and Base Call Accuracy. The central column shows the probability of an incorrect base call associated with a phred quality score and the representative accuracy of sequence composition.

The “per base sequence content” module displays the base content (Y axis) of all sequences by position in a read (X axis). In a random DNA library, the user would expect little to no difference between the bases of a sequencing run, meaning the lines in the plot should run parallel with each other. It is likely that FastQC will flag this module with a warning due to adapters changing the value of “G” and “C” base content (GC) (Figure SI 1A)^6^. Upon trimming the sample of adapters (step 3), the user can check the sequences again to see little difference between the bases from the 10^th^ to 74^th^ base pair (10-74bp).

For ancient DNA samples, it is important to note that the first 9 base positions are individual measures that tend to deviate by 10% due to the addition of oligonucleotides at the library preparation phase. The subsequent positions are binned averages, resulting in what should be smooth lines (Figure SI 1B) between 10-74bp. It has been noted that % base composition at the ends of reads that have undergone aggressive adapter trimming (such as that in step 3) are likely to be spurious with sudden deviations in composition^6^. For this reason, it is likely a trimmed sample will flag as failed with the last base position falsely deviating by more that 20%.

**Figure SI 1.**
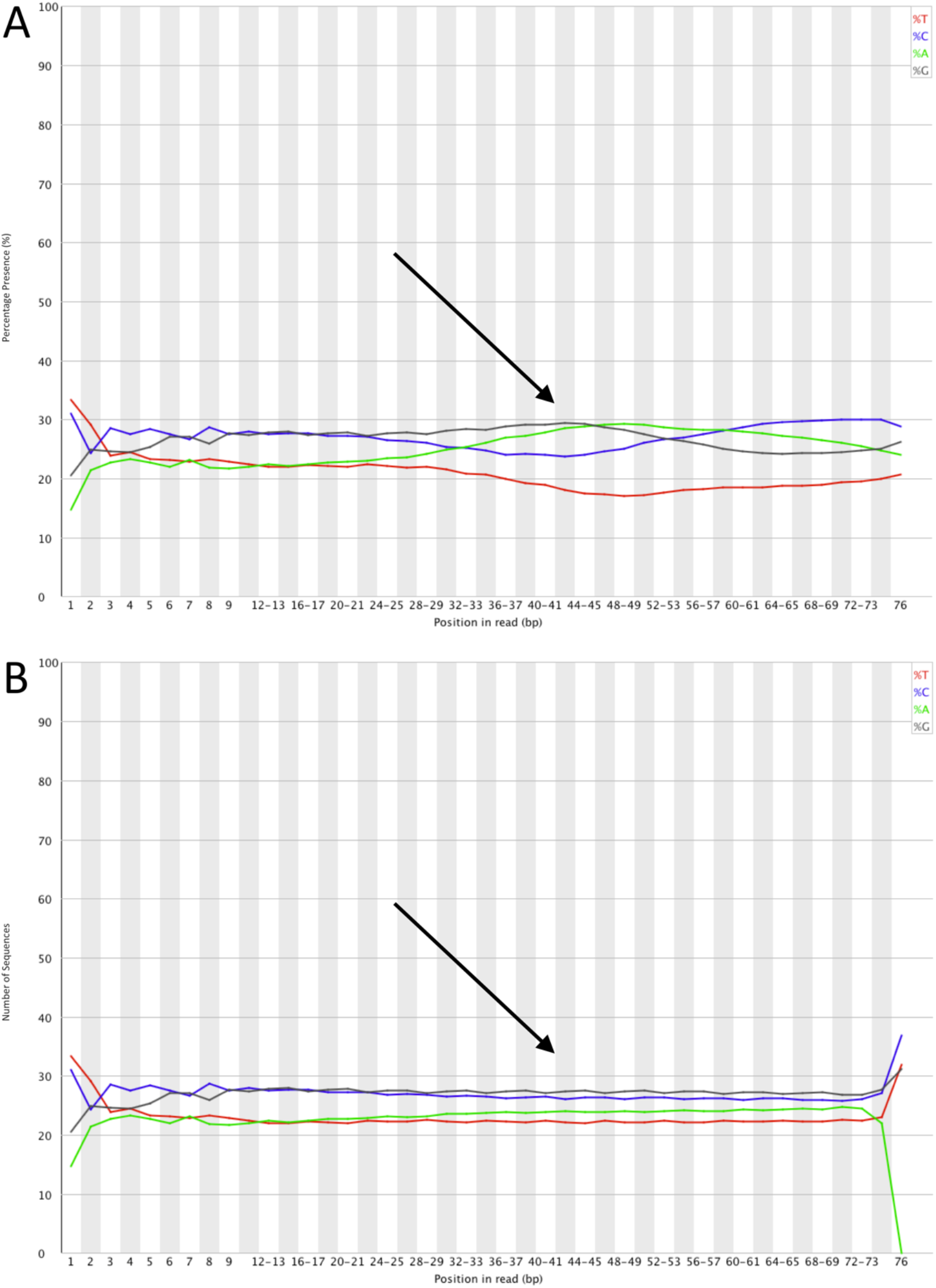
Sequence Content Across all Bases Using FastQC. (**A**) Prior to adapter trimming. (**B**) Post-trimming of adapters. The arrow highlights an area of difference.

The “per sequence GC content” module measures the mean GC content (X axis) across the length of each sequence (Y axis). Typically, a user would expect to see normal distribution of GC content where a central peak corresponds to the overall GC content of the underlying genome (i.e. 41.6% in the *Homo sapiens* genome if a sample is taken from human bone). While a small shift of an expected GC distribution is indicative of a systematic bias independent of base position, an unusual shaped distribution typically indicates the presence of other underlying genomes^6^, or contamination. As the FastQC program does not know the GC content of the underlying genome, the modal GC content is calculated from observed data and used to build a reference distribution. A warning is raised when the sum of deviations from the modal distribution represents more than 15% of reads (Figure SI 2A). Failure is flagged when deviations amount to more than 30% of all reads. By its very nature a metagenomic sample it is expected to have multiple underlying genomes represented and thus a warning or failure of this module is expected. Upon extracting the mapped sequences (step 19) aligned to a desired genome, the user can import into FastQC and compare the mapped %GC to compare to expected GC content. In this case the modal GC content should conform to the desired genomes overall %GC, with a normal peak distribution pattern (Figure SI 2B).

**Figure SI 2.**
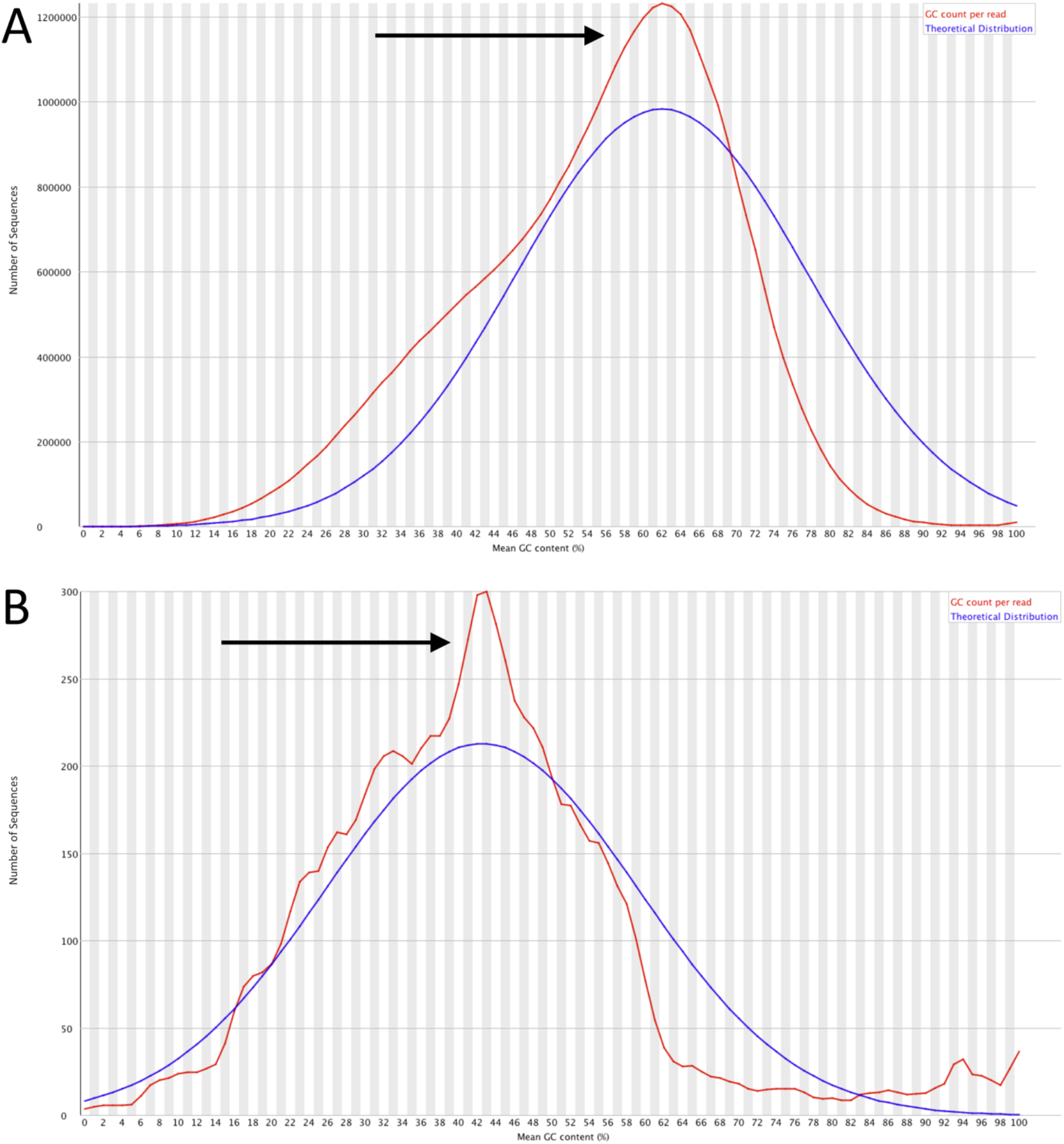
GC Content Across all Sequences Mapped to a Desired Genome (i.e. *Homo sapiens*) Using FastQC. (**A**) Prior to the extraction of mapped sequences. (**B**) Post extraction of mapped sequences with expected %GC for *Homo sapiens*. The mean percentage of “G” and “C” bases are plotted for all sequences within a sample. The arrow highlights an area of difference.

The “sequence length distribution” module measures the sequence length (X axis) over all sequences (Y axis). If the user performs FastQC prior to the removal of adapters (step 3), the length distribution will always peak at 76bp as the max read length setting on the NextSeq platform using the 500/550 high output V2 (75 cycle) reagent kit (Figure SI 3A). Upon trimming the sample of adapters (step 3) the user can check the sequences again to see a more distributed sequence pattern. Note that the module will flag a warning if all sequences are not the same length^6^. For ancient DNA samples, following the trimming of adapter indexes, the user should expect a distribution of sequences, usually accumulating in a short peak around the 35-50bp mark and a larger peak at the 74-76bp mark (Figure SI 3B).

**Figure SI 3.**
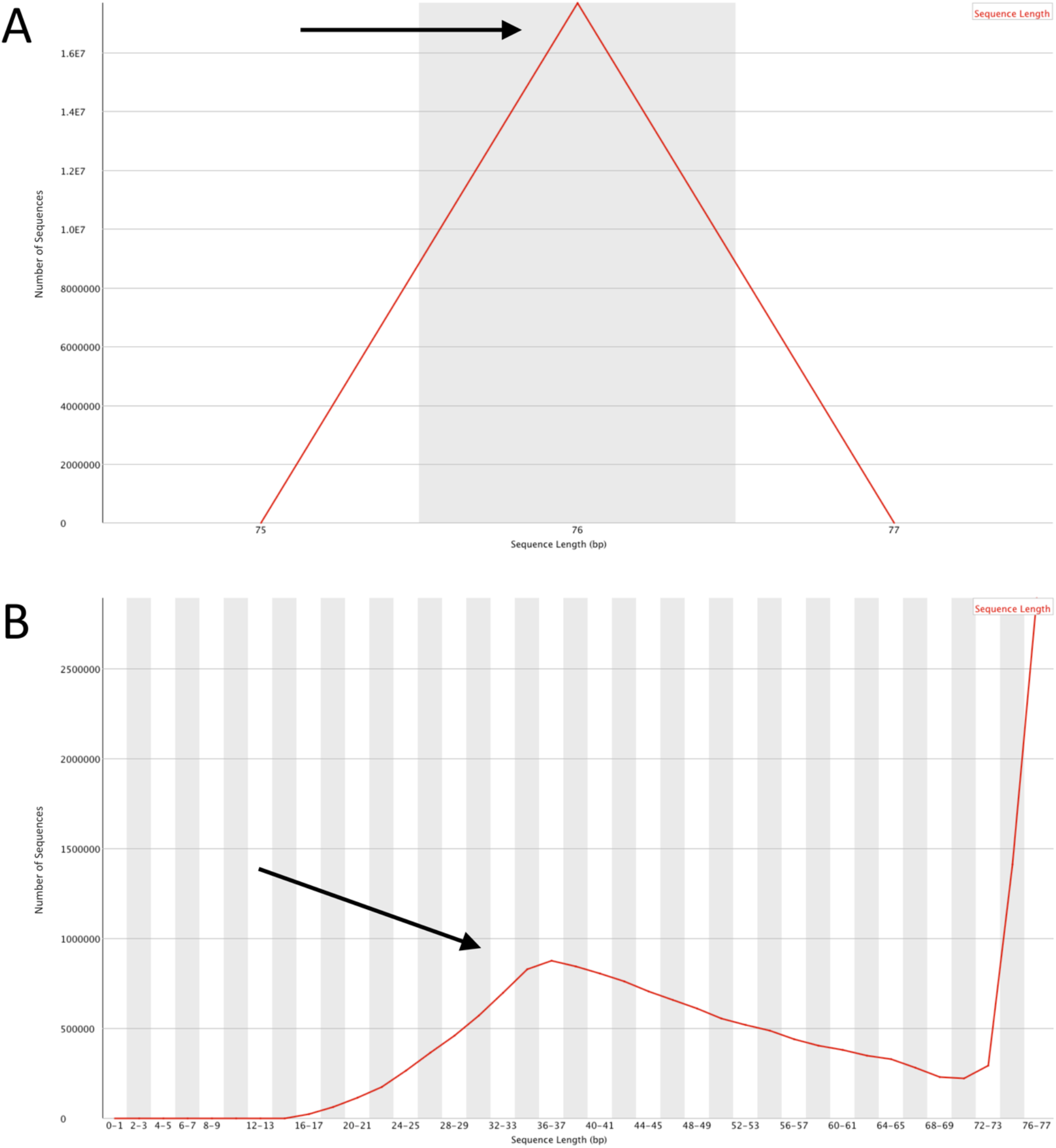
Distribution of Sequences by Length (bp) Using FastQC. (**A**) Prior to adapter trimming. (**B**) Post-trimming of adapters. All sequences present within a sample sorted according to length of sequence strands. The arrow highlights an area of difference.

The “adapter content” module displays a cumulative percentage of the proportion of the user’s library (Y axis) which has seen adapter sequences at each position (X axis). Once an adapter sequence has been read, it is counted as present through to the end of a read, thereby increasing the percentage of adapter content as read length continues. The user should expect to fail this module (adapter presence on more than 10% of reads) if analysing sequences prior to the trimming of adapters. Post removal of adapter sequences (step 3), the user should expect to pass this module with 0% of all sequences showing evidence for adapters.

#### Step 3. Removal of Adapter Sequences (cutadapt)

Adapter sequences are synthesised oligonucleotides that can be added to the ends of DNA for a variety of applications. In aDNA research, adapters provide two functions, to stabilise authentic damaged sequences and to attach index sequences for sample identification after sequencing^8^. Post sequencing adapters are removed to leave only authentic DNA sequences.

**cutadapt** is a software package that allows users to search all reads within a given sample for a specified adapter sequence and filter reads according to a minimum desired threshold, otherwise known as trimming or a cut^9^. To run from the Terminal, navigate to the directory containing the **combined.fastq.gz** file (from step 1) and type:

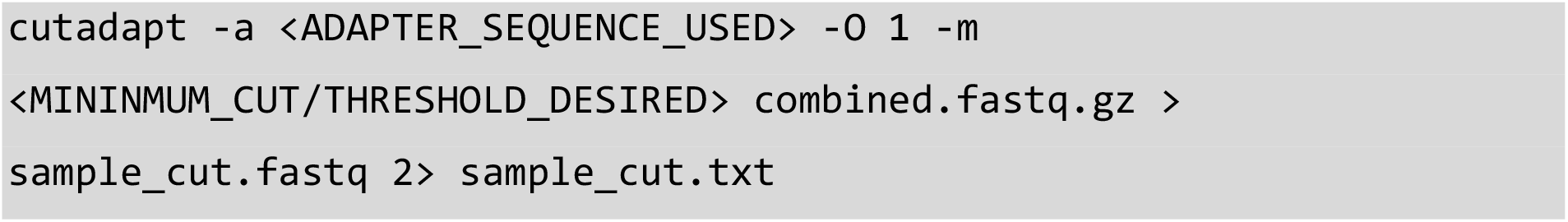

*Example:

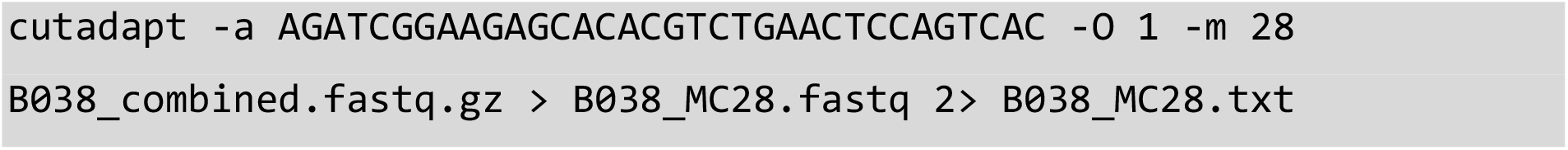

*In the example “MC28” refers to minimum cut (length) used.

The **-a** option refers to the adapter sequence used during the library preparation phase ligated to the 3 prime (’) end of a sequence. The adapter and anything that follows is removed and discarded from a sequence. For aDNA the majority of authentic aDNA sequences are shorter than the maximum sequencing read length (76bp), thus part of the adapter is usually found at the 3’ end of reads^10^.

The **-O** option is the minimum overlap length of bases that can match between the specified adapter and a read before removal of those bases. If a read contains a partial adapter sequence shorter than the minimal overlap length, no match is registered and no bases removed. For **cutadapt**, the default setting is “3” (bp)^9^, for aDNA “1” (bp) should be used, meaning if overlap between the adapter and a read is shorter than 1bp the read is not altered, thereby reducing the number of authentic aDNA bases from being removed erroneously.

The **-m** option parameter allows the user to discard processed reads that are shorter than the specified length (bp) after adapter removal. For aDNA a minimum length value of 28bp is used, being the lowest identifiable size of authentic ancient reads.

This program uses a standard input (**stdin**) and output (**stdout**) format, allowing the user to output both a trimmed sequences file (using the redirection command >), and a redirected standard error stream (**stderr**) and discarded reads report in the format of a text file (using the redirection command **2>**). Upon completion of **cutadapt** the user may want to quality check sequences using FastQC as outlined in step 2.

#### Step 4. Quality Assurance of NGS Data (1): Confirm Total Number of Sequences (1)

Occasionally errors can occur with the format conversion, manipulation, or unzipping of a file, it is best practice to introduce checks to ensure data is consistent (integrity) and reduce possible complications with downstream bioinformatic processes.

The **wc** utility program allows users to bioinformatically count the number of lines, words, and bytes contained within the specified input file

The addition of the **-l** option instructs the command to only count the total number of lines within the specified input file. The number of lines counted can then be divided by four to give the user the total number of sequences within the trimmed FASTQ file.

The total number of lines counted are divided by four on account of the formatting of FASTQ files. Each sequence present is represented by four lines of code^11^. The first and third line consist of sequence identifier information with first line identified with an “@” symbol, and the third a “+” character, the second line contains the raw sequence data in IUPAC convention, and the fourth encodes quality values for the sequence data within the second line.

Using the Terminal, navigate to the directory containing the trimmed **sample_cut.fastq** file and type the following:

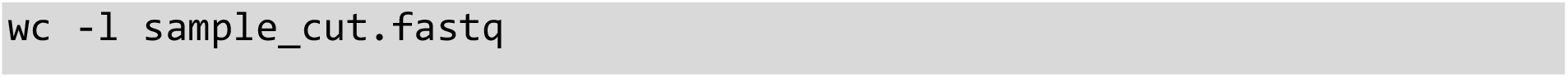

Example:

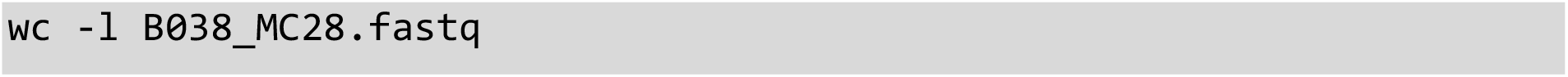

Divide the resulting number by four to get the true sequence value. Then, using the **sample_cut.txt** file output in step 3, check the summary section for “Reads written (passing filters)”, this will display the total number of sequences within the trimmed **sample_cut.fastq** file. Both numbers should match.

#### Step 5. Convert FASTQ File to FASTA Format

The conversion of FASTQ files to FASTA format is performed for a number of reasons. First, given that per base sequence quality has been assessed within step 2, the base quality scores are no longer need for downstream processing and as such can be removed to reduce file size, this is automatically performed during conversion into FASTA format. Secondly, the reduction in working file size gives the user the added benefit of faster processing speeds. Lastly, FASTA format has been widely used within bioinformatics for over 30 years, as such many programs used for downstream processing such as BLAST^12^ rely on this format. To begin formatting, navigate to the directory containing the **sample_cut.fastq** file and type:

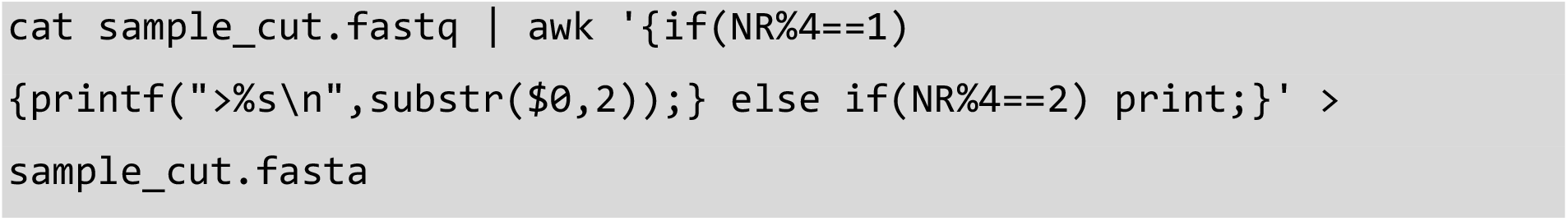

Example:

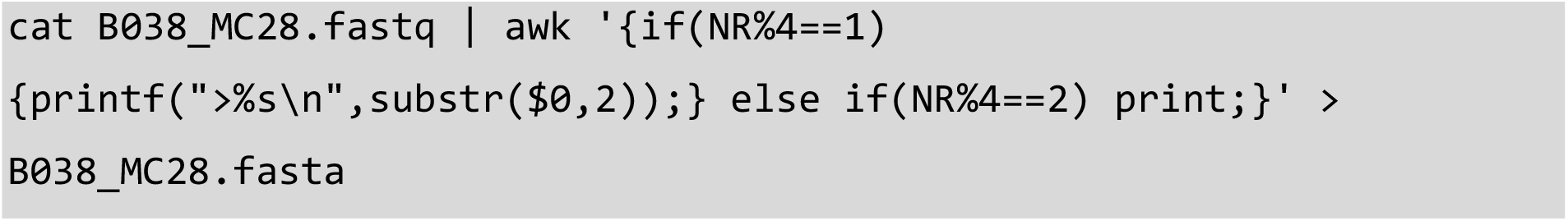

The **cat** utility program is used to read the contents of the FASTQ file and ‘pipes’ the **stdout** into the **awk** program as **stdin**. Pipe, represented by the **|** character, is a command line program which allows the output of the 1^st^ program (in this case **cat**) to serve as the input into a 2^nd^ program (in this case **awk**), like a pipeline.

The **awk** program is a tool to write simple yet effective statements in order to manipulate text. It defines text patterns that are to be searched for in each line of the input file. Here the **awk** program reads that if the Number of Records variable (NR), also defined as the line number, when divided by four has a remainder of 1 (denoted as the **%** character in **NR%4==1**) then it will format the line as follows, the **printf** command is invoked and this will print a > character, followed by a format specifier to specify the text as a string (**%s**) followed by a newline character (**\n**) with the string to be decided using the **substr** command. **$0** specifies the whole line to be inputted, and 2 specifies to get the whole line from character **2** onwards (this forms the sequence header), otherwise if the NR variable when divided by four has a remainder of 2, then print the corresponding line (the raw sequence data). Lines 3 and 4 are ignored from the **awk** command. Finally, this manipulated text file is written to a FASTA file using the redirection command **>**.

#### Step 6. Quality Assurance of NGS Data (2): Confirm Total Number of Sequences (2)

As mentioned in step 4, occasionally errors can occur when formatting, manipulating or unzipping a file. It is therefore best practice to check the number of sequences within a file remain consistent with the originating file. To do this we again use the **wc** utility program with the **-l** parameter.

Unlike FASTQ format, FASTA files consist of two lines of code^11^. The first line consists of sequence identifier information or a short description for the sequence, while the second line contains the raw sequence data encoded to IUPAC conventions. As such the resulting total number of lines counted are divided by two to get the true number of sequences present within the FASTA file.

Within the Terminal, make sure the directory contains the **sample_cut.fasta** file, and type:

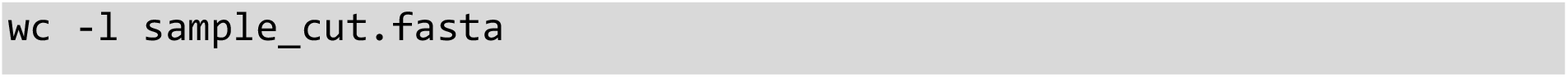

Example:

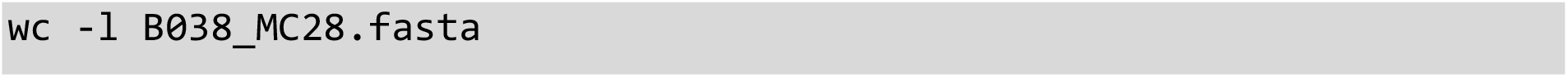

Divide the resulting number by two for the total number of sequences present. The number should match the total number of sequences present in the originating FASTQ file outlined in step 4.

#### Step 7. Create a Non-Redundant Sample File (FastX-Toolkit)

A non-redundant sample refers to one without sequence repetitions, otherwise referred to as duplicates. The more sequences present within a sample file, the longer and less efficient the processing speed of downstream bioinformatic processes. This is a well-documented issue when comparing sample sequences to a genomic database^13,14^ (Step 10).

FastX toolkit is a package of command line tools for the pre-processing of FASTQ and FASTA files^15^. The collapser program **fastx_collapser**, merges all duplicate sequences for a region of coding into a single representative sequence, while maintaining read counts. This is performed for all duplicates within a sample until only unique reads remain.

To run the program in the Terminal, navigate to the file containing the **sample_cut.fasta** and type the following:

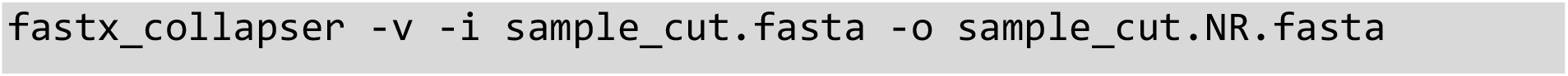

*Example:

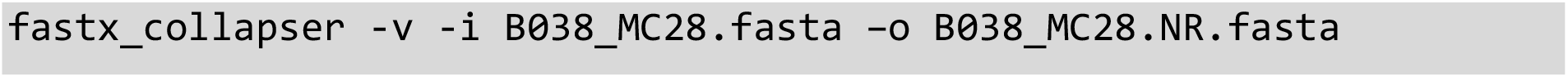

*In the example **NR** refers to a non-redundant sample file.

The **-v** option instructs the command to verbose the number of processed reads within the given sample.

The **-i** option refers to the input file, in this case the **sample_cut.fasta**

The **-o** option is the outputting non-redundant file. The resulting output file, should be drastically reduced in size to the inputting FASTA file.

Note, do not remove the original FASTA file as this will be used for in-depth alignments downstream (step 15).

#### Step 8. Split the Non-Redundant Sample File into Ten Files of Equal Proportion (pyfasta)

To further improve the processing speed of comparative analysis (step 10), the non-redundant sample file can be split into ten files of equal proportion. As multiple regions of coding are extracted from origin sources of DNA, the splitting of a non-redundant sample will not affect the identification of a taxon’s presence within a sample.

**pyfasta** is package of tools utilising python via a command line interface for the preprocessing of FASTA files^16^. The split command **pyfasta split** allows users to distribute sequencing reads between multiple files via various means.

Here, the **-n** option is used, referring to the number of outputting files desired. Ten files will result in each being 10% representative of the total sequences, while two files represent 50% each.

To run from the Terminal, navigate to the directory containing the non-redundant **sample_cut.NR.fasta** file, and type:

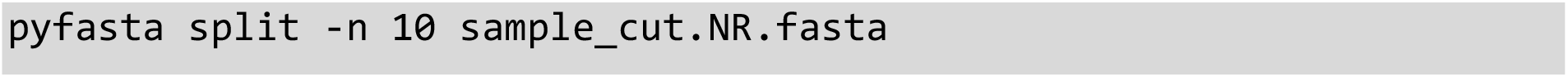

*Example:

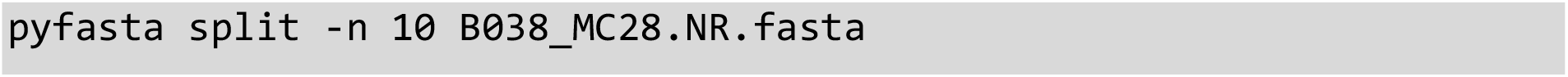

*Outputting files will be labelled 00 – 09, this is useful for the following step (step 9).

To validate that resulting files are representative of an entire samples sequences, the mean percentage difference, and standard error between expected hits based on the 10% files and actual hits achieved with the 100% file were calculated. The expected total hits predicted by the 10% file was accurate to the 100% file within −0.007% (±1.101 SEM). Representing a difference of 0.07 hits within 1000.

#### Step 9. Randomly Select One of the Non-Redundant Sample Files

In order eliminate the possibility for selection bias. A randomised number generator is used to determine which of the representative files (step 8) would be used for comparative analysis (step 10).

For this, the **/dev/urandom** number generator available in the majority of Linux distributions used. In order to invoke it, the utility program **grep** is used. **Grep** allows users to search plain text data sets for lines matching a regular expression.

To run using the Terminal, type:

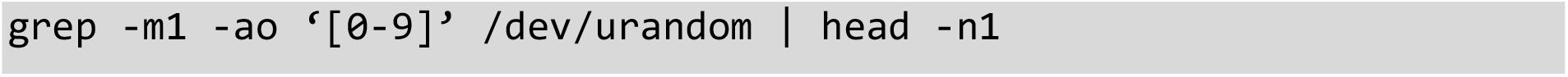

The **-m** option ensures that only 1 line is returned from **/dev/urandom**. **-a** ensures that the line is processed as text and the **o** combination will only print matched, non-empty lines. **-ao** can also be written as **-a -o** in the above program if preferred. A simple regular expression is also specified to provide a single-character number range for **grep** to only return a single-digit number, specified with the **‘[0-9]’** regular expression pattern. These three options along with the regular expression, sanitise the output of the **/dev/urandom** program to ensure only single digit number characters are returned. This is then piped into the **head** command. The **head** command, with the **-n1** parameter will ensure that only the first number piped from **grep** is returned.

The process will output a number corresponding to one of the split non-redundant sample files. This will become the users working file for subsequent steps.

#### Step 10. Option 1. Compare Sample Sequences to a localised NCBI database (BLAST)

Using this option, the representative sample file is used to compare its sequences to the entire National Centre for Biotechnology Information’s (NCBI) genomic database. Basic Local Alignment Search Tool (BLAST) is a multi-platform algorithm that allows users to query sample sequences against a specified database^12,17,18^. To run from the Terminal, navigate to the folder containing the randomly selected representative sample file and type:

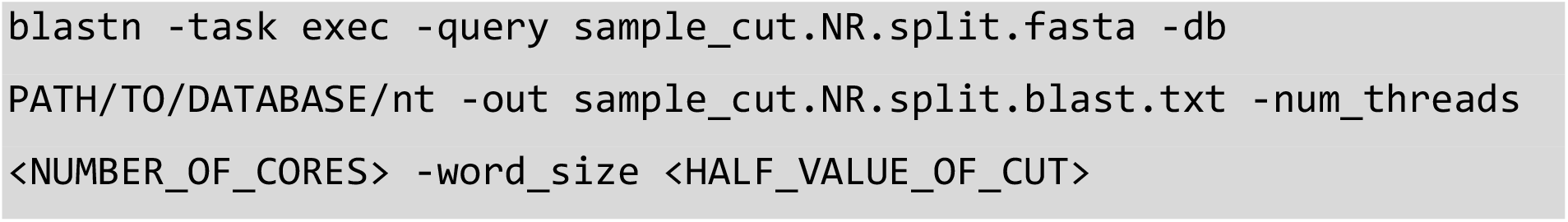

*Example:

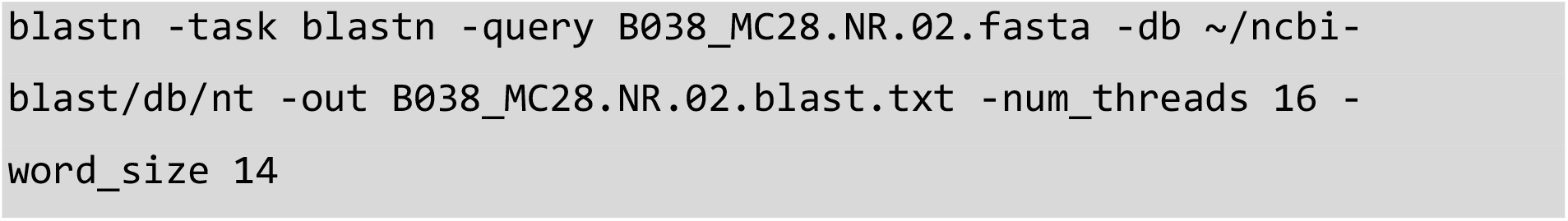

*The **split** refers to the split file (00-09) randomly selected in step 9. In this example **02** is used.

The **-task exec** option allows users to specify the type of search parameters best suited for the sample. The **-task blastn** and **-task blastn-short** options are best suited for interspecies comparisons using short sample sequences^19^. The later optimised for sequences shorter than 50bp. However, as most aDNA is 30-70bp in length^10,20,21^, we recommend the user utilise the **task blastn** option.

The **-db** option refers to the location of the **database**. Here, the user specifies the path to the genomic database of choice.

The **-num_threads** option allows users to specify the amount of processing power committable to a BLAST search. Each core in a computer processor typically has two threads available, we recommend specifying the maximum number of threads available to the user for fast processing. Thus, for a 32 core unit, a **num_threads** of 64 can be used.

The **-word_size** parameter specifies the number of base pairs required to confirm a match between a sample sequence and a reference sequence. Using **-task blastn** this value is automatically set to 11bp^19^. We recommend using a **word_size** half the value of the smallest available sample sequences. In this case because we trimmed samples (step 3) at a value of 28bp the **word_size** used is 14.

The output or **-out** file, contains all genomic matches and associated expect values. Text or **txt** format is used due to its versatility with any word processing program and subsequent alignment tools.

#### Step 10. Option 2. Part 1. Make a *de novo* Comparative Database (BLAST)

Using this option, the user can create a *de novo* genomic database for comparison with a representative sample file. This option can drastically reduce the amount of time committed to the blasting process. We recommend the use of this option if the user wishes to compare sequences to a specific database such as invertebrates or mammals. This option can also be used if searching for specific genomic alignments such as those identified in a zooarchaeological or archaeobotanical study of a site. However this will introduce a selection bias not representative of a true metagenomic study.

For the purpose of this example, we downloaded the entire invertebrate database in the format of individual FASTA files following NCBI guidelines^19^. Once downloaded, the reference FASTA files (ending in the file extension **.fna**) are placed into a new directory folder and concatenated into a single FASTA file using the **cat** program (step 1). To run, open the Terminal and navigate to the newly created directory folder. Type:

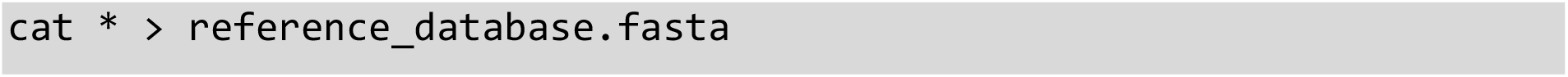

Example:

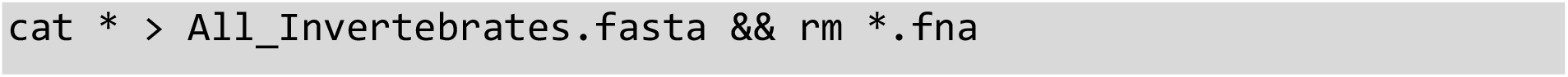

The **\*** wildcard refers to all files within a specified directory.

The **&&** logical operator is used to chain commands together in one execution, allowing a user to achieve a secondary action after the first action has been completed successfully. In this example, the removal of individual FASTA file after concatenation using the **rm** command, leaving only the full concatenated reference file (“**.fasta**”).

To create a *de novo* database, the **makeblastdb** command as part of the BLAST suite is used^19^. The command converts a FASTA file into a reference database. To run from the Terminal, navigate to the directory containing the concatenated reference file and type the following:

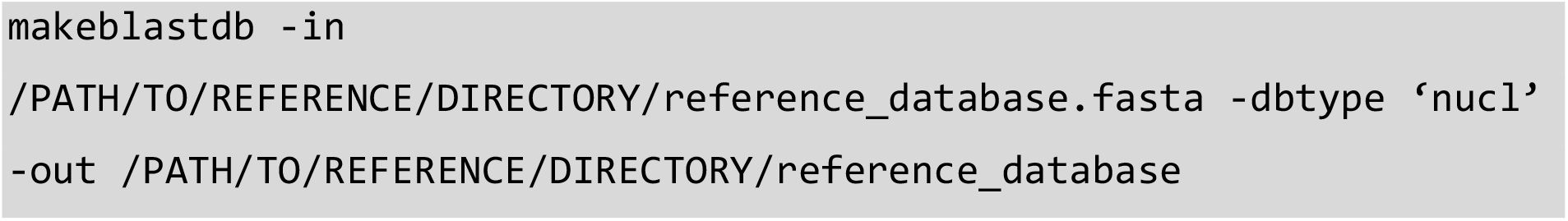

Example:

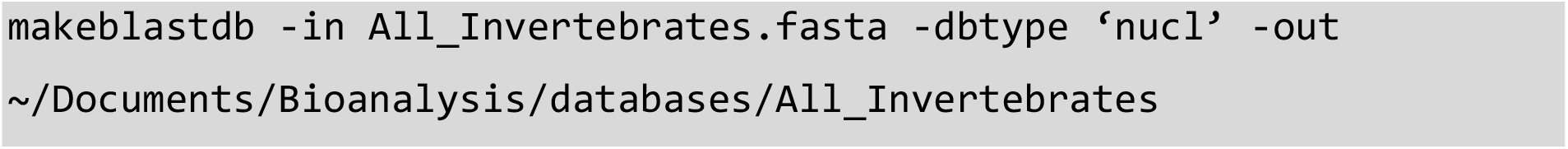

The **-dbtype** option refers to the type of reference sequences used. In this case **nucl** is used, short for **nucl**eotide.

In some situations a user may wish to use a genomic sequence not yet available via the traditional sources of NCBI or the University of California (UCSC)^22,23^. If this is the case, we recommend the user ensure sequences within the desired reference file be assigned unique identifier information to allow for downstream taxonomic assignment. This process should be performed before concatenation. To check this information using the **grep** program type:

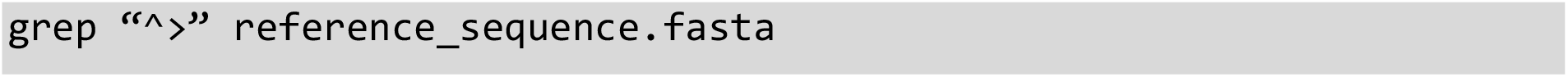

Example:

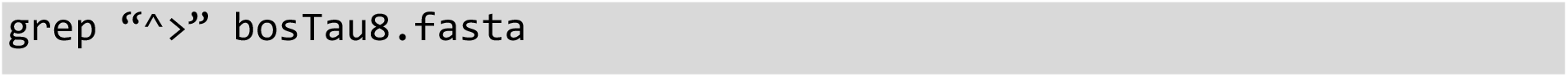

The **^>** regular expression in **grep** will print all lines that begin with the **>** character to the FASTA file specified and print it to the Terminal window as **stdout**

In situations where this information is missing. Identifier data can be added using the following in-line perl script:

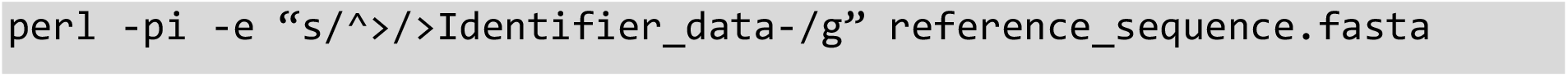

Example:

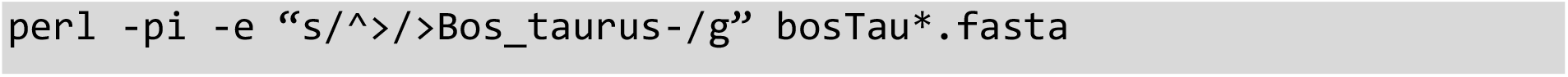

The **-p** option causes perl to assume a while loop will be used in the script. In combination with **i**, **-pi** specifies that files processed by the **<>** construct in the while loop are to be edited in-place. The **-e** option is used to specify one line of script entered into the command line. The script entered here is a search and replace operation to be applied to the FASTA file passed in after it. This is done by beginning with the **s/** operator, followed by the regular expression **^>** which will match all lines that begin with the **>** character, this will be replaced with the Identifier data and a **–** symbol (in the above example this is the text “Bos_taurus-”). Finally, the **/g** modifier applies the regular expression globally, i.e. to the whole file.

In the example above the **\*** wildcard is used to match all files that begin with the letters “bosTau” and end in “.fasta”

#### Step 10. Option 2. Part 2. Compare Sample Sequences to *de novo* Database (BLAST)

As specified previously, BLAST is a multi-platform algorithm that allows users to query sample sequences against a specified database^12^. In this example a *de novo* database is used. In the Terminal, navigate to the folder containing the randomly selected representative sample file and type:

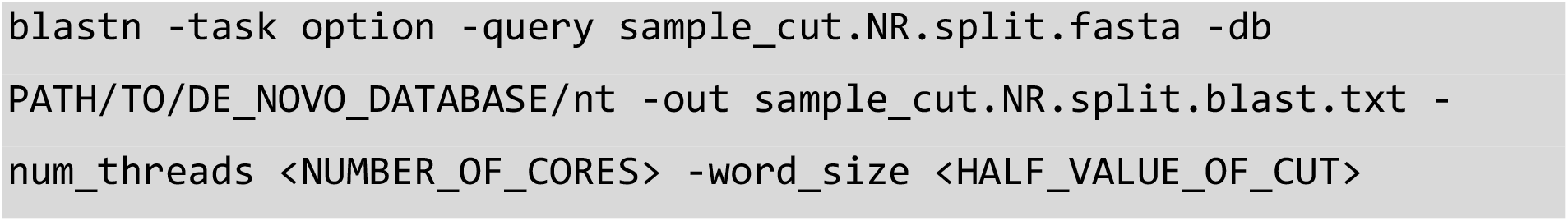

Example:

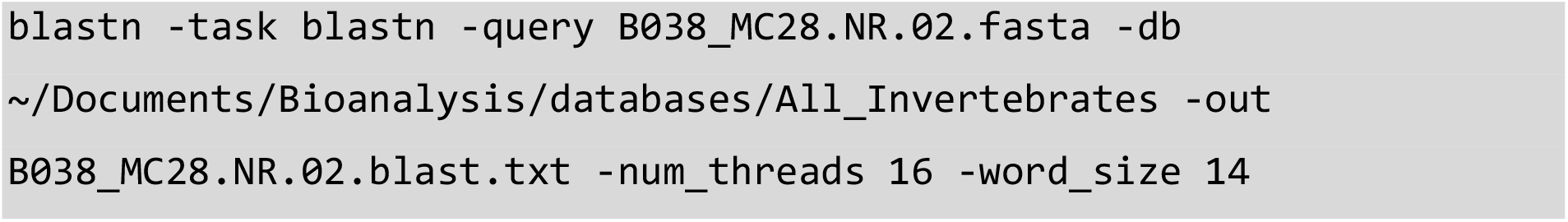

Information regarding BLAST options used can be found in Step 10 (Option 1).

#### Step 11. Compress BLAST file (Gzip)

Following a successful BLAST, it is not uncommon that files will be several gigabytes in size. To optimise storage space and reduce processing time of subsequent assignment (Step 12) we would recommend zipping the “txt” file using the **gzip** program. **gzip** is a single-file, lossless data compression tool, that allows users to specify the level and speed of data compression from worst but fastest (**1**) to best but slowest (**9**). To run in the Terminal, navigate to the folder containing the “txt” file and type:

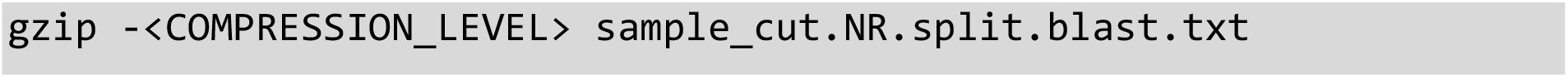

Example:

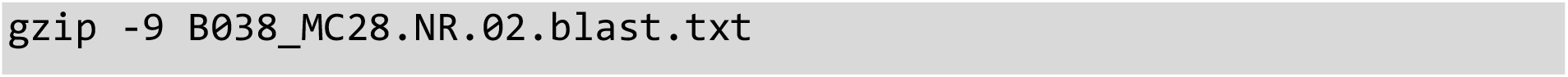

The resulting file will contain the file extension **.gz**

#### Step 12. Import BLAST Data into MEGAN

“MEtaGenome ANalyzer” or “MEGAN”, is a computer program that allows users to import and analyse large datasets of compared genomic sequences (via BLAST or other genomic comparison tools)^24^. During data import, MEGAN assigns a taxon identification to processed read results based on NCBI taxonomy. Using the lowest common ancestor (LCA) algorithm, reads are assigned across a taxonomy (i.e. order, genus, tribe, etc.). The sequences which have a min-score within a specified percentage of the best alignments within a taxonomy are binned into the lowest possible common ancestor position. Sequences aligning to multiple taxa within a grouping are binned into a higher taxonomic level^24^ (i.e. sequences assigned to the genera *Bos* and *Capra* will be binned into the family *Bovidae*).

Sequences are imported using a min-score (bit-score) of 40 within the top 10% of best alignments, and the default “naïve” LCA algorithm. A minimum of 1% of the total assigned reads is necessary to accept a taxon as present and use for downstream analyses^25,26^ (Paper 1, 3).

The resulting taxonomic allocations inform the user on which reference sequences to download for in-depth sequence alignment downstream (step 14). It is important to note that any taxonomic assignments are representative of 10% the total sample pool as outlined in step 8, and can be multiplied by 10 to get the total expected genomic hits by taxonomy.

#### Step 13. Cross Check BLAST Data using MGmapper (Optional)

In some situations, the user may wish to cross-check BLAST data using a second method of mass comparative alignment. MGmapper is a web-based package that allows users to process raw sequencing data and perform reference-based sequence alignments (using NCBI genomic database) with post-processing taxonomic assignment at a species level^27^. It is important to note that distribution amongst an entire taxonomy is not applicable using this method, allowing only for genus and species-based comparisons.

To run MGmapper, search for the following web address: https://cge.cbs.dtu.dk/services/MGmapper/. Using the default settings in double-stranded best-mode with adapter trimming and a minimum alignment score of 20, upload the concatenated FASTQ file from step 1.

In best-mode, reads are assigned to solely one reference sequence after mapping to all specified genomic databases. We recommend using a minimum alignment score half the value of that used for MEGAN, in this case 20. This is because MGmapper’s four criteria to positively identify a taxonomy are optimised for larger sample sequence lengths, and are more prone to false negatives at lower bp lengths^27^.

Genomic assignments (positive and negative) are output into downloadable **xlsx** Microsoft Excel files, accessible for 48 hours online.

#### Step 14. Part 1. Build an FM-Index from a NCBI or UCSC FASTA Reference File (BWA)

In order to perform an in-depth genomic alignment, a reference genome file in FASTA format must be provided. We would recommend using a well maintained and reviewed source for reference sequences such as NCBI or UCSC^22,23^. Reference genomes to be downloaded are determined by the genomic assignments achieved using BLAST (and MGmapper).

Using NCBI as an example for how to download a reference file, visit: https://www.ncbi.nlm.nih.gov/. From the drop-down menu located next to the search bar, select “genome”. Type the name of the species desired within the search bar, preferably in Latin, and click enter. Once the new page has loaded, a box can be seen above the organism overview with three headings including one named “reference genome”. Here, the user can download sequences in FASTA format by clicking the tab “genome”. The download will result in a zipped FASTA file (**fna.gz**). Next, rename the file after the species, place the file within a new directory folder of the same name, and unzip using the gzip decompress command. From the Terminal, navigate to the directory created and type:

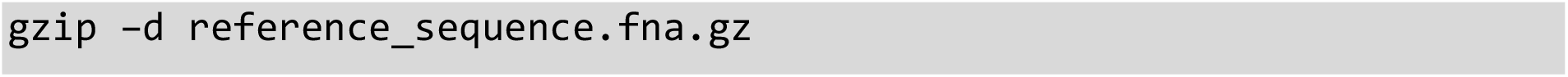

Example:

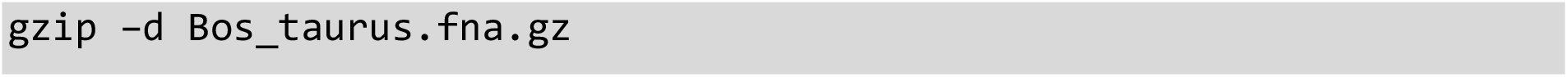

The **-d** option instructs the tool to “decompress” the file selected.

“Burrows-Wheeler Aligner” or “BWA” is a software package for mapping low-divergent sequences to a large reference genome^28^. Because the software uses low-divergent sequences it is considered more stringent than mass alignment tools such as BLAST. BWA requires a specific FM-index set for alignment to take place. This can be constructed using the **bwa index** command. To run in the Terminal, navigate to the directory containing the blasted **txt.gz** file and type:

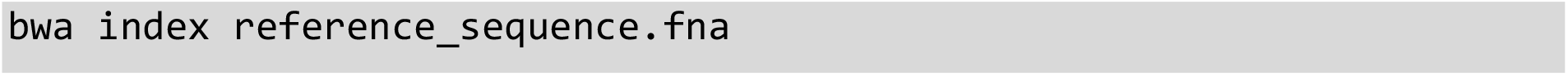

Example:

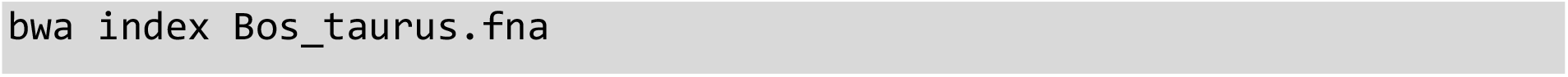

The **index** command will result in six of the required seven index files for an alignment to take place, including the original **.fna** file, these include: **.fna.amb**, **.fna.ann**, **.fna.bwt**, **.fna.pac**, and **.fna.sa**. All files will have an identical base-name, allowing subsequent tools to recognise an index in its entirety regardless of the file extension.

#### Step 14. Part 2. Retrieve Sequence Identifier Information by Creating a FAI File (SAMtools)

The “FASTA index” or “FAI” file contains all sequence identifier information for a reference genome, including chromosome names, chromosome lengths, offset of the first base of each chromosome sequence, and the length of each FASTA line. This information allows subsequent tools or the user to efficiently query specific regions of a reference genome sequence.

SAMtools is a set of utility commands that primarily allows users to view, read, write, and edit SAM, BAM and CRAM formatted files^29^. Additionally, SAMtools can extract sequence identifier information from FASTQ and FASTA indexed reference files. Using a FASTA reference file, a FAI file can be created using the **faidx** command.

To run from the Terminal, navigate to the directory created in Step 14 (Part 1), containing the reference genome **.fna** file. Type:

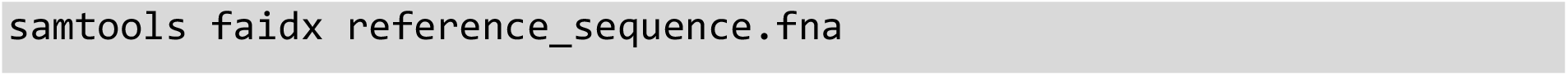

Example:

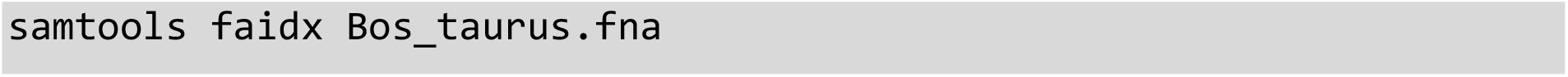

The resulting file will contain an identical base-name with a **.fai** file extension and the **.fna** extension, reading as **reference-sequence.fna.fai**.

#### Step 15. Compare All Sample Sequences to A Genomic Reference Sequence (BWA)

As mentioned in the previous step BWA is a software package for mapping low-divergent sequences to a large reference genome^28^, and is considered more stringent than methods of mass alignment (Paper 2). In this step, BWA is used to align (also referred to as mapping) sample sequences to a single reference genome using the **aln** command. We recommend the use of **bwa aln** over **bwa-mem** for aDNA sequences (30-70bp), as **bwa-mem** is optimised for sequences >70bp^30^, and it has been noted by users that **bwa aln** performs better with reads <70bp^31,32^. In order to perform alignment, the user will need the original FASTA file representing all sample sequences (generated in step 5). Using the Terminal, navigate to the directory containing the original sample FASTA file and type:

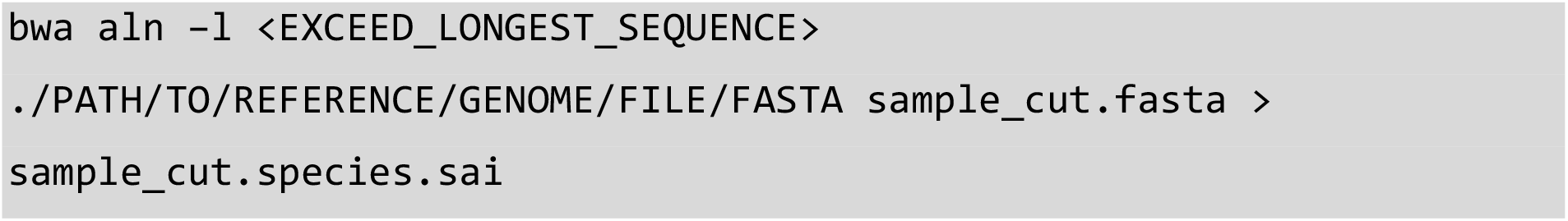

Example:

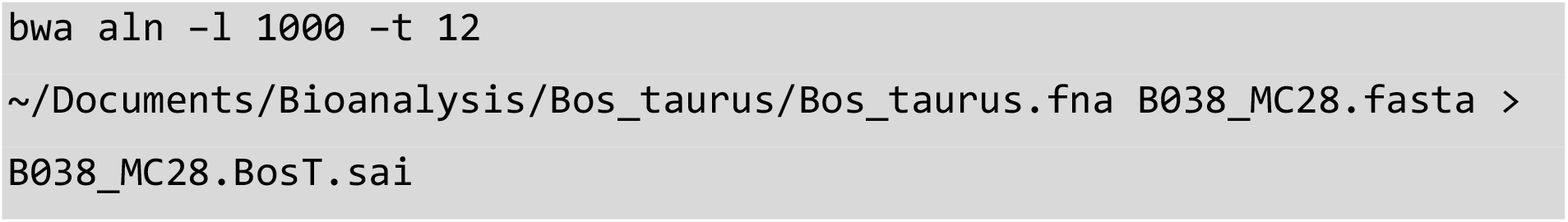

The **-l** option refers to “seed length” as part of the seeding process. Seeding can be explained as the finding of exact matches of part of a sample sequence with part of the reference sequence. The larger the seed length required (i.e. 300bp) the faster the alignment process but greater the chance for loss of accuracy. If used correctly a balance may be achievable. In the case of aDNA fragments, seeding is not recommended, owing to base substitutions and its highly fragmented nature. To disable seeding a seed length larger than the longest sample sequence can be specified, thus allowing damaged aDNA sequences to be aligned^33^. We recommend using a seed length of 1000.

The **-t** option refers to the processing power in the form of “threads” that the user wishes to dedicate to the alignment process. The more threads available the faster the alignment process. Here we use 12.

The resulting file with the **.sai** extension is an intermediate file containing “suffix array indexes” for interpretation and conversion into SAM format.

#### Step 16. Convert the Aligned Reads to SAM format (BWA)

Using the **.sai** file generated through alignment in step 15, BWA is further used to convert aligned sequences into “sequence alignment map” format or “SAM”. SAM is the most widely used format for the storing and manipulation of NGS generated nucleotide sequences. As such, the conversion of **.sai** files to **.sam** is essential for use with downstream packages. To convert in the Terminal using BWA^28^, and in the same directory as the **.sai** file type:

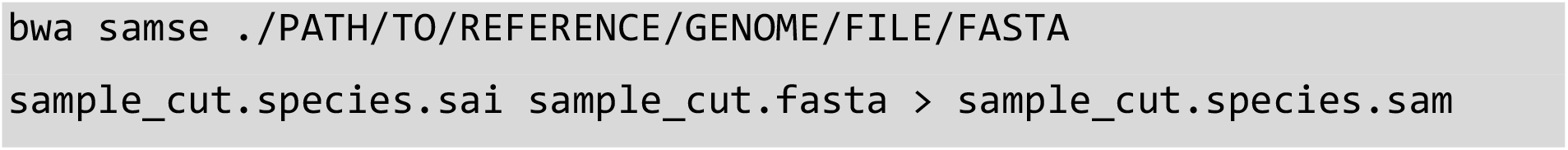

Example:

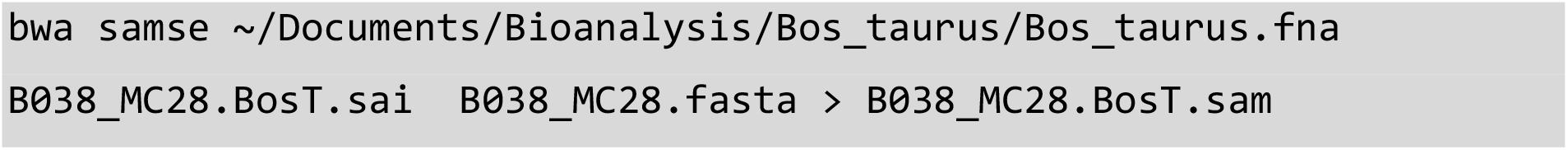

The **samse** command generates alignments given single-end reads. Repetitive hits will be randomly chosen.

#### Step 17. Quality Assurance of NGS Data (3): Confirm Number of Sequences (3)

As previously mentioned, errors can occur with any format conversion and/or file manipulation. As such it is best practice to check data is consistent, throughout the bioinformatic pipeline and reduce possible complications downstream. Unlike previous quality checks, the **view** command of the SAMtools software package is used^29^. In the same Terminal directory as the previous step, type the following:

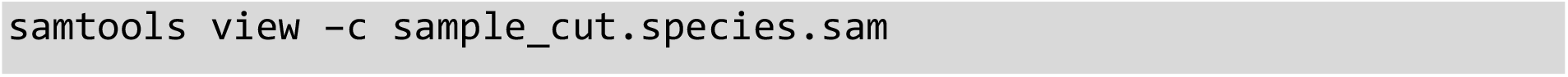

Example:

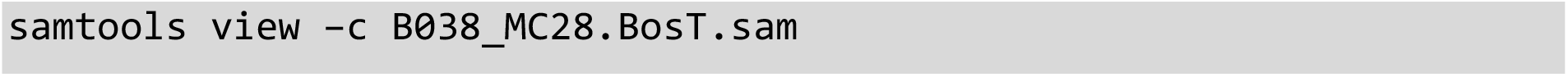

The **-c** option refers to “count”, instructing the Terminal to print only the number of alignments present within a file to **stdout**. Without this parameter all sequences are printed to the Terminal window.

The resulting number generated is representative of the exact amount of sequences within the specified file and should match the total amount of sequences identified in previous quality assurance steps (4, 6).

#### Step 18. Part 1. Clip Nucleotide Overhangs (Picard Tools) (Optional)

A DNA overhang is a portion of unpaired nucleotides at the end of a DNA strand. During the library preparation phase of an aDNA extraction protocol. Enzymes are typically added to remove 3’ overhangs and fill-in 5’ overhanging ends^8,34^. The complimentary fill-in sequence to these authentic overhangs play an important role for the interpretation of aDNA deamination damage patterns^34^.

Occasionally, artificial overhangs occur during the amplification phase using polymerase chain reaction (PCR). These artificial overhangs are typically small and palindromic in nature^35^. In order to ensure artificial PCR overhangs are removed allowing for a more accurate assessment of DNA damage patterns, the soft-clipping of nucleotide overhangs may be performed.

Picard tools are a set of java encoded command-line tools for manipulating NGS data in SAM, BAM, CRAM and VCF formats^36^. The **CleanSam** command performs soft-clipping of nucleotide overhangs beyond the end of reference alignment^36^, thereby removing artificial PCR artefacts. To run, open the Terminal and navigate to the directory containing the aligned SAM file and type:

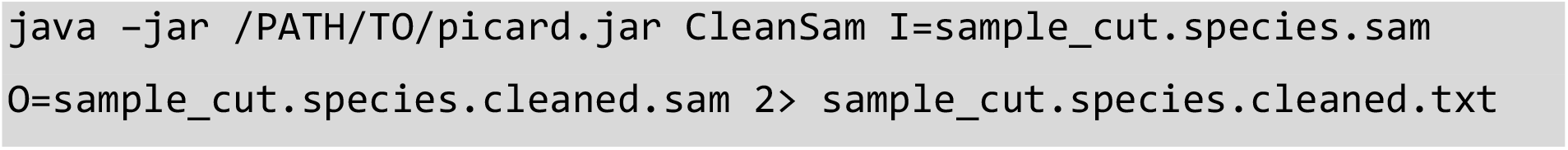

Example:

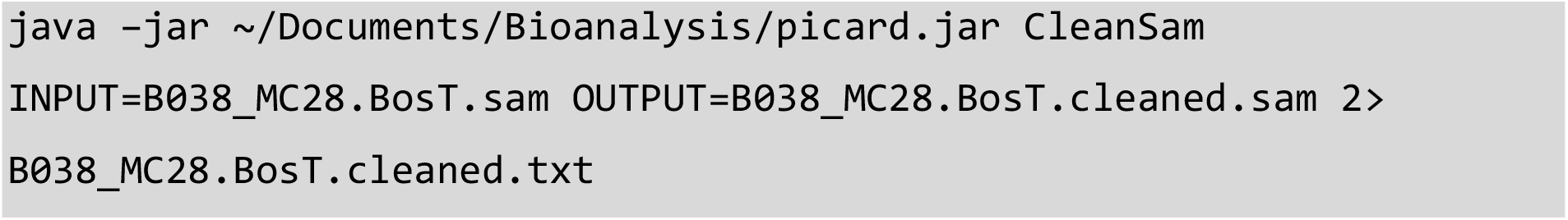

The **I** or **INPUT** parameter refers to the original inputting file.

The **O** or **OUTPUT** parameter refers to the desired outputting file.

Similar to previous steps, the inclusion of the **2>** redirection operator redirects Terminal error output into a specified written file format, which is used as a report. This allows the user to view information regarding which sequences received soft-clipping of overhangs.

We recommend setting **picard** as an environment variable in your **~/.bash_profile**, instead of evoking **java –jar /PATH/TO/picard.jar** each time. This can be achieved by inserting the below line into your bash profile

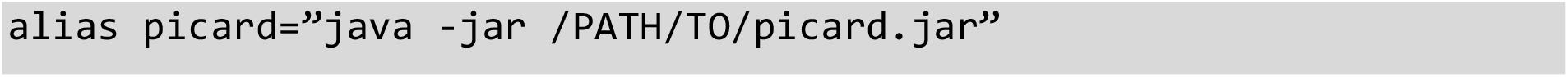

You will need to reference the absolute path to the location of **picard.jar**

In our experience, very few sequences possess nucleotide overhangs. This is most likely a result of the type of enzymes used during the DNA library preparation and amplification phases^8,10^. For this reason, this step is marked as “optional”.

#### Step 18. Part 2. Quality Assurance of NGS Data (4): Confirm Number of Sequences (4) (Optional)

The clipping of DNA overhangs, even if present, should not result in the loss of DNA sequences within a sample file. Occasionally, errors can occur when outputting data into a new file. As such quality assurance steps are needed to ensure the data remains consistent and reduce downstream complications. In the Terminal, navigate to the directory containing the file ending in **.cleaned.sam** and type:

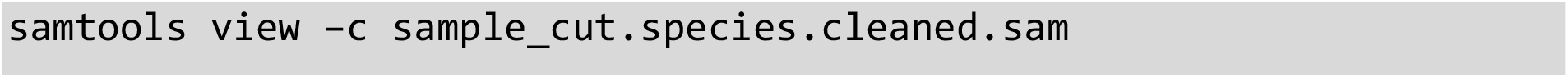

Example:

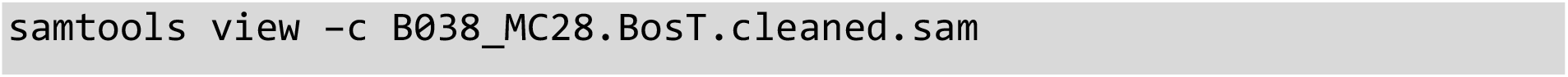

The printed number to **stdout** is representative of the exact amount of sequences within the specified file and should match the total amount of sequences identified in previous quality assurance steps (4, 6, 17).

#### Step 19. Extract Mapped (Aligned) Reads (SAMtools)

Not all of a sample’s sequences will be aligned to a reference genome. For accurate downstream assessment of a taxonomy’s ancient authenticity, those mapped reads must be separated from the remaining non-aligned sequences. This is performed using the SAMtools “view” function^29^. Using the Terminal, in the directory containing the last worked upon file (either ending in **.sam** or **.cleaned.sam**), type the following:

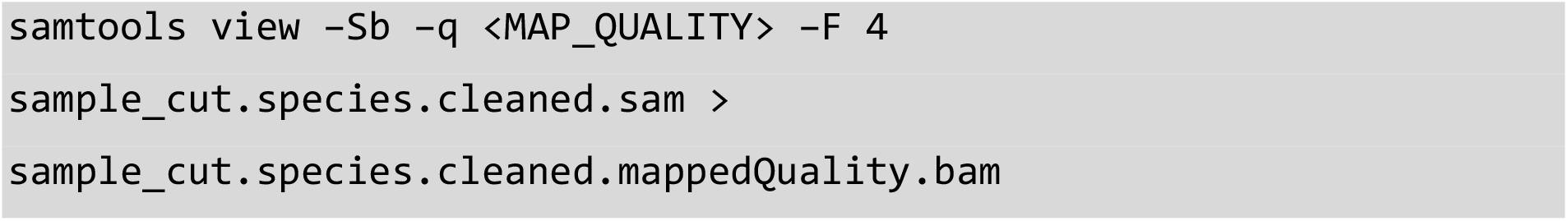

Example:

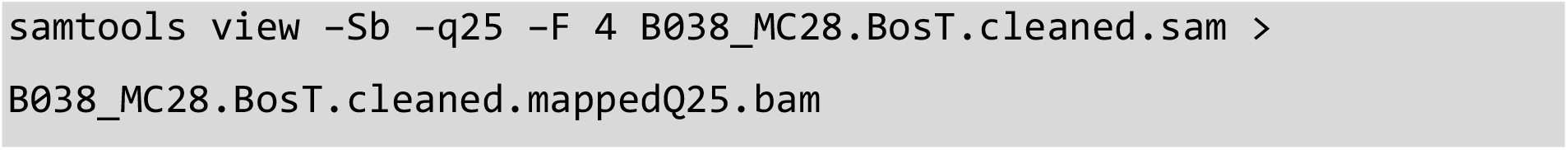

The **-Sb** option refers to the input file as SAM format **S** and the desired output file as BAM format **b**. As previously mentioned, this can also be expresses as **-S -b** in the above command. BAM or “Binary Alignment Map” is the compressed binary representation of SAM^29^. While SAM format is designed to be readable by conventional text-based processing programs, allowing human visualisation of NGS data, the BAM format is designed for quick computational processing, ideal for subsequent processes.

The **-q** option refers to the desired mapping quality of genomic alignments. All alignments with a mapping quality less than the desired threshold are skipped. For aDNA it is standard to use a map quality score between 25 and 30^25,33^. We found that aDNA alignments fell within a map quality score between 25 and 30. Here we use 25.

The **–F** option relates to the “filtering flag”. Reads matching the specified flag are segmented out. For this step we use a flag of “4” (Decimal) or “0×4” (Hexadecimal), resulting in segmentation of unmapped reads from those mapped reads^29^.

The resulting BAM file will be greatly reduced in size and contain only the mapped sample reads to the desired reference genome.

#### Step 20. Quality Assurance of NGS Data (5): Confirm Number of Mapped Reads (1)

As highlighted in previous quality assurance steps, to ensure data consistency throughout the bioinformatic pipeline and to reduce chances of downstream complications, it is critical to implement quality assurance checks. At this stage the user has extracted all mapped reads from a sample, meaning the number of sample sequences within the working file have reduced compared to that identified in previous quality assurance steps (4, 6, 17, 18). As such, it is important to identify the new working number of sequences within the most recent file (**.bam**) and check its consistency with the originating input (**.sam**).

To identify the number of mapped sequences present within the BAM file, the SAMtools **view** command with the **-c** option as described in step 17, is used^29^. To run from the Terminal, navigate to the directory containing the BAM file and type:

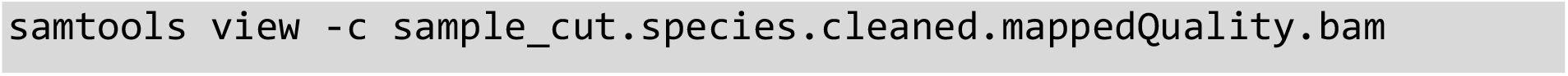

Example:

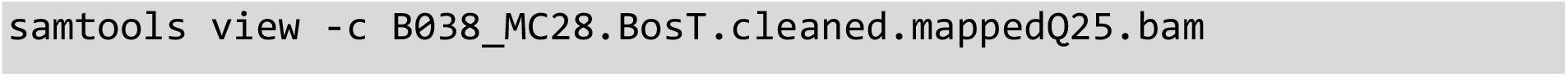

The resulting number represents the exact amount of mapped sequences present within the file.

To test the consistency of the outputted mapped data with that from the originating input file used in step 19, the same SAMtools **view** command is used. However, the lack of a specified output parameter used in conjunction with the **-c** option, instructs the command to print the exact number of mapped sequences within a file to the Terminal window as **stdout**. To run in the Terminal, navigate to the directory containing the SAM file and type:

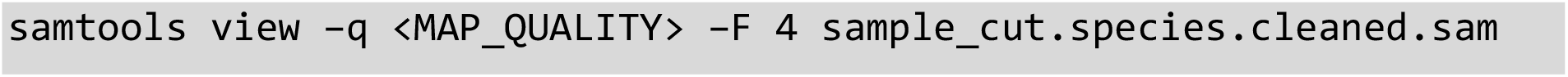

Example:

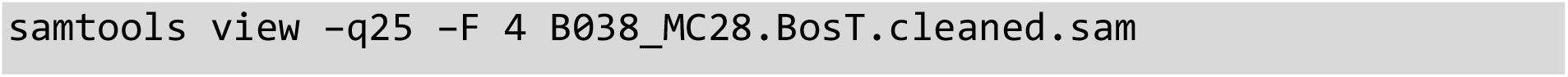

The resulting number should match that identified from the BAM file.

At this point, a minimum threshold of 250 genomic hits are necessary for a taxon to be processed downstream. This is because alignments with less than 250 reads were often found insufficient for **mapDamage** to plot damage patterns effectively.

#### Step 21. Sort Mapped Reads by Leftmost Coordinates (SAMtools)

The sorting of DNA sequences by order of occurrence along a reference genome is required for most downstream applications. This is particularly true for the removal of PCR duplicates (explained in step 23). Typically, sorting is performed using the mapped coordinate position of a sequence against the reference genome.

SAMtools **sort** command uses the leftmost mapped coordinate position of a sequence to accomplish this^29^. To run from the Terminal, navigate to the directory containing the file ending in **.mappedQuality.bam** and type the following:

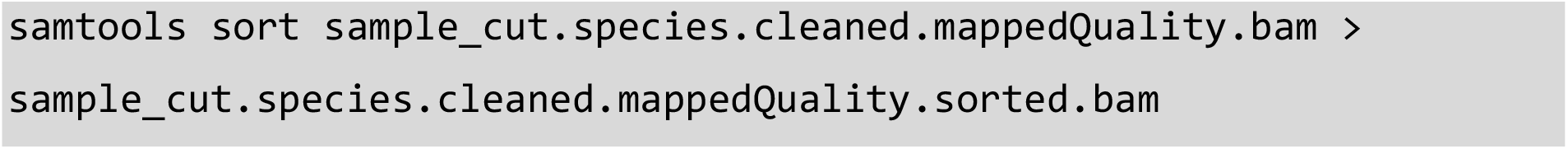

Example:

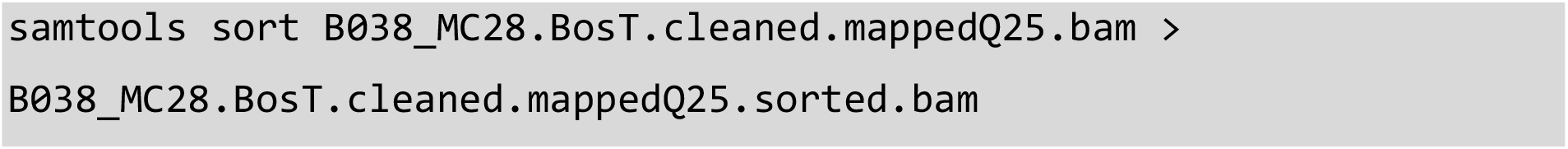

The resulting BAM file will now contain all mapped sequences in order of occurrence along the reference genome.

#### Step 22. Quality Assurance of NGS Data (6): Confirm Number of Mapped Reads (2)

To check that sample data remains consistent throughout the pipeline, with no errors occurring during a bioinformatic process, a file is quality-assessed to reduce the likelihood for downstream complications. As with previous steps, the SAMtool’s **view** command in conjunction with the **-c** option is used to read and print the amount of sequences present within a BAM file^29^. To run in the Terminal, navigate to the directory containing the file ending in **.sorted.bam** and type:

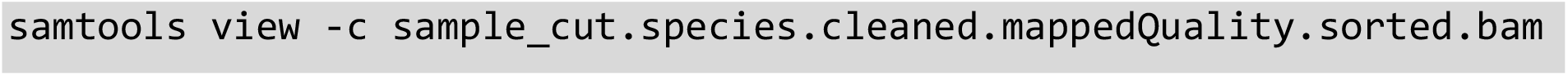

Example:

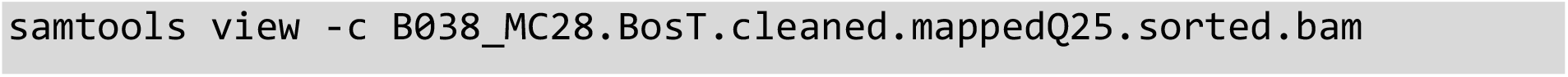

The resulting number should match that identified from the inputting BAM file in step 20.

#### Step 23. Option 1: Remove Duplicate Sequences from Mapped Data using 5’ coordinate position (SAMtools or Picard)

The definition of a PCR duplicate is complicated and subject to much debate within the scientific community. For the purposes of this study we define a duplicate as the presence of two or more identical DNA sequences.

Presence of PCR duplicates can be problematic in the assessment of authentic DNA sequences. The most common reason being the potential for amplification bias, also referred to as base composition bias, introduced during library construction resulting in proportional over representation of specific areas of coding^37^. To ensure the integrity of authentic DNA data, and mitigate the potential effects of duplicate sequences, they are bioinformatically removed.

Method of duplicate removal can vary as can the parameters defining a duplicate sequence. The **rmdup** command of SAMtools identifies PCR duplicates by external coordinate location of outer mapped reads and removes them^29,38^. If two or more reads have the exact same 5’ start position coordinates, the highest map quality score is retained and the others removed. The same can also be accomplished on the reverse 3’ end of a sequence depending on removal option selected. However, it is important to note that the **rmdup** command does not work for unpaired sequences or those mapped to different chromosomes. Meaning a sequence with the same 5’ start coordinate as another sequence but mapped to a different chromosome will be removed as a duplicate.

Picard’s **MarkDuplicates** command likewise uses the 5’ coordinates as a means for duplicate removal, however differs from SAMtool’s **rmdup**, by taking into account the intrachromosomal sequences^36,38^. Additionally, Picard takes into account soft-clipping at the 5’ start position of mapped reads and makes calculations based on where the 5’ start position would be if the entire sequence were mapped to the reference genome ^36,38^. However, the use of external coordinate location as a method for duplicate removal in both methods cannot account for internal sequence variations such as single nucleotide polymorphisms (SNPs), resulting in a potential loss of authentic DNA sequences.

Depending on the size of the originating file, processing speed and memory consumption of the duplicate removal process may be taken into consideration. Previous studies have shown SAMtools as more proficient in this regard using substantially less memory than Picard^38^. For this reason, both options have been detailed below. Where memory is not of concern, we would recommend the use of Picard’s **MarkDuplicates** over SAMtool’s **rmdup**.

To run SAMtool’s **rmdup** from the Terminal, navigate to the directory containing the file ending in **.sorted.bam** and type:

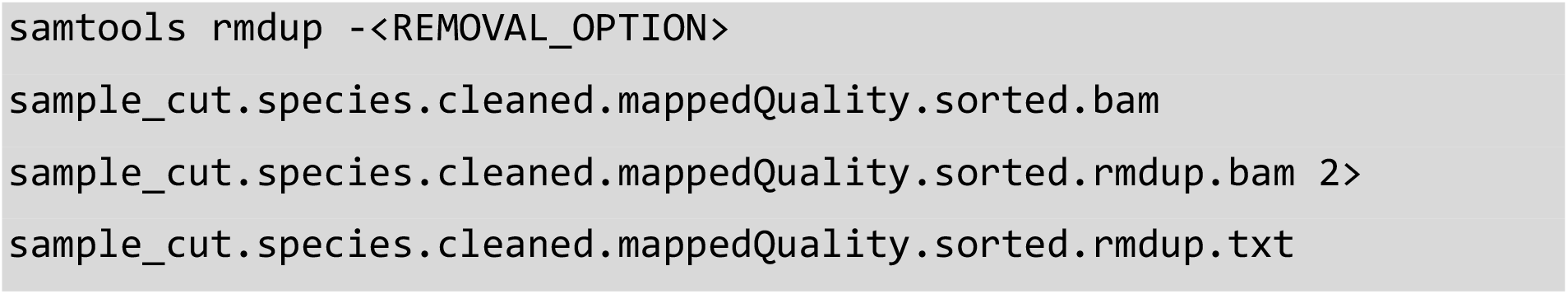

Example:

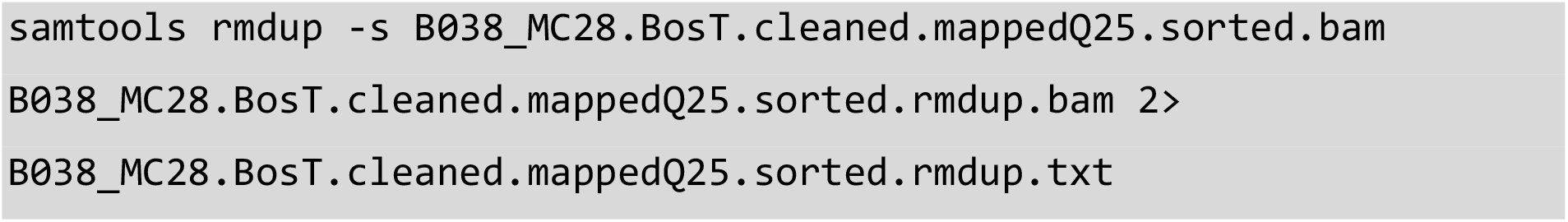

The **<REMOVAL_OPTION>** option allows users to select the type of external coordinate matching for duplicate removal. The **-s** (lower-case s) option removes duplicates for single-end matches at the 5’ location only. The **-S** (upper-case S) option removes duplicates for single and paired-end matches, meaning matches made at the 5’ end of a sequence or at the 3’ end separately will be removed. Sequences have been shown to decline in base quality at the 3’ end of a read^1^, usually resulting in the soft clipping of these bases during BWA alignment. As such, the removal of sequences based solely on a 3’ match is not advised. We would recommend the use of the **-s** (lower-case s) option to reduce the likelihood of authentic DNA removal.

To run Picard’s **MarkDuplicates** from the Terminal, navigate to the directory containing the file ending in **.sorted.bam** and type the following:

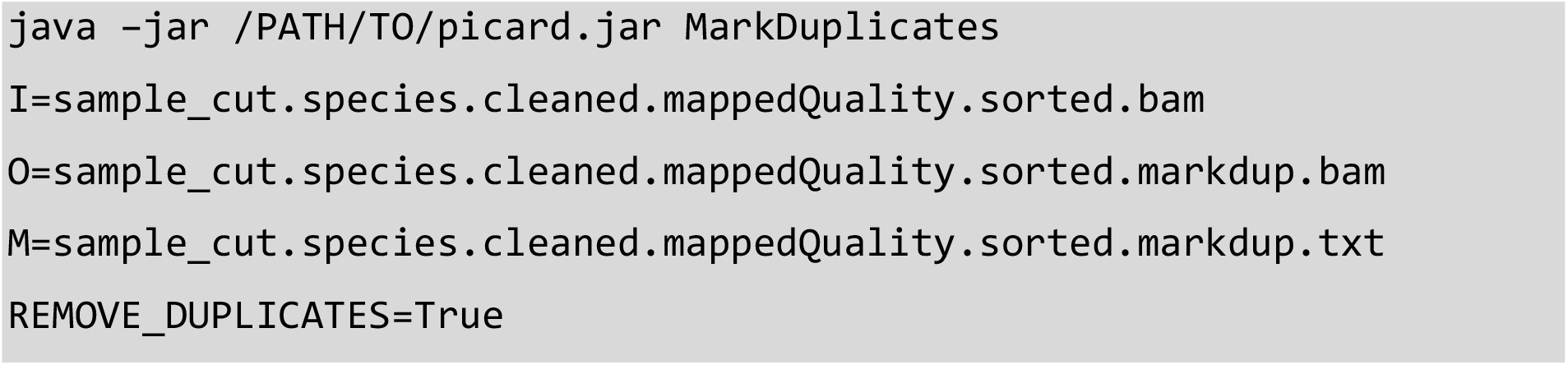

Example:

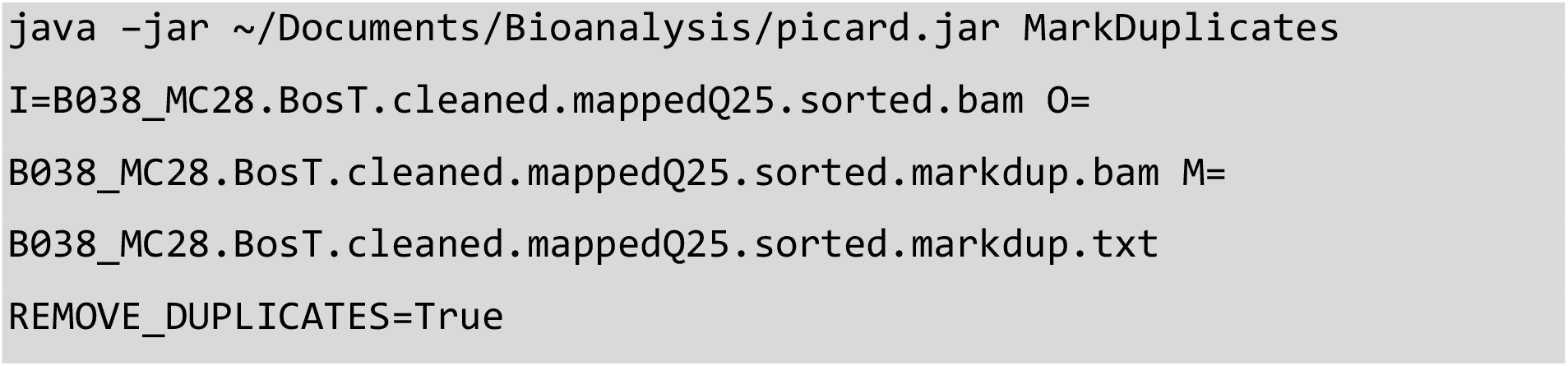

The **M** or **METRICS_FILE** option refers to the text-based log detaining the duplicate sequences removed.

The **REMOVE_DUPLICATES** option used in conjunction with the Boolean **True** instructs the process to remove the duplicate sequences from the outputting file.

Both methods result in a BAM file containing “unique” DNA sequences. The number of duplicate sequences removed are available from the resulting **txt** file.

#### Step 23. Option 2: Remove Duplicate Sequences from Mapped Data (aweSAM)

As mentioned above, both SAMtools and Picard tools identify PCR duplicates by external coordinate location of outer mapped reads at the 5’ potion^29,36^. However, depending on the type of DNA sample sequenced, DNA can share the same outer coordinate location against a reference genome but not represent the same DNA fragment. Instead possessing different internal variations, this is commonly seen in the occurrence of SNPs.

Due to the multi-origin nature of a metagenomic study, we recommend the use of a duplicate removal process that takes into account the coordinate position of a sequence at both the 5’ and 3’ ends as well as accompanying strand information.

aweSAM is a SAM assembly collapser that uses a sequence’s coordinates (5’ and 3’) and strand information as the unique insert identifiers, while keeping the read with the highest mapping quality score^39^. The use of multiple unique insert identifiers allows users to conserve reads that may be lost through other duplicate removal command tools. It should be noted that depending on the size of the input file this process can be time intensive. If time is of concern, we would recommend using one of the duplicate removal functions listed in Option 1.

To run aweSAM, first create a bash shell script of aweSAM_collapser by downloading the script at the link below and ensuring the runtime permissions are updated to allow you to execute the file using **sudo chmod** command. A downloadable aweSAM script can be found here: https://gist.github.com/jakeenk.

Lastly, edit the final line of code within the aweSAM_collapser script to also report the total number of duplicate sequences removed. The script can be edited using the **vim** program, followed by using **i** to go into Insert mode, then pressing **Esc** followed by typing **:wq** to save and quit. This can be performed using the **echo** program to print to **stdout**. The line should now read:

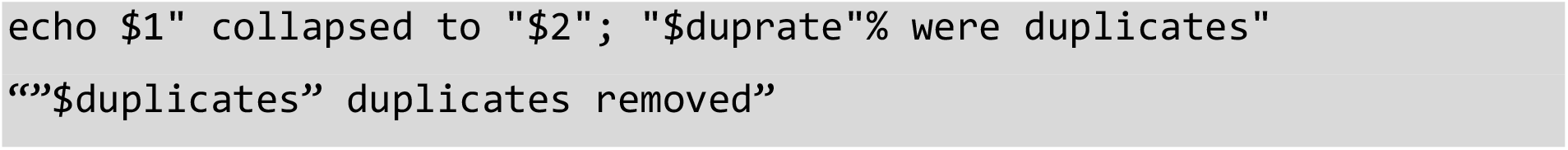

Once created, open the Terminal and navigate to the directory containing the “sorted.bam” file, then type the following:

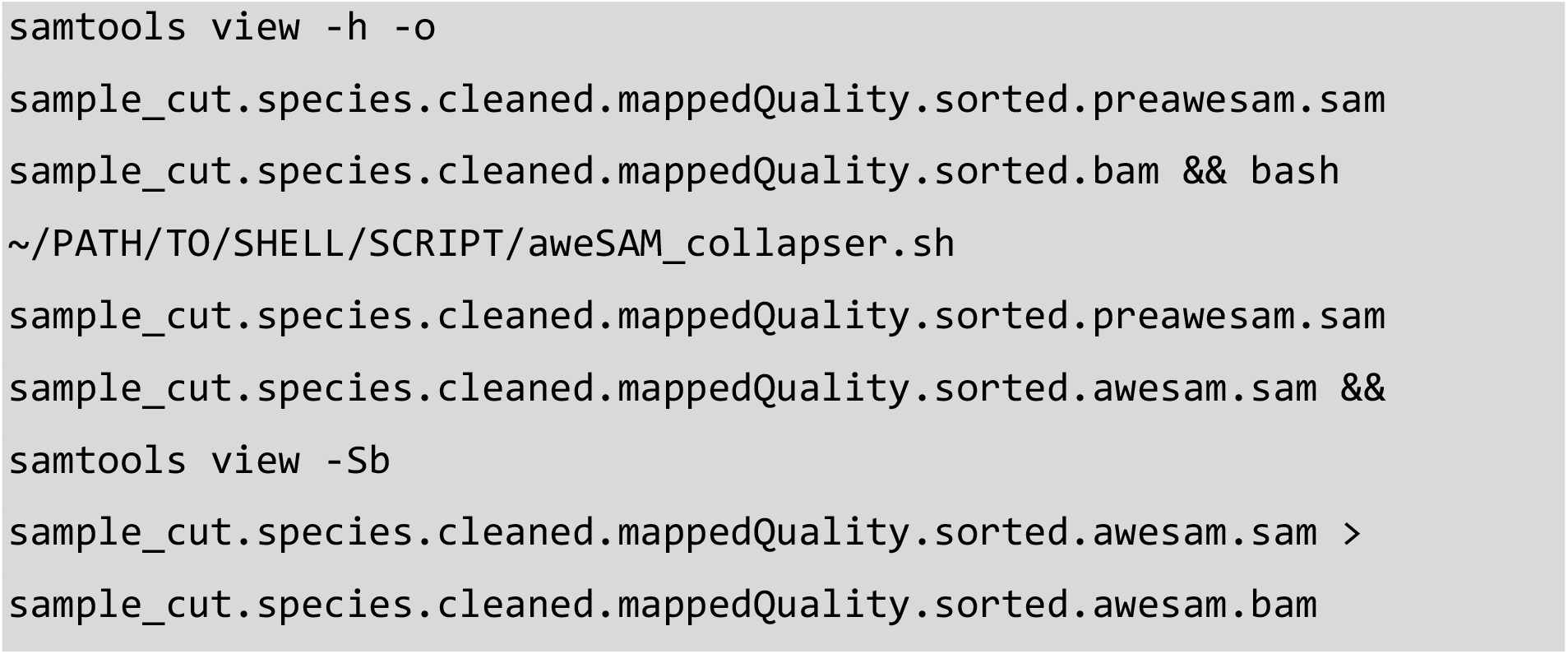

Example:

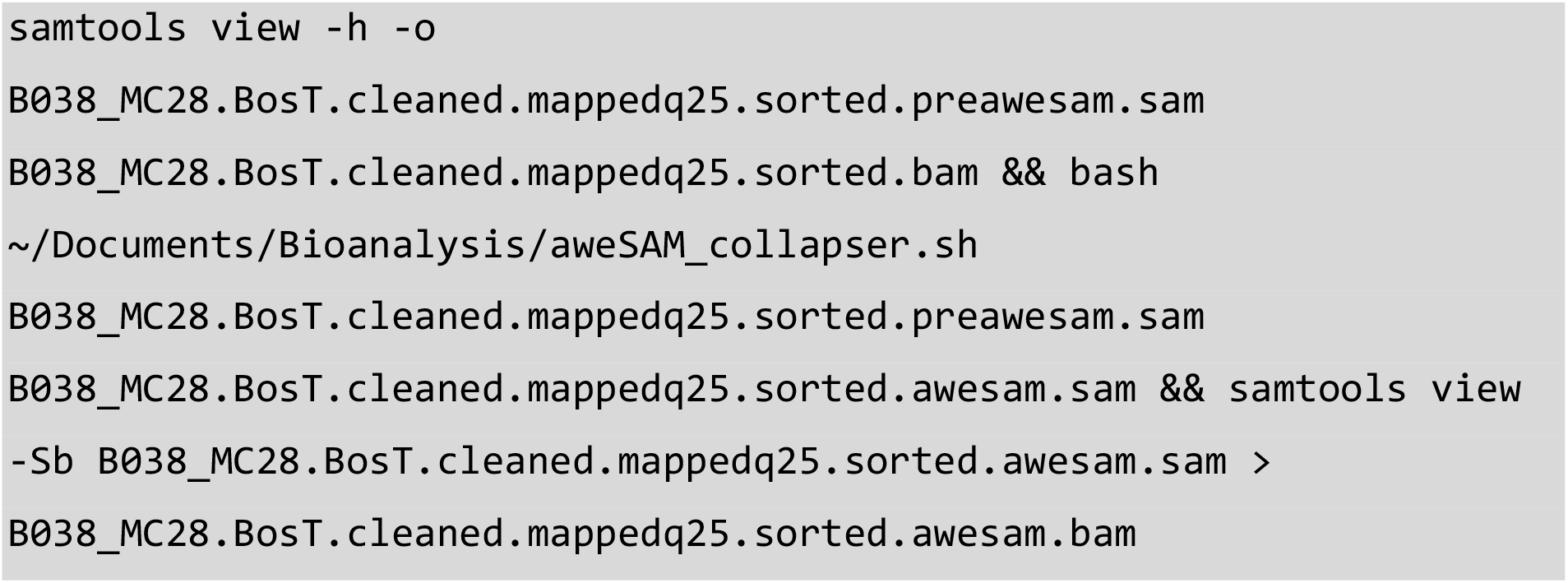

The **-h** option informs the command to include header information in the outputting file.

The **-o** option refers to the desired outputting file.

The use of **bash** here instructs the process to read a series of executable commands contained within the aweSAM_collapser shell script.

The resulting **awesam.bam** file contains unique DNA sequences. The percentage and number of duplicate sequences removed from the originating input file are printed to the Terminal window.

The **samtools** and **bash** programs are all linked together using the **&&** logical operator, which will allow the subsequent program to run if the previous program runs successfully. The output of the last **samtools** is redirected to a BAM file using **>**.

#### Step 24. Quality Assurance of NGS Data (7): Confirm Final Number of Reads

Quality assurance of NGS data is critical for ensuring consistent data flow and reducing the chances of downstream complications. By this step the user has removed all possible PCR duplicates from a mapped DNA sample. This means that the total number of DNA sequences within the most recently outputted file has reduced compared to the amount identified from the input file during the previous quality assurance step (22). As such, the new working number of sequences must be identified and checked for consistency.

First, we utilise the **view** command of SAMtools in conjunction with the **-c** option, as described in step 17, to identify the new number of DNA sequences^29^. Using the Terminal, navigate to the directory containing the duplicates removed BAM file (in this example, the file ending in **.awesam.bam**) and type:

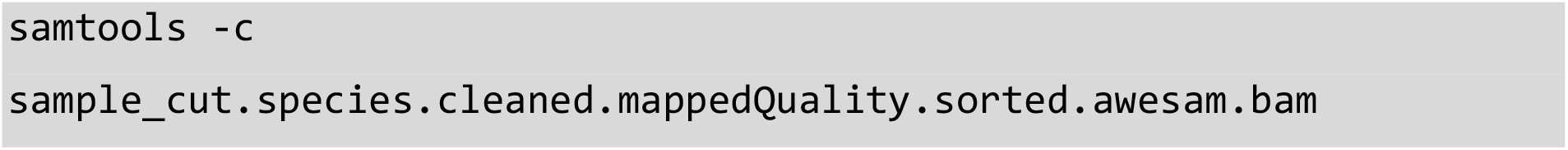

Example:

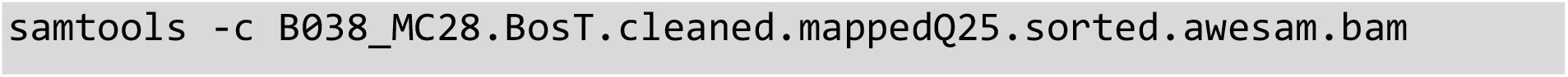

The exact number of unique sample sequences will be printed within the Terminal window.

To check the consistency of data flow. The total number of duplicates removed are added to the remaining number of DNA sequences identified within the newest BAM file. The resulting number should amount to the sequences present within originating input file identified in steps 20 and 22.

#### Step 25. Map Deamination Damage Profile of Mapped Reads (MapDamage 2.0 / Python / R)

MapDamage is a computation framework written in Python and R, that tracks and quantifies aDNA damage patterns among input sequences generated by NGS platforms^40,41^. This is performed using a statistical model based on the damage profile of aDNA fragments described by Briggs^34^. Full details of this framework can be accessed via publication^40,41^.

To run from the Terminal, navigate to directory containing the duplicate removed BAM file (in this example, the file ending in **.awesam.bam**) and type:

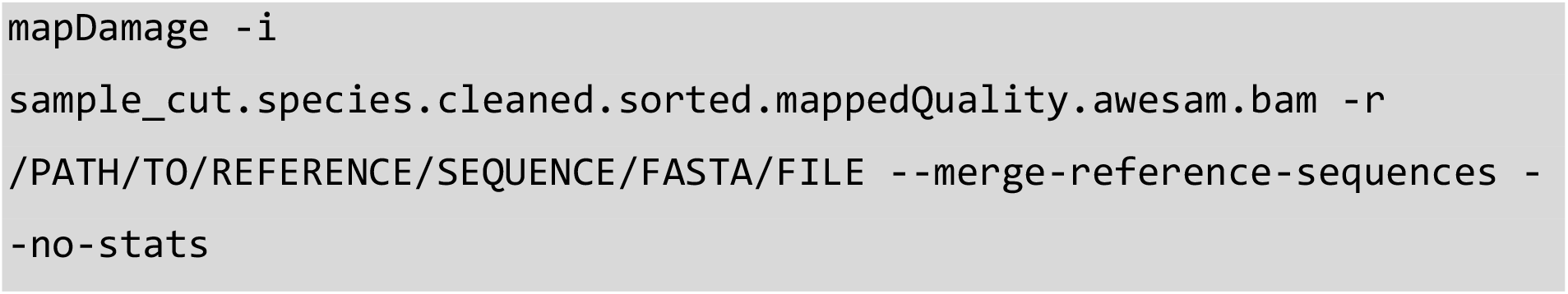

Example:

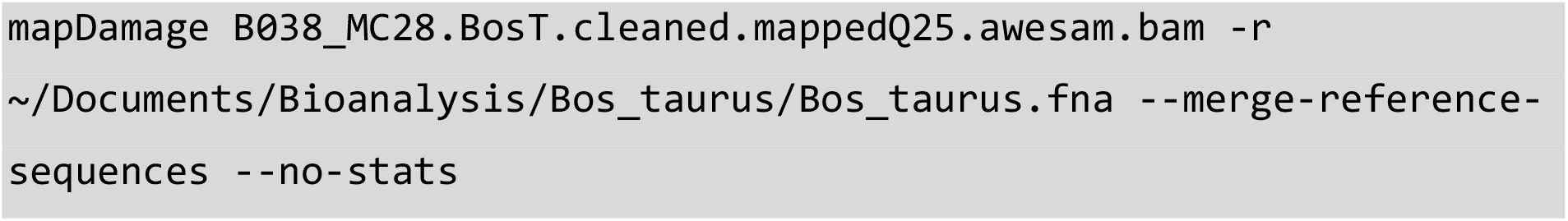

The **-i** option refers to the file containing sample sequences to be tested.

The **-r** option refers to the location of the reference genome to which the sample sequences have been aligned.

The **--merge-reference-sequences** option instructs the program to ignore reference sequence names when tabulating reads. This greatly reduces the amount of memory and disk space used during computation, reducing the likelihood of a code 9 kill error. We would strongly recommend the use of this option for metagenomic studies or those with large sample sizes.

The **--no-stats** option disables the statistical estimation of posterior intervals and distributions. Removal of this option activates statistical estimation by default. The use of this option greatly reduces the processing time required for mapDamage to complete and as such we would recommend the use of this option if statistical estimation is not required. In the case of this study, statistical estimation of posterior intervals and distributions are not required for the confirmation of an ancient taxa.

Completion of the process will result in the creation of a new directory under the same name as the inputting file. Details for each output file can be found disclosed by the authors online at: https://ginolhac.github.io/mapDamage/.

Taxa can be identified as ancient by assessing the frequency of base substitutions located at the Terminal ends of sample sequences. This data can be accessed via the **5pCtoT_freq.txt** file for 5’ C>T substitutions and the **3pGtoA_freq.txt** file for G>A substitutions at the 3’ end. The same data can be graphically visualised by opening the **fragmentation_plot.pdf** file. A threshold of ≥0.05 (representing 5% damage) can be used to identify potentially ancient taxa using an exploratory extraction protocol such as the Collin method^42,43^ (Papers 1, 3). These taxa identifications can be used for the selection of probes for subsequent DNA capture techniques and reassessed using a higher damage threshold. Any taxa reaching a threshold ≥0.10 (10% damage) can be considered definitively ancient in origin.

