## Supplemental User Manual for "An open-sourced bioinformatic pipeline for the processing of Next-Generation Sequencing derived nucleotide reads: Identification and authentication of ancient metagenomic DNA"

```
echo $PATH
```

Where $PATH is a global variable in your Terminal to define where executable commands are located.

In instances when you are installing new programs, you may choose not to include them in your computer's default PATH directories. This allows you to maintain custom programs in a separate directory as well as ensure that default programs are not overwritten by a program of the same name. In order to add a custom directory to your PATH, you can type the below command:

```
export PATH=$PATH:/location/of/your/custom/directory
```

It is important to note however, that the above setting will modify your PATH for that shell session only. Additional shell sessions will not load the custom directory into the PATH. In order to ensure that your custom directory is loaded into the PATH at every shell launch varies depending on the Linux distribution being used. Here, we will use the `.bash_profile` file for the bash shell.

In a new shell window, type the below command to launch `vim`, a command line text editor using the Terminal window.

```
vim ~/.bash_profile
```

Press the `i` character to enter insert mode. Next, add the export line described above to this file. Once inserted, press the `Esc` key followed by typing `:wq!` and then press `Enter`. `Esc` will exit insert mode, the `:` will enter a command, `w` will write to the file being edited and `q!` will exit the vim program.

Now, with each launch of your shell, your custom directory will be loaded into the PATH.

**Step 1. Concatenate Sequencer Output Files**

Upon completion of a NextSeq sequencing run, the system will output four zipped files (fastq.gz) for each sample sequenced. In this step the user will combine these four files into one large zipped file representing all extracted sequences for the given sample. The `cat` program, short for "concatenate" (meaning "to link") allows the Terminal to read multiple files sequentially and output them to the Terminal. To run from the Terminal, navigate to the directory containing the desired NextSeq output files and type:

```
cat sample_file_1.fastq.gz sample_file_2.fastq.gz
sample_file_3.fastq.gz sample_file_4.fastq.gz >
sample_combined.fastq.gz
```

*Example:

```
cat B038_S1_L001_R1_001.fastq.gz B038_S1_L002_R1_001.fastq.gz
B038_S1_L003_R1_001.fastq.gz B038_S1_L004_R1_001.fastq.gz >
B038_combined.fastq.gz
```

*In the example "B038" represents a unique sample code

The redirection command, executed by using `>`, is used to write the Terminal output to a specified file, instead of to the Terminal output, known as standard out or `stdout`. This file will either be created if it doesn't exist or overwritten if it does exist.
Note that within the NextSeq output file names the "L001-4" refers to the lane within the sequencing flowcell in which the data originated. Combining the four files into one reduces the need for repeated processing at subsequent steps.

**Step 2. Quality Control of NGS data (FastQC)**

FastQC[6] is a software package that allows the user to perform quality control checks on raw sequence data coming from NGS platforms. FastQC can be run as a program or directly using the Terminal. To run from the Terminal, navigate to file directory and type the following:

```
fastqc -t <THREADS_AVAILABLE> sample.fastq.gz
```

Example:

```
fastqc -t 12 B038_combined.fastq.gz
```

The `-t` option refers to the amount of processing power or “threads” the user wants to dedicate to the program. Increasing the amount of threads available will increase the processing speed. For a large file, such as those resulting from metagenomic studies, we would recommend using a minimum of 12 threads. The number of threads available to your program would depend on the number of processors and cores each processor has in your computer.

This will result in the generation of two files: a `sample.fastqc.html` containing the quality analysis and a zipped folder (containing the same information).

From the “basic statistics” module shows the total number of sequences, sequences tagged as poor quality, largest sequence length, and %GC can be viewed.

The “per base sequence quality” module displays the average phred quality score (Y axis) of all sequences by base position within a read (X axis). For aDNA sequences, the user should still obtain high phred quality scores for the majority of sequences (preferably above 30). Phred scores are important for understanding the quality of obtained sequences[7]. The higher the phred value the better base cell accuracy is achieved (Table SI 1).

**Table SI 1.** Understanding Phred Quality Scores and Base Call Accuracy. The central column shows the probability of an incorrect base call associated with a phred quality score and the representative accuracy of sequence composition.

| Phred Quality Score | Probability of incorrect base call | Base call accuracy |
|---|---|---|
| 10 | 1 in 10 | 90% |
| 20 | 1 in 100 | 99% |
| 30 | 1 in 1000 | 99.90% |
| 40 | 1 in 10000 | 99.99% |
| 50 | 1 in 100000 | 99.999% |

The "per base sequence content" module displays the base content (Y axis) of all sequences by position in a read (X axis). In a random DNA library, the user would expect little to no difference between the bases of a sequencing run, meaning the lines in the plot should run parallel with each other. It is likely that FastQC will flag this module with a warning due to adapters changing the value of "G" and "C" base content (GC) (Figure SI 1A)[6]. Upon trimming the sample of adapters (step 3), the user can check the sequences again to see little difference between the bases from the $10^{th}$ to $74^{th}$ base pair (10-74bp).

For ancient DNA samples, it is important to note that the first 9 base positions are individual measures that tend to deviate by 10% due to the addition of oligonucleotides at the library preparation phase. The subsequent positions are binned averages, resulting in what should be smooth lines (Figure SI 1B) between 10-74bp. It has been noted that % base composition at the ends of reads that have undergone aggressive adapter trimming (such as that in step 3) are likely to be spurious with sudden deviations in composition[6]. For this reason, it is likely a trimmed sample will flag as failed with the last base position falsely deviating by more that 20%.

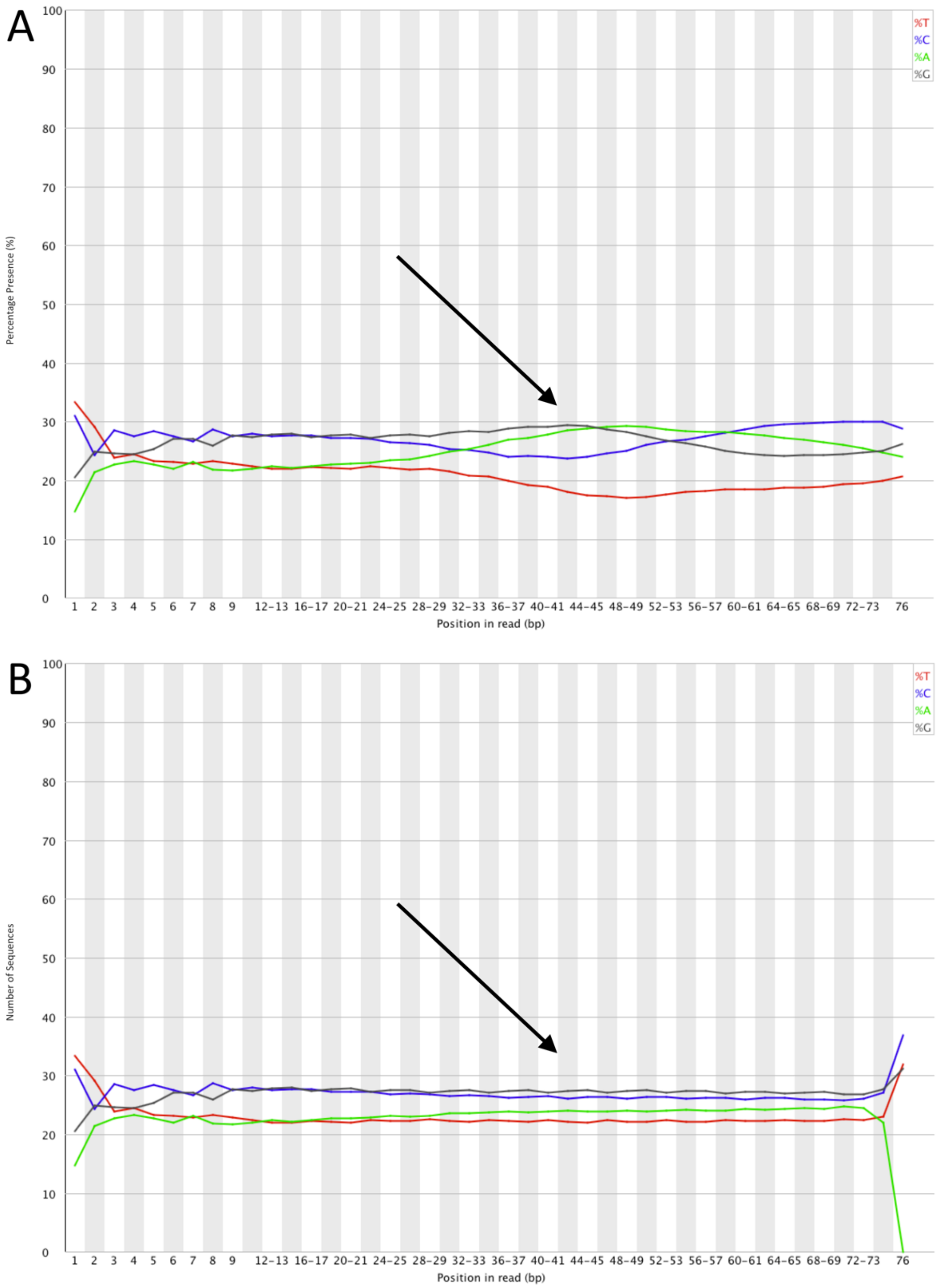


**Figure SI 1.** Sequence Content Across all Bases Using FastQC. (**A**) Prior to adapter trimming. (**B**) Post-trimming of adapters. The arrow highlights an area of difference.

The “per sequence GC content” module measures the mean GC content (X axis) across the length of each sequence (Y axis). Typically, a user would expect to see normal distribution of GC content where a central peak corresponds to the overall GC content of the underlying genome (i.e. 41.6% in the *Homo sapiens* genome if a sample is taken from human bone). While a small shift of an expected GC distribution is indicative of a systematic bias independent of base position, an unusual shaped distribution typically indicates the presence of other underlying genomes[6], or contamination. As the FastQC program does not know the GC content of the underlying genome, the modal GC content is calculated from observed data and used to build a reference distribution. A warning is raised when the sum of deviations from the modal distribution represents more than 15% of reads (Figure SI 2A). Failure is flagged when deviations amount to more than 30% of all reads. By its very nature a metagenomic sample it is expected to have multiple underlying genomes represented and thus a warning or failure of this module is expected. Upon extracting the mapped sequences (step 19) aligned to a desired genome, the user can import into FastQC and compare the mapped %GC to compare to expected GC content. In this case the modal GC content should conform to the desired genomes overall %GC, with a normal peak distribution pattern (Figure SI 2B).

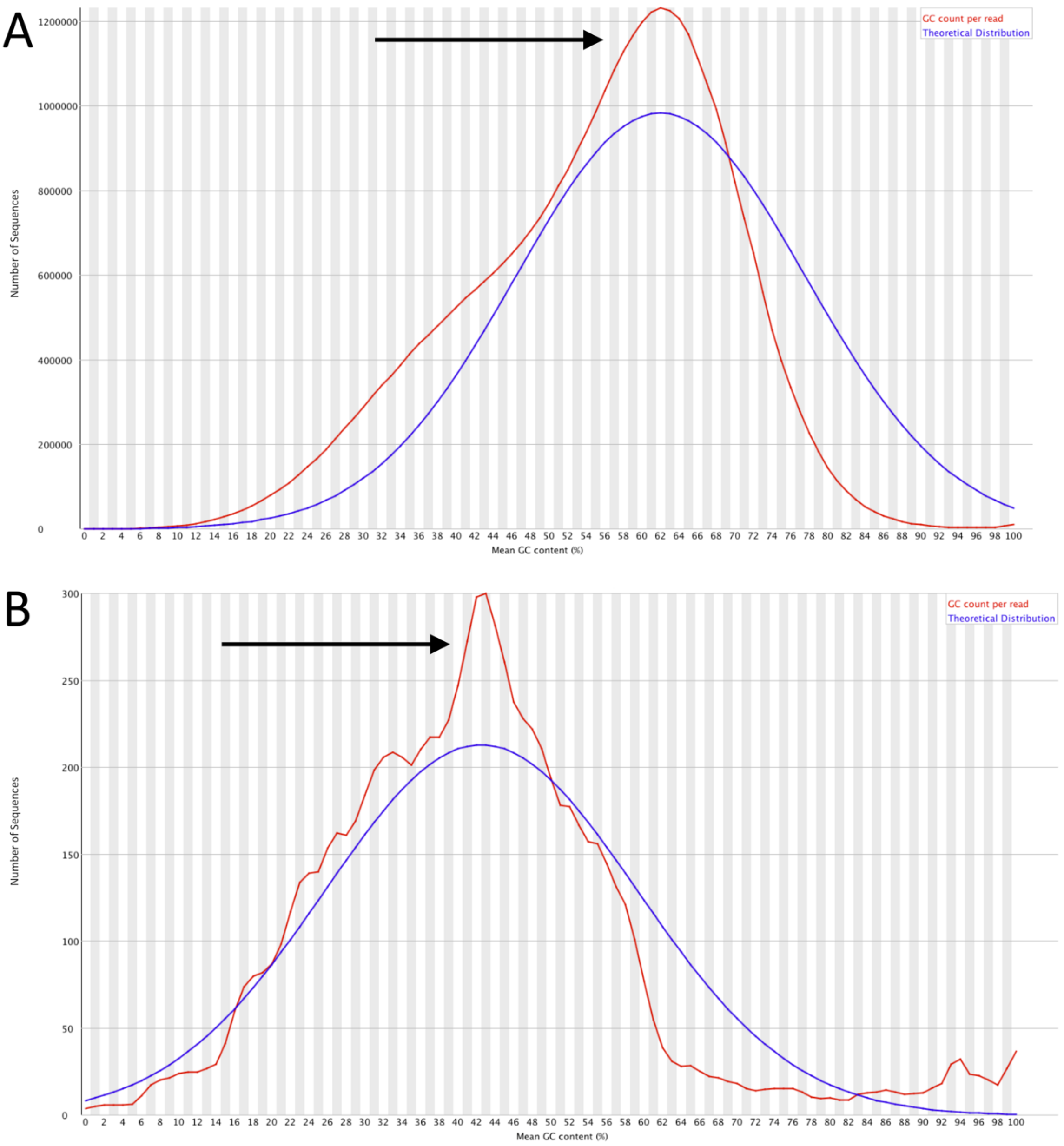


**Figure SI 2.** GC Content Across all Sequences Mapped to a Desired Genome (i.e. *Homo sapiens*) Using FastQC. (**A**) Prior to the extraction of mapped sequences. (**B**) Post extraction of mapped sequences with expected %GC for *Homo sapiens*. The mean percentage of “G” and “C” bases are plotted for all sequences within a sample. The arrow highlights an area of difference.

The “sequence length distribution” module measures the sequence length (X axis) over all sequences (Y axis). If the user performs FastQC prior to the removal of adapters (step 3), the length distribution will always peak at 76bp as the max read length setting on the NextSeq platform using the 500/550 high output V2 (75 cycle) reagent kit (Figure SI 3A). Upon trimming the sample of adapters (step 3) the user can check the sequences again to see a more distributed sequence pattern. Note that the module will flag a warning if all sequences are not the same length[6]. For ancient DNA samples, following the trimming of adapter indexes, the user should expect a distribution of sequences, usually accumulating in a short peak around the 35-50bp mark and a larger peak at the 74-76bp mark (Figure SI 3B).

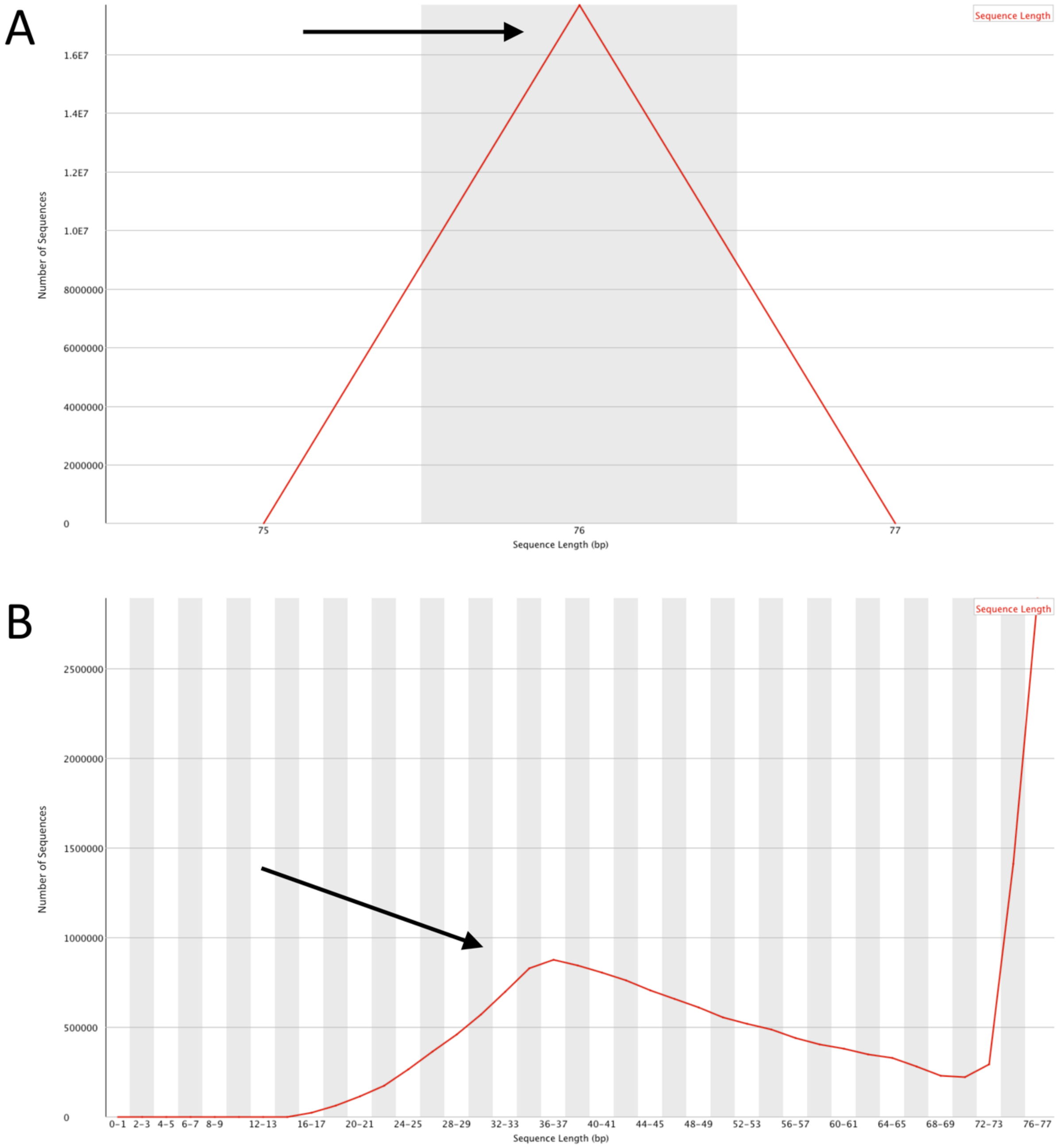

**Figure SI 3.** Distribution of Sequences by Length (bp) Using FastQC. (**A**) Prior to adapter trimming. (**B**) Post-trimming of adapters. All sequences present within a sample sorted according to length of sequence strands. The arrow highlights an area of difference.

The “adapter content” module displays a cumulative percentage of the proportion of the user’s library (Y axis) which has seen adapter sequences at each position (X axis). Once an adapter sequence has been read, it is counted as present through to the end of a read, thereby increasing the percentage of adapter content as read length continues. The user should expect to fail this module (adapter presence on more than 10% of reads) if analysing sequences prior to the trimming of adapters. Post removal of adapter sequences (step 3), the user should expect to pass this module with 0% of all sequences showing evidence for adapters.

**Step 3. Removal of Adapter Sequences (cutadapt)**

Adapter sequences are synthesised oligonucleotides that can be added to the ends of DNA for a variety of applications. In aDNA research, adapters provide two functions, to stabilise authentic damaged sequences and to attach index sequences for sample identification after sequencing[8]. Post sequencing adapters are removed to leave only authentic DNA sequences.

`cutadapt` is a software package that allows users to search all reads within a given sample for a specified adapter sequence and filter reads according to a minimum desired threshold, otherwise known as trimming or a cut[9]. To run from the Terminal, navigate to the directory containing the `combined.fastq.gz` file (from step 1) and type:

```
cutadapt -a <ADAPTER_SEQUENCE_USED> -O 1 -m
<MININMUM_CUT/THRESHOLD_DESIRED> combined.fastq.gz >
sample_cut.fastq 2> sample_cut.txt
```

*Example:

```
cutadapt -a AGATCGGAAGAGCACACGTCTGAACTCCAGTCAC -O 1 -m 28
B038_combined.fastq.gz > B038_MC28.fastq 2> B038_MC28.txt
```

*In the example “MC28” refers to minimum cut (length) used.

Using the Terminal, navigate to the directory containing the trimmed `sample_cut.fastq` file and type the following:

```
wc -l sample_cut.fastq
```

Example:

```
wc -l B038_MC28.fastq
```

Divide the resulting number by four to get the true sequence value. Then, using the `sample_cut.txt` file output in step 3, check the summary section for “Reads written (passing filters)”, this will display the total number of sequences within the trimmed `sample_cut.fastq` file. Both numbers should match.

**Step 5. Convert FASTQ File to FASTA Format**

The conversion of FASTQ files to FASTA format is performed for a number of reasons. First, given that per base sequence quality has been assessed within step 2, the base quality scores are no longer need for downstream processing and as such can be removed to reduce file size, this is automatically performed during conversion into FASTA format. Secondly, the reduction in working file size gives the user the added benefit of faster processing speeds. Lastly, FASTA format has been widely used within bioinformatics for over 30 years, as such many programs used for downstream processing such as BLAST[12] rely on this format. To begin formatting, navigate to the directory containing the `sample_cut.fastq` file and type:

```
cat sample_cut.fastq | awk '{if(NR%4==1)
{printf(">%s\n",substr($0,2));} else if(NR%4==2) print;}' >
sample_cut.fasta
```

Example:

```
cat B038_MC28.fastq | awk '{if(NR%4==1)
{printf(">%s\n",substr($0,2));} else if(NR%4==2) print;}' >
B038_MC28.fasta
```

The `cat` utility program is used to read the contents of the FASTQ file and 'pipes' the `stdout` into the `awk` program as `stdin`. Pipe, represented by the `|` character, is a command line program which allows the output of the 1st program (in this case `cat`) to serve as the input into a 2nd program (in this case `awk`), like a pipeline.

The `awk` program is a tool to write simple yet effective statements in order to manipulate text. It defines text patterns that are to be searched for in each line of the input file. Here the `awk` program reads that if the Number of Records variable (NR), also defined as the line number, when divided by four has a remainder of 1 (denoted as the `%` character in `NR%4==1`) then it will format the line as follows, the `printf` command is invoked and this will print a > character, followed by a format specifier to specify the text as a string (`%s`) followed by a newline character (`\n`) with the string to be decided using the `substr` command. `$0` specifies the whole line to be inputted, and `2` specifies to get the whole line from character 2 onwards (this forms the sequence header), otherwise if the NR variable when divided by four has a remainder of 2, then print the corresponding line (the raw sequence data). Lines 3 and 4 are ignored from the `awk` command. Finally, this manipulated text file is written to a FASTA file using the redirection command `>`.

**Step 6. Quality Assurance of NGS Data (2): Confirm Total Number of Sequences (2)**

As mentioned in step 4, occasionally errors can occur when formatting, manipulating or unzipping a file. It is therefore best practice to check the number of sequences within a file remain consistent with the originating file. To do this we again use the `wc` utility program with the `-l` parameter.

Unlike FASTQ format, FASTA files consist of two lines of code[11]. The first line consists of sequence identifier information or a short description for the sequence, while the second line contains the raw sequence data encoded to IUPAC conventions. As such the resulting total number of lines counted are divided by two to get the true number of sequences present within the FASTA file.

Within the Terminal, make sure the directory contains the `sample_cut.fasta` file, and type:

```
wc -l sample_cut.fasta
```

Example:

```
wc -l B038_MC28.fasta
```

Divide the resulting number by two for the total number of sequences present. The number should match the total number of sequences present in the originating FASTQ file outlined in step 4.

**Step 7. Create a Non-Redundant Sample File (FastX-Toolkit)**

A non-redundant sample refers to one without sequence repetitions, otherwise referred to as duplicates. The more sequences present within a sample file, the longer and less efficient the processing speed of downstream bioinformatic processes. This is a well-documented issue when comparing sample sequences to a genomic database[13,14] (Step 10).

FastX toolkit is a package of command line tools for the pre-processing of FASTQ and FASTA files[15]. The collapser program `fastx_collapser`, merges all duplicate sequences for a region of coding into a single representative sequence, while maintaining read counts. This is performed for all duplicates within a sample until only unique reads remain.

To run the program in the Terminal, navigate to the file containing the `sample_cut.fasta` and type the following:

```
fastx_collapser -v -i sample_cut.fasta -o sample_cut.NR.fasta
```

*Example:

```
fastx_collapser -v -i B038_MC28.fasta –o B038_MC28.NR.fasta
```

*In the example `NR` refers to a non-redundant sample file.

The `-v` option instructs the command to verbose the number of processed reads within the given sample.

The `-i` option refers to the input file, in this case the `sample_cut.fasta`

The `-o` option is the outputting non-redundant file. The resulting output file, should be drastically reduced in size to the inputting FASTA file.

Note, do not remove the original FASTA file as this will be used for in-depth alignments downstream (step 15).

**Step 8. Split the Non-Redundant Sample File into Ten Files of Equal Proportion (pyfasta)**

To further improve the processing speed of comparative analysis (step 10), the non-redundant sample file can be split into ten files of equal proportion. As multiple regions of coding are extracted from origin sources of DNA, the splitting of a non-redundant sample will not affect the identification of a taxon's presence within a sample.

`pyfasta` is package of tools utilising python via a command line interface for the pre-processing of FASTA files[16]. The split command `pyfasta split` allows users to distribute sequencing reads between multiple files via various means.

Here, the `-n` option is used, referring to the number of outputting files desired. Ten files will result in each being 10% representative of the total sequences, while two files represent 50% each.

To run from the Terminal, navigate to the directory containing the non-redundant `sample_cut.NR.fasta` file, and type:

```
pyfasta split -n 10 sample_cut.NR.fasta
```

*Example:

```
pyfasta split -n 10 B038_MC28.NR.fasta
```

*Outputting files will be labelled 00 – 09, this is useful for the following step (step 9).

To validate that resulting files are representative of an entire samples sequences, the mean percentage difference, and standard error between expected hits based on the 10% files and actual hits achieved with the 100% file were calculated. The expected total hits predicted by the 10% file was accurate to the 100% file within –0.007% (±1.101 SEM). Representing a difference of 0.07 hits within 1000.

**Step 9. Randomly Select One of the Non-Redundant Sample Files**

In order eliminate the possibility for selection bias. A randomised number generator is used to determine which of the representative files (step 8) would be used for comparative analysis (step 10).

For this, the `/dev/urandom` number generator available in the majority of Linux distributions used. In order to invoke it, the utility program `grep` is used. `Grep` allows users to search plain text data sets for lines matching a regular expression.

To run using the Terminal, type:

```
grep -m1 -ao '[0-9]' /dev/urandom | head -n1
```

The `-m` option ensures that only 1 line is returned from `/dev/urandom`. `-a` ensures that the line is processed as text and the `o` combination will only print matched, non-empty lines. `-ao` can also be written as `-a -o` in the above program if preferred. A simple regular expression is also specified to provide a single-character number range for `grep` to only return a single-digit number, specified with the `'[0-9]'` regular expression pattern. These three options along with the regular expression, sanitise the output of the `/dev/urandom` program to ensure only single digit number characters are returned. This is then piped into the `head` command. The `head` command, with the `-n1` parameter will ensure that only the first number piped from `grep` is returned.

The process will output a number corresponding to one of the split non-redundant sample files. This will become the users working file for subsequent steps.

**Step 10. Option 1. Compare Sample Sequences to a localised NCBI database (BLAST)**

Using this option, the representative sample file is used to compare its sequences to the entire National Centre for Biotechnology Information's (NCBI) genomic database. Basic Local Alignment Search Tool (BLAST) is a multi-platform algorithm that allows users to query sample sequences against a specified database[12,17,18]. To run from the

Terminal, navigate to the folder containing the randomly selected representative sample file and type:

```
blastn -task exec -query sample_cut.NR.split.fasta -db
PATH/TO/DATABASE/nt -out sample_cut.NR.split.blast.txt -num_threads
<NUMBER_OF_CORES> -word_size <HALF_VALUE_OF_CUT>
```

*Example:

```
blastn -task blastn -query B038_MC28.NR.02.fasta -db ~/ncbi-
blast/db/nt -out B038_MC28.NR.02.blast.txt -num_threads 16 -
word_size 14
```

*The `split` refers to the split file (00-09) randomly selected in step 9. In this example `02` is used.

```
cat * > reference_database.fasta
```

Example:

```
cat * > All_Invertebrates.fasta && rm *.fna
```

The `*` wildcard refers to all files within a specified directory.

The `&&` logical operator is used to chain commands together in one execution, allowing a user to achieve a secondary action after the first action has been completed successfully. In this example, the removal of individual FASTA file after concatenation using the `rm` command, leaving only the full concatenated reference file ("`.fasta`").

To create a *de novo* database, the `makeblastdb` command as part of the BLAST suite is used[19]. The command converts a FASTA file into a reference database. To run from the Terminal, navigate to the directory containing the concatenated reference file and type the following:

```
makeblastdb -in
/PATH/TO/REFERENCE/DIRECTORY/reference_database.fasta -dbtype ‘nucl’
-out /PATH/TO/REFERENCE/DIRECTORY/reference_database
```

Example:

```
makeblastdb -in All_Invertebrates.fasta -dbtype ‘nucl’ -out
~/Documents/Bioanalysis/databases/All_Invertebrates
```

The `-dbtype` option refers to the type of reference sequences used. In this case `nucl` is used, short for **nucl**eotide.

In some situations a user may wish to use a genomic sequence not yet available via the traditional sources of NCBI or the University of California (UCSC)[22,23]. If this is the case, we recommend the user ensure sequences within the desired reference file be assigned unique identifier information to allow for downstream taxonomic assignment. This process should be performed before concatenation. To check this information using the `grep` program type:

```
grep “^>” reference_sequence.fasta
```

Example:

```
grep “^>” bosTau8.fasta
```

The `^>` regular expression in `grep` will print all lines that begin with the `>` character to the FASTA file specified and print it to the Terminal window as `stdout`

In situations where this information is missing. Identifier data can be added using the following in-line perl script:

```
perl -pi -e “s/^>/>Identifier_data-/g” reference_sequence.fasta
```

Example:

```
perl -pi -e “s/^>/>Bos_taurus-/g” bosTau*.fasta
```

The `-p` option causes perl to assume a while loop will be used in the script. In combination with `i`, `-pi` specifies that files processed by the `<>` construct in the while loop are to be edited in-place. The `-e` option is used to specify one line of script entered into the command line. The script entered here is a search and replace operation to be applied to the FASTA file passed in after it. This is done by beginning with the `s/` operator, followed by the regular expression `^>` which will match all lines that begin with the `>` character, this will be replaced with the Identifier data and a `-` symbol (in the above example this is the text “Bos_taurus-”). Finally, the `/g` modifier applies the regular expression globally, i.e. to the whole file.
In the example above the `*` wildcard is used to match all files that begin with the letters “bosTau” and end in “.fasta”

**Step 10. Option 2. Part 2. Compare Sample Sequences to *de novo* Database (BLAST)**

As specified previously, BLAST is a multi-platform algorithm that allows users to query sample sequences against a specified database[12]. In this example a *de novo* database is used. In the Terminal, navigate to the folder containing the randomly selected representative sample file and type:

```
blastn -task option -query sample_cut.NR.split.fasta -db
PATH/TO/DE_NOVO_DATABASE/nt -out sample_cut.NR.split.blast.txt -
num_threads <NUMBER_OF_CORES> -word_size <HALF_VALUE_OF_CUT>
```

Example:

```
blastn -task blastn -query B038_MC28.NR.02.fasta -db
~/Documents/Bioanalysis/databases/All_Invertebrates -out
B038_MC28.NR.02.blast.txt -num_threads 16 -word_size 14
```

Information regarding BLAST options used can be found in Step 10 (Option 1).

**Step 11. Compress BLAST file (Gzip)**

Following a successful BLAST, it is not uncommon that files will be several gigabytes in size. To optimise storage space and reduce processing time of subsequent assignment (Step 12) we would recommend zipping the “txt” file using the `gzip` program. `gzip` is a single-file, lossless data compression tool, that allows users to specify the level and speed of data compression from worst but fastest (`1`) to best but slowest (`9`). To run in the Terminal, navigate to the folder containing the “txt” file and type:

```
gzip -<COMPRESSION_LEVEL> sample_cut.NR.split.blast.txt
```

Example:

```
gzip -9 B038_MC28.NR.02.blast.txt
```

The resulting file will contain the file extension `.gz`

**Step 12. Import BLAST Data into MEGAN**

“MEtaGenome ANalyzer” or “MEGAN”, is a computer program that allows users to import and analyse large datasets of compared genomic sequences (via BLAST or other genomic comparison tools)[24]. During data import, MEGAN assigns a taxon identification to processed read results based on NCBI taxonomy. Using the lowest common ancestor (LCA) algorithm, reads are assigned across a taxonomy (i.e. order, genus, tribe, etc.). The sequences which have a min-score within a specified percentage of the best alignments within a taxonomy are binned into the lowest possible common ancestor position. Sequences aligning to multiple taxa within a grouping are binned into a higher taxonomic level[24] (i.e. sequences assigned to the genera *Bos* and *Capra* will be binned into the family *Bovidae*).

```
gzip –d reference_sequence.fna.gz
```

Example:

```
gzip –d Bos_taurus.fna.gz
```

The `-d` option instructs the tool to "decompress" the file selected.

"Burrows-Wheeler Aligner" or "BWA" is a software package for mapping low-divergent sequences to a large reference genome[28]. Because the software uses low-divergent sequences it is considered more stringent than mass alignment tools such as BLAST. BWA requires a specific FM-index set for alignment to take place. This can be constructed using the `bwa index` command. To run in the Terminal, navigate to the directory containing the blasted `txt.gz` file and type:

```
bwa index reference_sequence.fna
```

Example:

```
bwa index Bos_taurus.fna
```

The `index` command will result in six of the required seven index files for an alignment to take place, including the original `.fna` file, these include: `.fna.amb`, `.fna.ann`, `.fna.bwt`, `.fna.pac`, and `.fna.sa`. All files will have an identical base-name, allowing subsequent tools to recognise an index in its entirety regardless of the file extension.

**Step 14. Part 2. Retrieve Sequence Identifier Information by Creating a FAI File (SAMtools)**

The "FASTA index" or "FAI" file contains all sequence identifier information for a reference genome, including chromosome names, chromosome lengths, offset of the first base of each chromosome sequence, and the length of each FASTA line. This information allows subsequent tools or the user to efficiently query specific regions of a reference genome sequence.

SAMtools is a set of utility commands that primarily allows users to view, read, write, and edit SAM, BAM and CRAM formatted files[29]. Additionally, SAMtools can extract sequence identifier information from FASTQ and FASTA indexed reference files. Using a FASTA reference file, a FAI file can be created using the `faidx` command.

To run from the Terminal, navigate to the directory created in Step 14 (Part 1), containing the reference genome `.fna` file. Type:

```
samtools faidx reference_sequence.fna
```

Example:

```
samtools faidx Bos_taurus.fna
```

The resulting file will contain an identical base-name with a `.fai` file extension and the `.fna` extension, reading as `reference-sequence.fna.fai`.

**Step 15. Compare All Sample Sequences to A Genomic Reference Sequence (BWA)**

As mentioned in the previous step BWA is a software package for mapping low-divergent sequences to a large reference genome[28], and is considered more stringent than methods of mass alignment (Paper 2). In this step, BWA is used to align (also referred to as mapping) sample sequences to a single reference genome using the `aln` command. We recommend the use of `bwa aln` over `bwa-mem` for aDNA sequences (30-70bp), as `bwa-mem` is optimised for sequences >70bp[30], and it has been noted by users that `bwa aln` performs better with reads <70bp[31,32]. In order to perform alignment, the user will need the original FASTA file representing all sample sequences (generated in step 5). Using the Terminal, navigate to the directory containing the original sample FASTA file and type:

```
bwa aln –l <EXCEED_LONGEST_SEQUENCE>
./PATH/TO/REFERENCE/GENOME/FILE/FASTA sample_cut.fasta >
sample_cut.species.sai
```

Example:

```
bwa aln –l 1000 –t 12
~/Documents/Bioanalysis/Bos_taurus/Bos_taurus.fna B038_MC28.fasta >
B038_MC28.BosT.sai
```

The `-l` option refers to “seed length” as part of the seeding process. Seeding can be explained as the finding of exact matches of part of a sample sequence with part of the reference sequence. The larger the seed length required (i.e. 300bp) the faster the alignment process but greater the chance for loss of accuracy. If used correctly a balance may be achievable. In the case of aDNA fragments, seeding is not recommended, owing to base substitutions and its highly fragmented nature. To disable seeding a seed length larger than the longest sample sequence can be specified, thus allowing damaged aDNA sequences to be aligned[33]. We recommend using a seed length of 1000.

The `-t` option refers to the processing power in the form of “threads” that the user wishes to dedicate to the alignment process. The more threads available the faster the alignment process. Here we use 12.

The resulting file with the `.sai` extension is an intermediate file containing “suffix array indexes” for interpretation and conversion into SAM format.

**Step 16. Convert the Aligned Reads to SAM format (BWA)**

Using the `.sai` file generated through alignment in step 15, BWA is further used to convert aligned sequences into “sequence alignment map” format or “SAM”. SAM is the most widely used format for the storing and manipulation of NGS generated nucleotide sequences. As such, the conversion of `.sai` files to `.sam` is essential for use with downstream packages. To convert in the Terminal using BWA[28], and in the same directory as the `.sai` file type:

```
bwa samse ./PATH/TO/REFERENCE/GENOME/FILE/FASTA
sample_cut.species.sai sample_cut.fasta > sample_cut.species.sam
```

Example:

```
bwa samse ~/Documents/Bioanalysis/Bos_taurus/Bos_taurus.fna
B038_MC28.BosT.sai  B038_MC28.fasta > B038_MC28.BosT.sam
```

The `samse` command generates alignments given single-end reads. Repetitive hits will be randomly chosen.

**Step 17. Quality Assurance of NGS Data (3): Confirm Number of Sequences (3)**

As previously mentioned, errors can occur with any format conversion and/or file manipulation. As such it is best practice to check data is consistent, throughout the bioinformatic pipeline and reduce possible complications downstream. Unlike previous quality checks, the `view` command of the SAMtools software package is used[29]. In the same Terminal directory as the previous step, type the following:

```
samtools view –c sample_cut.species.sam
```

Example:

```
samtools view –c B038_MC28.BosT.sam
```

The `-c` option refers to “count”, instructing the Terminal to print only the number of alignments present within a file to `stdout`. Without this parameter all sequences are printed to the Terminal window.

The resulting number generated is representative of the exact amount of sequences within the specified file and should match the total amount of sequences identified in previous quality assurance steps (4, 6).

**Step 18. Part 1. Clip Nucleotide Overhangs (Picard Tools) (Optional)**

A DNA overhang is a portion of unpaired nucleotides at the end of a DNA strand. During the library preparation phase of an aDNA extraction protocol. Enzymes are typically added to remove 3’ overhangs and fill-in 5’ overhanging ends[8,34]. The complimentary fill-in sequence to these authentic overhangs play an important role for the interpretation of aDNA deamination damage patterns[34].

Occasionally, artificial overhangs occur during the amplification phase using polymerase chain reaction (PCR). These artificial overhangs are typically small and palindromic in nature[35]. In order to ensure artificial PCR overhangs are removed allowing for a more accurate assessment of DNA damage patterns, the soft-clipping of nucleotide overhangs may be performed.

Picard tools are a set of java encoded command-line tools for manipulating NGS data in SAM, BAM, CRAM and VCF formats[36]. The `CleanSam` command performs soft-clipping of nucleotide overhangs beyond the end of reference alignment[36], thereby removing artificial PCR artefacts. To run, open the Terminal and navigate to the directory containing the aligned SAM file and type:

```
java -jar /PATH/TO/picard.jar CleanSam I=sample_cut.species.sam
O=sample_cut.species.cleaned.sam 2> sample_cut.species.cleaned.txt
```

Example:

```
java -jar ~/Documents/Bioanalysis/picard.jar CleanSam
INPUT=B038_MC28.BosT.sam OUTPUT=B038_MC28.BosT.cleaned.sam 2>
B038_MC28.BosT.cleaned.txt
```

The `I` or `INPUT` parameter refers to the original inputting file.

The `O` or `OUTPUT` parameter refers to the desired outputting file.

Similar to previous steps, the inclusion of the `2>` redirection operator redirects Terminal error output into a specified written file format, which is used as a report. This allows the user to view information regarding which sequences received soft-clipping of overhangs.

We recommend setting `picard` as an environment variable in your `~/.bash_profile`, instead of evoking `java -jar /PATH/TO/picard.jar` each time. This can be achieved by inserting the below line into your bash profile

```
alias picard=”java -jar /PATH/TO/picard.jar”
```

You will need to reference the absolute path to the location of `picard.jar`

In our experience, very few sequences possess nucleotide overhangs. This is most likely a result of the type of enzymes used during the DNA library preparation and amplification phases[8,10]. For this reason, this step is marked as “optional”.

**Step 18. Part 2. Quality Assurance of NGS Data (4): Confirm Number of Sequences (4) (Optional)**

The clipping of DNA overhangs, even if present, should not result in the loss of DNA sequences within a sample file. Occasionally, errors can occur when outputting data into a new file. As such quality assurance steps are needed to ensure the data remains consistent and reduce downstream complications. In the Terminal, navigate to the directory containing the file ending in `.cleaned.sam` and type:

```
samtools view –c sample_cut.species.cleaned.sam
```

Example:

```
samtools view –c B038_MC28.BosT.cleaned.sam
```

The printed number to `stdout` is representative of the exact amount of sequences within the specified file and should match the total amount of sequences identified in previous quality assurance steps (4, 6, 17).

**Step 19. Extract Mapped (Aligned) Reads (SAMtools)**

Not all of a sample’s sequences will be aligned to a reference genome. For accurate downstream assessment of a taxonomy’s ancient authenticity, those mapped reads must be separated from the remaining non-aligned sequences. This is performed using the SAMtools “view” function[29]. Using the Terminal, in the directory containing the last worked upon file (either ending in `.sam` or `.cleaned.sam`), type the following:

```
samtools view –Sb –q <MAP_QUALITY> –F 4
sample_cut.species.cleaned.sam >
sample_cut.species.cleaned.mappedQuality.bam
```

Example:

```
samtools view –Sb –q25 –F 4 B038_MC28.BosT.cleaned.sam >
B038_MC28.BosT.cleaned.mappedQ25.bam
```

The `-Sb` option refers to the input file as SAM format `S` and the desired output file as BAM format `b`. As previously mentioned, this can also be expresses as `-S -b` in the above command. BAM or "Binary Alignment Map" is the compressed binary representation of SAM[29]. While SAM format is designed to be readable by conventional text-based processing programs, allowing human visualisation of NGS data, the BAM format is designed for quick computational processing, ideal for subsequent processes.

The `-q` option refers to the desired mapping quality of genomic alignments. All alignments with a mapping quality less than the desired threshold are skipped. For aDNA it is standard to use a map quality score between 25 and 30[25,33]. We found that aDNA alignments fell within a map quality score between 25 and 30. Here we use 25.

The `–F` option relates to the "filtering flag". Reads matching the specified flag are segmented out. For this step we use a flag of "4" (Decimal) or "0x4" (Hexadecimal), resulting in segmentation of unmapped reads from those mapped reads[29].

The resulting BAM file will be greatly reduced in size and contain only the mapped sample reads to the desired reference genome.

**Step 20. Quality Assurance of NGS Data (5): Confirm Number of Mapped Reads (1)**

As highlighted in previous quality assurance steps, to ensure data consistency throughout the bioinformatic pipeline and to reduce chances of downstream complications, it is critical to implement quality assurance checks. At this stage the user has extracted all mapped reads from a sample, meaning the number of sample sequences within the working file have reduced compared to that identified in previous quality assurance steps (4, 6, 17, 18). As such, it is important to identify the new working number of sequences within the most recent file (`.bam`) and check its consistency with the originating input (`.sam`).

To identify the number of mapped sequences present within the BAM file, the SAMtools `view` command with the `-c` option as described in step 17, is used[29]. To run from the Terminal, navigate to the directory containing the BAM file and type:

```
samtools view -c sample_cut.species.cleaned.mappedQuality.bam
```

Example:

```
samtools view -c B038_MC28.BosT.cleaned.mappedQ25.bam
```

The resulting number represents the exact amount of mapped sequences present within the file.

To test the consistency of the outputted mapped data with that from the originating input file used in step 19, the same SAMtools `view` command is used. However, the lack of a specified output parameter used in conjunction with the `-c` option, instructs the command to print the exact number of mapped sequences within a file to the Terminal window as `stdout`. To run in the Terminal, navigate to the directory containing the SAM file and type:

```
samtools view –q <MAP_QUALITY> –F 4 sample_cut.species.cleaned.sam
```

Example:

```
samtools view –q25 –F 4 B038_MC28.BosT.cleaned.sam
```

The resulting number should match that identified from the BAM file.

At this point, a minimum threshold of 250 genomic hits are necessary for a taxon to be processed downstream. This is because alignments with less than 250 reads were often found insufficient for `mapDamage` to plot damage patterns effectively.

**Step 21. Sort Mapped Reads by Leftmost Coordinates (SAMtools)**

The sorting of DNA sequences by order of occurrence along a reference genome is required for most downstream applications. This is particularly true for the removal of PCR duplicates (explained in step 23). Typically, sorting is performed using the mapped coordinate position of a sequence against the reference genome.

SAMtools `sort` command uses the leftmost mapped coordinate position of a sequence to accomplish this[29]. To run from the Terminal, navigate to the directory containing the file ending in `.mappedQuality.bam` and type the following:

```
samtools sort sample_cut.species.cleaned.mappedQuality.bam >
sample_cut.species.cleaned.mappedQuality.sorted.bam
```

Example:

```
samtools sort B038_MC28.BosT.cleaned.mappedQ25.bam >
B038_MC28.BosT.cleaned.mappedQ25.sorted.bam
```

The resulting BAM file will now contain all mapped sequences in order of occurrence along the reference genome.

**Step 22. Quality Assurance of NGS Data (6): Confirm Number of Mapped Reads (2)**

To check that sample data remains consistent throughout the pipeline, with no errors occurring during a bioinformatic process, a file is quality-assessed to reduce the likelihood for downstream complications. As with previous steps, the SAMtool's `view` command in conjunction with the `-c` option is used to read and print the amount of sequences present within a BAM file[29]. To run in the Terminal, navigate to the directory containing the file ending in `.sorted.bam` and type:

```
samtools view -c sample_cut.species.cleaned.mappedQuality.sorted.bam
```

Example:

```
samtools view -c B038_MC28.BosT.cleaned.mappedQ25.sorted.bam
```

The resulting number should match that identified from the inputting BAM file in step 20.

To run SAMtool's `rmdup` from the Terminal, navigate to the directory containing the file ending in `.sorted.bam` and type:

```
samtools rmdup -<REMOVAL_OPTION>
sample_cut.species.cleaned.mappedQuality.sorted.bam
sample_cut.species.cleaned.mappedQuality.sorted.rmdup.bam 2>
sample_cut.species.cleaned.mappedQuality.sorted.rmdup.txt
```

Example:

```
samtools rmdup -s B038_MC28.BosT.cleaned.mappedQ25.sorted.bam
B038_MC28.BosT.cleaned.mappedQ25.sorted.rmdup.bam 2>
B038_MC28.BosT.cleaned.mappedQ25.sorted.rmdup.txt
```

The `<REMOVAL_OPTION>` option allows users to select the type of external coordinate matching for duplicate removal. The `-s` (lower-case s) option removes duplicates for single-end matches at the 5' location only. The `-S` (upper-case S) option removes duplicates for single and paired-end matches, meaning matches made at the 5' end of a sequence or at the 3' end separately will be removed. Sequences have been shown to decline in base quality at the 3' end of a read[1], usually resulting in the soft clipping of these bases during BWA alignment. As such, the removal of sequences based solely on a 3' match is not advised. We would recommend the use of the `-s` (lower-case s) option to reduce the likelihood of authentic DNA removal.

To run Picard's `MarkDuplicates` from the Terminal, navigate to the directory containing the file ending in `.sorted.bam` and type the following:

```
java –jar /PATH/TO/picard.jar MarkDuplicates
I=sample_cut.species.cleaned.mappedQuality.sorted.bam
O=sample_cut.species.cleaned.mappedQuality.sorted.markdup.bam
M=sample_cut.species.cleaned.mappedQuality.sorted.markdup.txt
REMOVE_DUPLICATES=True
```

Example:

```
java –jar ~/Documents/Bioanalysis/picard.jar MarkDuplicates
I=B038_MC28.BosT.cleaned.mappedQ25.sorted.bam O=
B038_MC28.BosT.cleaned.mappedQ25.sorted.markdup.bam M=
B038_MC28.BosT.cleaned.mappedQ25.sorted.markdup.txt
REMOVE_DUPLICATES=True
```

```
echo $1" collapsed to "$2"; "$duprate"% were duplicates"
“”$duplicates” duplicates removed”
```

Once created, open the Terminal and navigate to the directory containing the “sorted.bam” file, then type the following:

```
samtools view -h -o
sample_cut.species.cleaned.mappedQuality.sorted.preawesam.sam
sample_cut.species.cleaned.mappedQuality.sorted.bam && bash
~/PATH/TO/SHELL/SCRIPT/aweSAM_collapser.sh
sample_cut.species.cleaned.mappedQuality.sorted.preawesam.sam
sample_cut.species.cleaned.mappedQuality.sorted.awesam.sam &&
samtools view -Sb
sample_cut.species.cleaned.mappedQuality.sorted.awesam.sam >
sample_cut.species.cleaned.mappedQuality.sorted.awesam.bam
```

Example:

```
samtools view -h -o
B038_MC28.BosT.cleaned.mappedq25.sorted.preawesam.sam
B038_MC28.BosT.cleaned.mappedq25.sorted.bam && bash
~/Documents/Bioanalysis/aweSAM_collapser.sh
B038_MC28.BosT.cleaned.mappedq25.sorted.preawesam.sam
B038_MC28.BosT.cleaned.mappedq25.sorted.awesam.sam && samtools view
-Sb B038_MC28.BosT.cleaned.mappedq25.sorted.awesam.sam >
B038_MC28.BosT.cleaned.mappedq25.sorted.awesam.bam
```

The `-h` option informs the command to include header information in the outputting file.

The `-o` option refers to the desired outputting file.

The use of `bash` here instructs the process to read a series of executable commands contained within the aweSAM_collapser shell script.

The resulting `awesam.bam` file contains unique DNA sequences. The percentage and number of duplicate sequences removed from the originating input file are printed to the Terminal window.

The `samtools` and `bash` programs are all linked together using the `&&` logical operator, which will allow the subsequent program to run if the previous program runs successfully. The output of the last `samtools` is redirected to a BAM file using `>`.

**Step 24. Quality Assurance of NGS Data (7): Confirm Final Number of Reads**

Quality assurance of NGS data is critical for ensuring consistent data flow and reducing the chances of downstream complications. By this step the user has removed all possible PCR duplicates from a mapped DNA sample. This means that the total number of DNA sequences within the most recently outputted file has reduced compared to the amount identified from the input file during the previous quality assurance step (22). As such, the new working number of sequences must be identified and checked for consistency.

First, we utilise the `view` command of SAMtools in conjunction with the `-c` option, as described in step 17, to identify the new number of DNA sequences[29]. Using the Terminal, navigate to the directory containing the duplicates removed BAM file (in this example, the file ending in `.awesam.bam`) and type:

```
samtools -c
sample_cut.species.cleaned.mappedQuality.sorted.awesam.bam
```

Example:

```
samtools -c B038_MC28.BosT.cleaned.mappedQ25.sorted.awesam.bam
```

The exact number of unique sample sequences will be printed within the Terminal window.

To check the consistency of data flow. The total number of duplicates removed are added to the remaining number of DNA sequences identified within the newest BAM file. The resulting number should amount to the sequences present within originating input file identified in steps 20 and 22.

**Step 25. Map Deamination Damage Profile of Mapped Reads (MapDamage 2.0 / Python / R)**

MapDamage is a computation framework written in Python and R, that tracks and quantifies aDNA damage patterns among input sequences generated by NGS platforms[40,41]. This is performed using a statistical model based on the damage profile of aDNA fragments described by Briggs[34]. Full details of this framework can be accessed via publication[40,41].

To run from the Terminal, navigate to directory containing the duplicate removed BAM file (in this example, the file ending in `.awesam.bam`) and type:

```
mapDamage -i
sample_cut.species.cleaned.sorted.mappedQuality.awesam.bam -r
/PATH/TO/REFERENCE/SEQUENCE/FASTA/FILE --merge-reference-sequences -
-no-stats
```

Example:

```
mapDamage B038_MC28.BosT.cleaned.mappedQ25.awesam.bam -r
~/Documents/Bioanalysis/Bos_taurus/Bos_taurus.fna --merge-reference-
sequences --no-stats
```

The `-i` option refers to the file containing sample sequences to be tested.
